# A short-lived peptide signal regulates cell-to-cell communication in *Listeria monocytogenes*

**DOI:** 10.1101/2023.07.04.547616

**Authors:** Benjamin S. Bejder, Fabrizio Monda, Bengt H. Gless, Martin S. Bojer, Hanne Ingmer, Christian A. Olsen

## Abstract

Quorum sensing (QS) is a mechanism that regulates group behavior in bacteria, and in Gram-positive bacteria, the communication molecules are often cyclic peptides, called autoinducing peptides (AIPs). We recently showed that pentameric thiolactone-containing AIPs from *Listeria monocytogenes*, and from other species, spontaneously undergo rapid rearrangement to homodetic cyclopeptides, which hampers our ability to study the activity of these short-lived compounds. Here, we developed chemically modified analogues that closely mimic the native AIPs while remaining structurally intact, by introducing N-methylation or thioester-to-thioether substitutions. The stablilized AIP analogues exhibit strong QS agonism in *L. monocytogenes* and allow structure–activity relationships to be studied. Our data provide evidence that loudly suggest that the most potent AIP is in fact the very short-lived thiolactone-containing pentamer. Further, we find that the QS system in *L. monocytogenes* is more promiscuous with respect to the structural diversity allowed for agonistic AIPs than reported for the more extensively studied QS systems in *Staphylococcus aureus* and *Staphylococcus epidermidis*. The developed compounds will be important for uncovering the biology of *L. monocytogenes*, and the design principles should be broadly applicable to the study of AIPs in other species.

## Introduction

*Listeria monocytogenes* is a notorious foodborne pathogenic Gram-positive bacterium, causing intracellular infections in humans and animals. It is found in soil and wastewater, and can persist in food-processing facilities, even after sanitization. When ingested, *L. monocytogenes* can cause disease ranging from gastroenteritis to life-threatening conditions in immunocompromised, pregnant, or elderly individuals. Being able to survive a wide range of environmental conditions requires tightly regulated gene expression^1,2^. This is in part accomplished by quorum sensing (QS)^3-9^ that in *L. monocytogenes* affects protein secretion, cell invasion, biofilm formation, and virulence gene expression. QS constitutes regulatory systems for sensing population density, through excretion and recognition of autoinducer molecules, causing changes in population behavior in different bacteria^10^. In Gram-positive bacteria, the best characterized QS system is that encoded by the <u>a</u>ccessory <u>g</u>ene regulator (*agr*) locus in *Staphylococcus aureus*^11^. In this system, the signaling molecules are cyclic, autoinducing peptides (AIPs), that commonly contain a thiolactone formed between the C-terminal carboxylate and the thiol side chain of a cysteine residue in the *i*–5 position from the C-terminus, together with a linear N-terminal exotail of varying length^12-16^. The *agr* locus encodes the AIP precursor peptide (AgrD) and a membrane-bound peptidase (AgrB) needed for the closing of the thiolactone ring. Additionally, it encodes a two-component system, consisting of the receptor-histidine kinase (AgrC) and response regulator (AgrA) that together comprise the AIP-sensing and signal transduction components^17^. The *agr* system of *L. monocytogenes*, like that of *S. aureus*, contains the four genes, *agrBDCA*, that are autoregulated and expressed from the same promoter^3,4^. The system has been known for 20 years and has been shown to affect the expression of hundreds of genes^6,18^, but its regulatory function is not yet well understood.

The identity of the *L. monocytogenes* AIP is debated. Previously, a hexameric AIP (**P1**) with a single amino acid exotail has been identified using LC-MS/MS^19^, and Zetzmann *et al*. provided evidence to suggest a pentameric AIP (**P2**) without an exotail (Figure 1)^8^. Previously, we failed to identify **P1** by using a thiolactone-trapping methodology^16,20^ and showed that AIPs without an exotail spontaneously undergo S→N acyl shift to give homodetic peptides, like **P3** (Fig. 1)^20^. Further, we were able to identify **P3** in supernatant from *L. monocytogenes* bacterial culture and showed that it can induce QS in a bioluminescent reporter strain of *L. monocytogenes*; albeit, at high concentrations (EC_50_ ∼8 μM) compared to those needed for autoinduction in Staphylococci^16^. Concurrently, a similar S→N acyl shift phenomenon was described in the bacterium *Ruminiclostridium cellulolyticum*^21^. Although, our work demonstrated the rapid rearrangement of **P2** to give **P3** at pH 7, the high concentration needed for **P3** to activate QS still led us to speculate whether the **P2** thiolactone could play a role in the activation of QS in *L. monocytogenes*.

**Figure 1.**
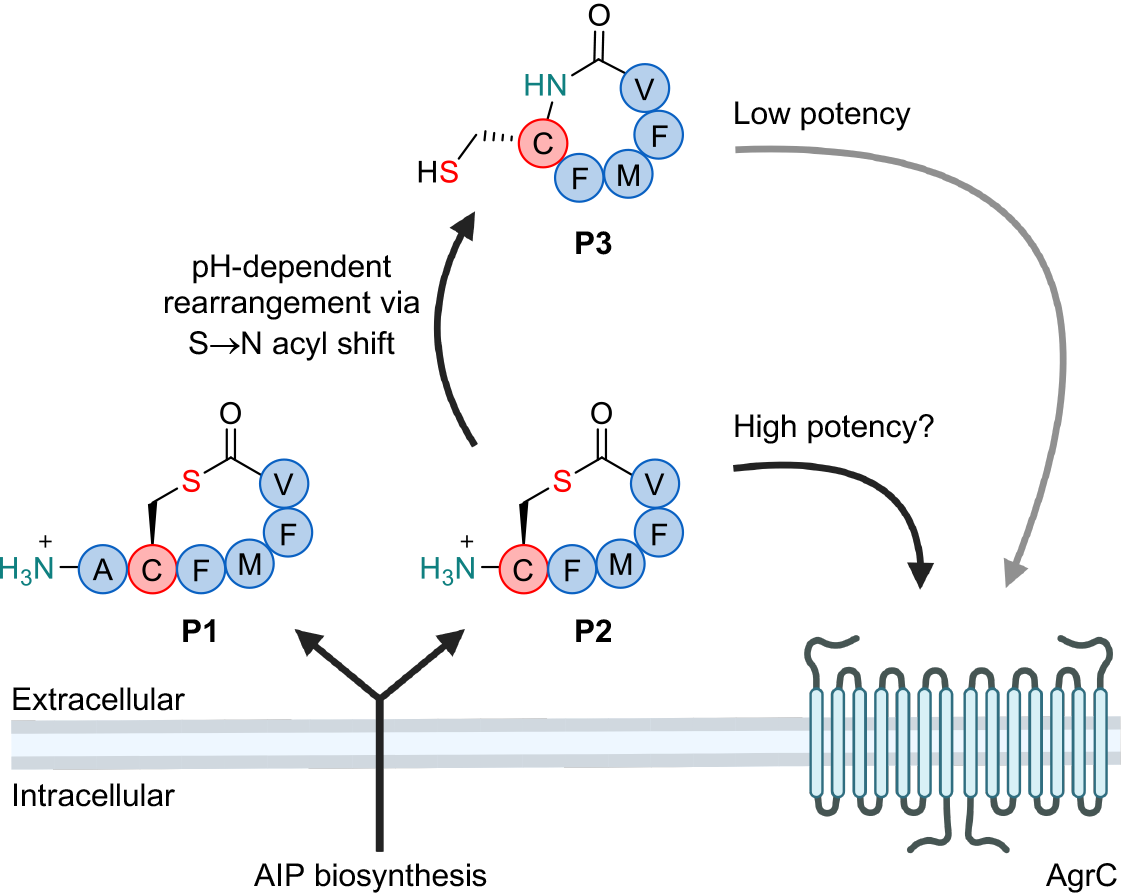
Cartoon representation of the *agr* system and proposed autoinducing peptides in *Listeria monocytogenes*. Part of the figure was prepared using BioRender.

Here, we report the development of stabilized analogues, mimicking the structure of **P2**, to investigate whether **P2** could act as short-lived but highly potent activator of the *L. monocytogenes* QS system.

## Results

### Assay optimization and evaluation of compounds P1–P3

First, we assessed the rate of rearrangement from **P2** to **P3** at 37 °C, which is the relevant temperature when the bacteria infect humans, and here **P2** rearranged with a half-life of 1.3 min at pH 7 (Fig. 2a). For *S. aureus*, the AIP concentration in cultures at early stationary phase have been determined to be ∼1 μM^22^, and if assuming a similar concentration of **P2**, the remaining concentration of thiolactone-containing peptide after 9 and 13 min would be <10 nM and <1 nM, respectively (Fig. 2b). Albeit it would be difficult to imagine building up a concentration as high as 1 μM for **P2**, because of its rapid, continuous degradation. Nevertheless, this exercise suggests that if **P2** were act as signaling molecule in a cell density sensing system, it would require rapid receptor activation at low peptide concentrations.

**Figure 2.**
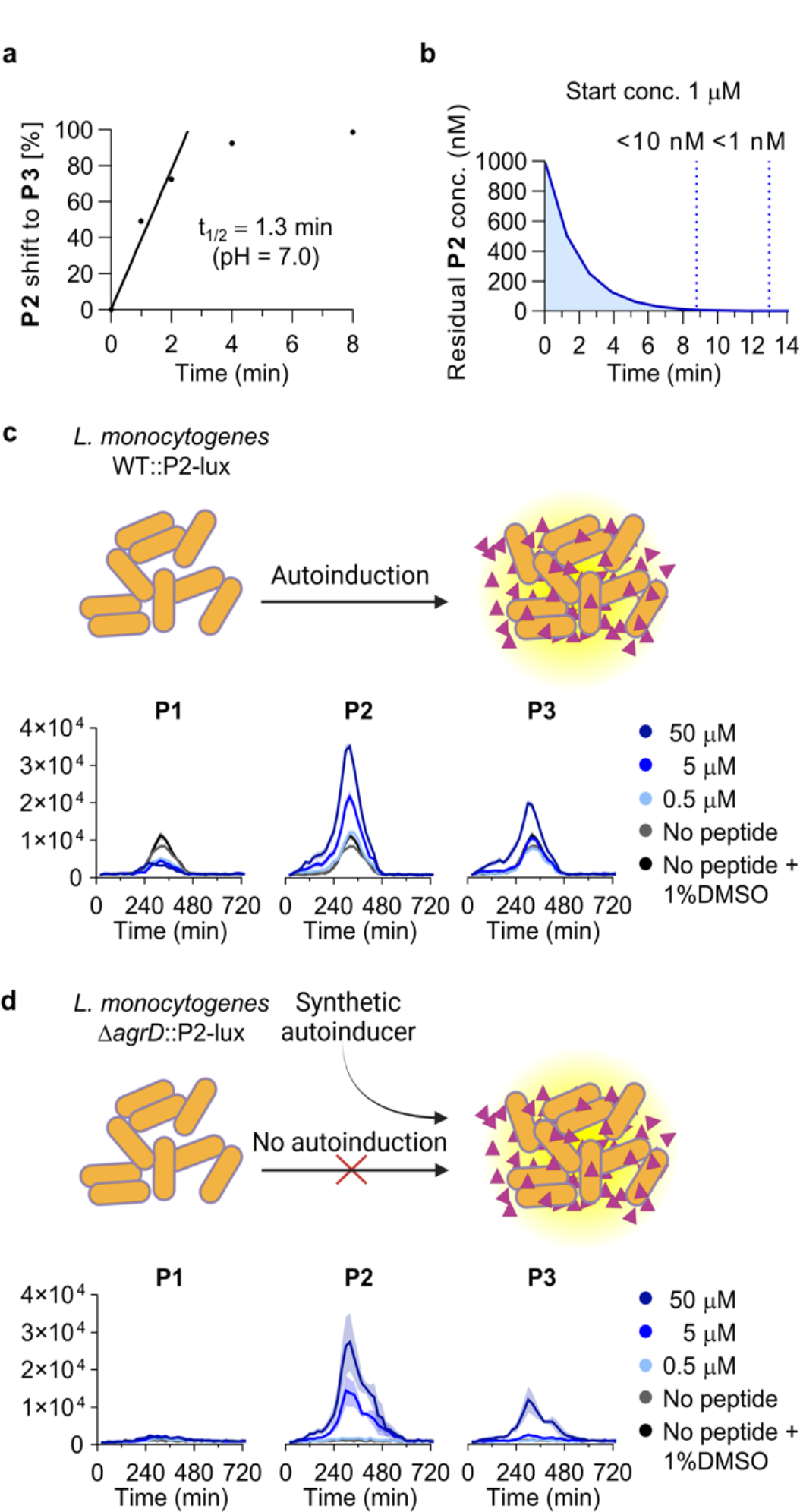
Rearrangement of P2 and activity of P1–P3. **a**, UPLC-based assessment of the rearrangement of **P2** to give **P3. b**, Plot of the decaying concentration of **P2** from a starting concentration of 10 μM (based on the determined t_1/2_ = 1.3 min). **c**, Effect of selected concentrations of **P1**–**P3** on the *agr* activity in luminescence-based reporter strain WT::P2-lux of *L. monocytogenes* grown at 37 °C in tryptic soy broth (TSB) medium. **d**, Effect of selected concentrations of **P1**–**P3** on the *agr* activity in luminescence-based reporter strain Δ*agrD*::P2-lux of *L. monocytogenes* grown at 37 °C in tryptic soy broth (TSB) medium. Shadings on the graphs represent the standard error of the mean (SEM). All data are based on at least three individual assays performed in at least technical duplicate. Part of the figure was prepared using BioRender.

To start to address this question, we compared the effects of **P1**–**P3** on *agr*-dependent bacterial reporter strains at 37 °C, using *L. monocytogenes* EGDe WT and a Δ*agrD* mutant strain, both carrying a chromosomal integration of the *agr* promoter fused to the *lux* operon, enabling a bioluminescent readout as measure of *agr* activity (referred to as WT::P2-lux and Δ*agrD*::P2-lux, respectively)^8^. To improve signal-to-noise we first optimized the assay conditions inspired by a study on luciferase-based reporter strains of *Lactococcus lactis*, in which the luciferase substrate flavin mononucleotide was added^23^. We found that addition of flavin mononucleotide (10 mg/L, ∼26 μM) amplified maximal luminescence readout of the WT::P2-lux reporter strain by ∼80% and total area under the curve by ∼90% when grown in tryptic soy broth (TSB), without affecting the growth (Supplementary Fig. 1). We then tested synthetic peptides **P1**–**P3** against the WT::P2-lux and Δ*agrD*::P2-lux reporter strains in TSB and brain heart infusion (BHI) medium at 37 °C. Importantly, **P2** was added directly from DMSO stock solution, to preserve the integrity of the thiolactone motif for as long as possible during the experiment (Fig. 2c,d and Supplementary Figs. 2–5). The trends previously observed for **P1** and **P3** at 30 °C were recapitulated under the optimized conditions. Treating WT::P2-lux with **P1** resulted in inhibition of the signal in both BHI and TSB media, while giving rise to a slight induction in the Δ*agrD*::P2-lux reporter strain after 240 min, which was most pronounced in BHI medium (Fig. 2c and Supplementary Figs. 2–5). The homodetic peptide **P3** did not cause inhibition of WT::P2-lux and produced early induction of *agr* in both TSB and BHI media (Fig. 2c, Supplementary Figs. 2 and 4). More interestingly, **P2** was able to increase *agr* activity 3-fold at 50 μM and 2-fold at 5 μM for WT::P2-lux in TSB, compared to the untreated WT::P2-lux (Fig. 2c, Supplementary Figs. 2 and 4).

A less pronounced increase in peak intensity was also observed for **P2** at 5 μM in BHI (Supplementary Fig. 4) and early induction of *agr* was observed in both TSB and BHI (Fig. 2c,d and Supplementary Figs. 2 and 4). For the activation of Δ*agrD*::P2-lux by **P2**, we observed full restoring of WT levels in BHI and surpassing the WT level in TSB, even at 5 μM compound concentration (Fig. 2d and Supplementary Figs. 3 and 5; see the Supplementary Information page 8 for further discussion). Even though **P2** is inherently unstable at near-neutral pH, its ability to activate *agr* at lower concentrations than **P1** and **P3** supports its function as a bona fide AIP of *L. monocytogenes*, but its potency remains elusive.

### Development of N-terminally modified mimics of P2

Previously, we synthesized an *N*-acetylated version of **P2**, to obtain a stable thiolactone mimic of **P2**, but this peptide showed no activation of Δ*agrD*::P2-lux and inhibited the *agr* activity of WT::P2-lux^20^, which we confirmed with the optimized assay protocol (Supplementary Fig. 6). Since the N-acetylated mimic cannot be protonated, we speculated if a positively charged amino group at the N-terminus, as present in **P2**, could be important for activity. We therefore designed a stabilized mimic of the compound by introducing two methyl groups instead of acetyl (**1**; Fig. 3a and Supplementary Fig. 7). The two methyl groups prevented S→N acyl shift as intended, and we observed >90% purity of compound **1** after incubation for 22 hrs in phosphate buffer (pH 7.0) at 37 °C (Fig. 3b and Supplementary Fig. 8).

**Figure 3.**
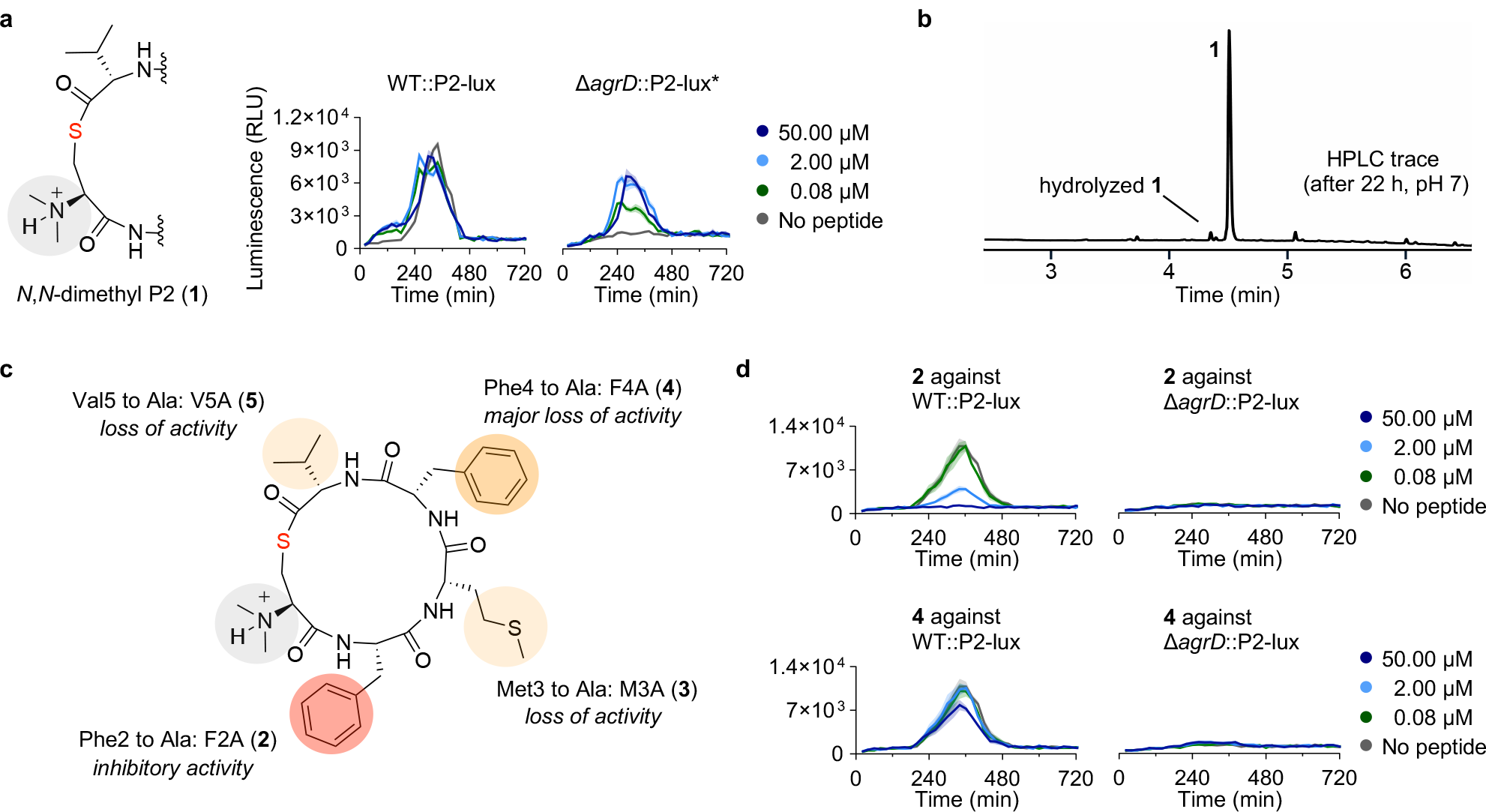
Chemical structures of peptide analogues 1–5, rearrangement evaluation of 1, and selected reporter strain assay data. **a**, Structural modification to give **1** and effect of selected concentrations of **1** on the *agr* activity in luminescence-based reporter strains of *L. monocytogenes* grown at 37 °C in tryptic soy broth (TSB) medium. **b**, HPLC analysis of the stability of **1** after incubation for 22 h in buffer. **c**, Overview of the effects of alanine substitution of compound **1. d**, Effects of selected concentrations of **2** and **4** on the *agr* activity in luminescence-based reporter strains of *L. monocytogenes* grown at 37 °C in tryptic soy broth (TSB) medium.

Remarkably, the thiolactone **1** caused early activation of *agr* in the WT::P2-lux strain and significant induction of *agr* in Δ*agrD*::P2-lux at just 80 nM compound concentration, displaying superior potency to any of the previously tested peptides (Fig. 3a, Table 1, Supplementary Fig. 9). However, thiolactone **1** was unable to increase *agr* activity above the WT levels, unlike the effect observed with **P2** and WT::P2-lux. Similarly, against Δ*agrD*::P2-lux we observed an upper limit of activation (∼6500 RLU) with this **P2** analogue, which was lower than the maximum luminescence of the WT::P2-lux strain (Fig. 3a and Supplementary Fig. 10). Nevertheless, the potency of **1** supports the hypothesis that *N*-acetylation is inappropriate for studying the biological activity of exotail-free thiolactone-containing AIPs. Dimethylation of the N-terminus therefore provides a novel architecture to achieve stabilization of these AIPs (Fig. 3b).

**Table 1.** Activation of *agr* in *L. monocytogenes*.^a^.

| compound | %-activation [2 μM] | %-activation [80 nM] |
| --- | --- | --- |
| **P1** | 11 ± 6<sup>b</sup> | inactive<sup>c,d</sup> |
| **P2** | **160 ± 51<sup>b</sup>** | inactive<sup>c,d</sup> |
| **P3** | 15 ± 6<sup>b</sup> | inactive<sup>c,d</sup> |
| **1** | 63 ± 5 | 35 ± 3 |
| **6** | 27 ± 3 | inactive<sup>c</sup> |
| **7** | **210 ± 10** | 38 ± 2 |
| **8** | **270 ± 10** | 43 ± 8 |
| **9** | 80 ± 5 | 74 ± 8 |
| **10** | **350 ± 12** | **274 ± 7** |

<sup>a</sup>Values are extracted from assays with Δ*agrD*::P2-lux and normalized to average maximum luminescence of untreated Δ*agrD*::P2-lux (0%) and average maximum luminescence of untreated WT::P2-lux (100%).
<sup>b</sup>Tested at 5 μM. <sup>c</sup>Inactive defined as less than 10% normalized activation. <sup>d</sup>Tested at 500 nM.

Having a potent **P2** mimic in hand, we performed an alanine scan based on **1** (peptides **2**–**5**; Fig. 3c), to study the effect of each position in **P2** on *agr* activity. We observed the most pronounced effect when substituting either phenylalanine (**2** and **4**), which resulted in a complete loss of activation of the Δ*agrD*::P2-lux reporter (Fig. 3d and Supplementary Figs. 9 and 10). Interestingly, compound **2** was able to fully inhibit *agr* activity in the WT::P2-lux strain at 2 and 50 μM concentration (Fig. 3d). In contrast to **2** and **4**, the alanine analogues **3** and **5** were partial agonists unable to reach the same maximum activation of Δ*agrD*::P2-lux as **1**. Compounds **3** and **5** also displayed inhibition of peak luminescence in the WT::P2-lux, with analog **5** being able to inhibit the activity by 20% at just 80 nM concentration (Supplementary Figs. 9 and 10). Although all alanine substitutions affected the agonistic effect, only modest inhibitors were discovered.

### Development of alternative stabilized analogues of P2

To explore alternative ways of stabilizing the AIP structures, we synthesized lactam analogues of **1** and **P2** (**6** and **7**, respectively; Fig. 4a,b and Supplementary Figs. 11 and 12). Based on structure-activity relationship (SAR) studies performed on AIPs from *S. aureus*, we expected the lactam substitution to cause a decrease in activity^24-28^. However, the lactam analog **7** proved to be an effective agonist that increased *agr* activity of WT::P2-lux above controls at concentrations down to 80 nM (Fig. 4b, Table 1, and Supplementary Fig. 13). At the highest concentration tested for **7** (50 μM), the culture reached same maximal level of *agr* activity in WT::P2-lux and Δ*agrD*::P2-lux, around 2.5-fold higher than the untreated WT control (Fig. 4b and Supplementary Fig. 13), which was slightly lower than the maximal activation achieved with the native, short-lived compound **P2**. With compound **6** we observed an upper maximal *agr* activation in Δ*agrD*::P2-lux similar to the effect observed for **1** and, contrary to compound **7**, this analog did not induce *agr* at 80 nM concentration (Fig. 4a, Table 1, and Supplementary Fig. 13). We further synthesized a lactone analog of **P2** (**8**; Figure 4c and Supplementary Fig. 14), which showed a similar degree of induction and potency as the lactam **7**, reaching roughly the same maximal luminescence levels in Δ*agrD*::P2-lux and half-maximal activation at 400 nM (Supplementary Fig. 15). However, the lactone also proved inherently unstable, which causes an underestimation of its activity, as discussed for compound **P2**. Thus, after 22 hrs at 37 °C and pH 7.0, 36% of **8** was hydrolyzed and 48% had undergone rearrangement through O→N acyl shift, to form its corresponding homodetic peptide (Figure 4d and Supplementary Fig. 15a,b).

**Figure 4.**
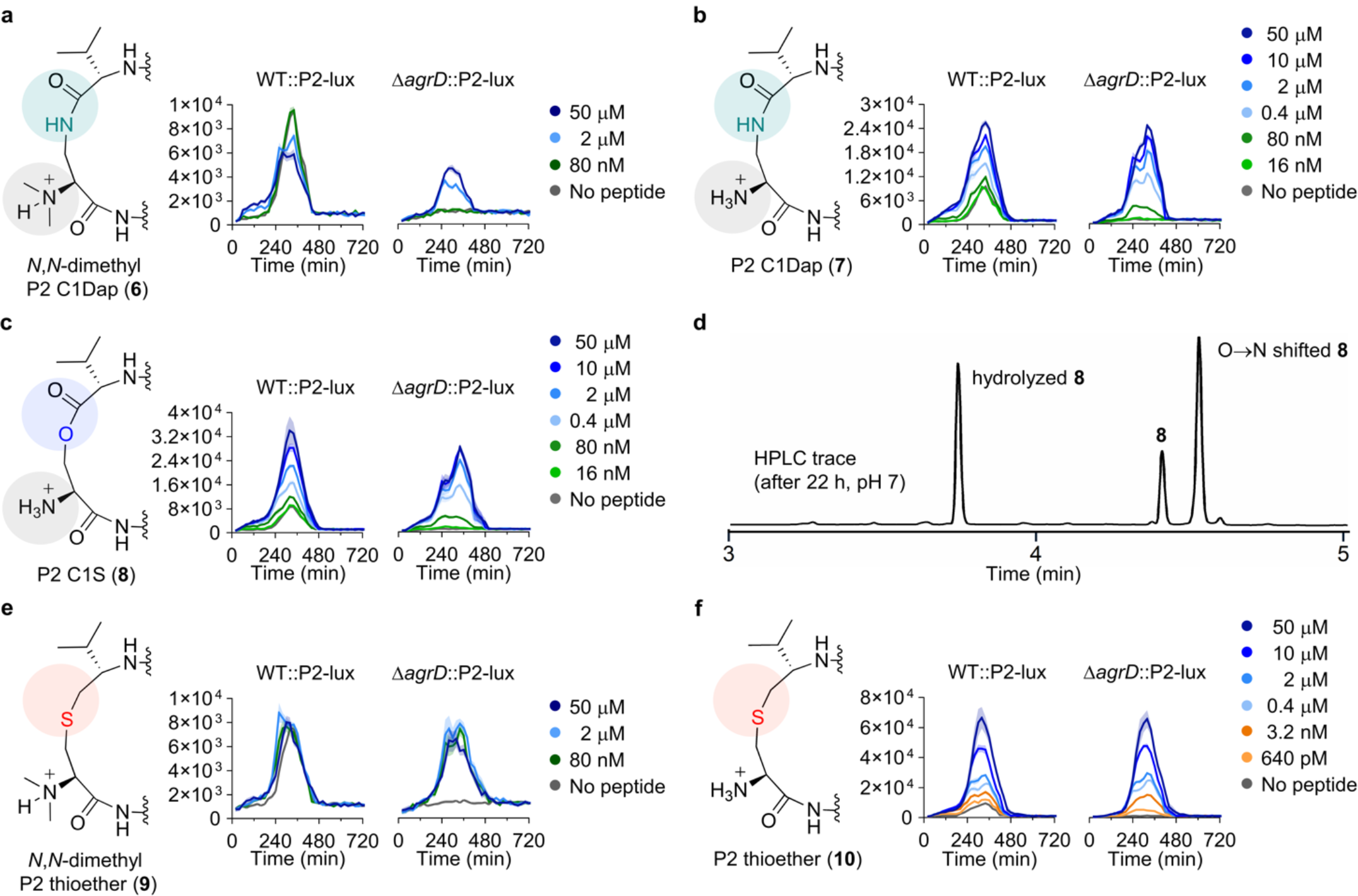
Chemical structures of peptide analogues 6–10, rearrangement evaluation of 8, and selected reporter strain assay data. **a**–**c**, Structural modifications to give **6**–**8** and effect of selected concentrations of **6**–**8** on the *agr* activity in the reporter strains of *L. monocytogenes*. **d**, HPLC trace showing the degree of rearrangement and hydrolysis of **8** after incubation in buffer for 22 h. **e**,**f**, Structural modifications to give **9** and **10** and effect of selected concentrations of **9** and **10** on the *agr* activity in the reporter strains of *L. monocytogenes*. Data are based on assays performed at 37 °C in tryptic soy broth (TSB) medium, representing at least three individual assays performed in at least duplicate unless otherwise is noted. Shadings on the graphs represent the standard error of the mean (SEM). *Based on two individual assays performed in triplicate.

Finally, we extended our series to include thioether analogues **9** and **10** (Figure 4e,f and Supplementary Figs. 16,17). When treating WT::P2-lux with the structurally closest analog of **P2**– compound **10**–we observed a striking 8-fold increase in *agr* activation compared to untreated WT. Compound **10** provided early induction in WT::P2-lux and increased *agr* activity of Δ*agrD*::P2-lux to ∼50% of the untreated WT levels at concentrations as low as 640 pM (Fig. 4f and Supplementary Fig. 18). Furthermore, at just 3.2 nM compound concentration, the luminescence levels observed for Δ*agrD*::P2-lux paralleled those of WT::P2-lux (Fig. 4f and Supplementary Fig. 18). For Δ*agrD*::P2-lux treated with compound **9**, we observed attenuated maximal activation, as for *N,N*-dimethylated analogues **1** and **6** (Fig. 4e and Supplementary Fig. 18). However, maximal induction of *agr* was observed at 80 nM concentration of **9**, rendering this compound a more potent mimic of the native AIP than the lactam **6** (Table 1).

In summary, we first investigated the mimicking of exotail-free thiolactone-containing AIPs by incorporating methyl groups at the N-terminus of **P2** to circumvent its S→N acyl shift to give **P3**. This new strategy for studying exotail-free AIPs enabled us to investigate the importance of each amino acid in the peptide through an alanine scan. While all positions are indispensable for full activity, the phenylalanine positions proved most important for retaining *agr* activation. The F2A (**2**) and F4A (**4**) analogues inhibited *agr*, but further efforts are necessary for developing potent antagonists of the *L. monocytogenes agr* system that can function as tool compounds.

Unlike what is known from studies of the paradigm *agr* systems in *S. aureus*, agonistic activity was retained when the native thiolactone functionality was substituted for a lactam (**7**) or a lactone (**8**) motif (Table 1). Moreover, an unprecedented thiolactone-to-thioether substitution, to give compound **10**, yielded the most potent inducer of the *L. monocytogenes agr* system reported so far (Table 1, Figure 4f, and Supplementary Fig. 18), which provides yet another novel architecture for studying AIPs. Comparison of lactam and thioether analogues with and without dimethylated N-termini (**6** vs **7** and **9** vs **10**; Table 1), revealed that N-methylation reduces *agr* activation, which should be considered when using this strategy to stabilize thiolactone-containing AIPs.

## Discussion

Collectively, our data show that the AgrC receptor in *L. monocytogenes* is more promiscuous with respect to activation than previously found for *S. aureus* systems. Further, **P2** is more active than **P1** and **P3**, despite its very short-lived nature, and development of stabilized analogues of **P2** further supported a role of this exotail-free thiolactone as a tight-binding native AIP. Our findings, in turn, raise several questions regarding the biological significance of this transient nature of **P2**.

It has been proposed that *agr* in *L. monocytogenes* only serves the purpose of sensing cell density^5^, and differential regulation of *agr* has been shown to depend on temperature and overlap with other regulons^18^. Further, heterogeneous *agr* activity at population level was observed for several strains of *L. monocytogenes*, and it is possible that this short-lived signal may function as a single-cell signal under certain conditions^29^.

This work adds to the potential effects of outside stimuli on *agr* regulation in *L. monocytogenes*, as the pH-dependent nature of the peptide signal (**P2**) could be speculated to provide a way to fine tune *agr* activity in response to changing pH in the environment. A compartmentalized regulatory role for **P2** could be imagined, possibly related to acidic host cell compartments such as the phagolysosome. The optimum activity of the important hemolysin, listeriolysin O (LLO), is similarly regulated. Inside the acidifying phagolysosome, LLO helps *L. monocytogenes* escape and upon release into the neutral environment of the host cell cytosol, it rapidly denatures^30,31^.

Further, *L. monocytogenes* infection relies on traversing the environments of the gastro-intestinal tract and its varying pH values^1^. It is therefore intriguing to start thinking about how such an adaptive peptide signal might play a role during this process, possibly even in concert with the more chemically stable peptides **P1** and **P3**.

This research provides essential insight into the peptide-mediated cell-to-cell communication of *L. monocytogenes* and provides tool compounds for future studies of the biological importance of *agr* regulation in this bacterium. More generally, several other bacteria also produce exotail-free AIPs, and we hope that the chemical framework provided in this study can help guide further research in their respective physiological regulation through cell-to-cell communication^21,32,33^.

## Methods

### Assay protocol for monitoring of S→N acyl shift

A microcentrifugal tube (1.5 mL) containing a solution of phosphate buffer (pH = 7, 100 mM) and MeCN (1:1) was preheated to 37 °C. Assays were initiated (t = 0 min) by the addition of 1 μL of peptide DMSO stock solution per 100 μL of buffer– MeCN solution (100 μM final peptide conc.) with subsequent mixing using an orbital shaker. Reactions were incubated at 37 °C under constant stirring and samples were taken at relevant time points and mixed (20:1, v/v) with a solution of TFA–water (1:1, v/v) to prevent further rearrangement. Samples were then analyzed by UPLC (gradient: 5–95 B% over 5 min) and ratios between exotail-free thiolactone and homodetic peptide was determined by integration of the areas under the corresponding peaks at a wavelength of λ = 215 nm. Rate constants (*k*) were calculated as the slope of plotting S→N acyl shift progression over time using GraphPad Prism 9.0 software.

### Luciferase-based assay protocol for measuring *agr* activity in *L. monocytogenes*

*L. monocytogenes* bioluminescent reporter strains (WT::P2-lux or Δ*agr*D::P2-lux) were streaked onto TSA plates containing 5 μg/mL chloramphenicol and grown at 37 °C overnight. Single colonies were used to inoculate overnight cultures in TSB/BHI containing 5 μg/mL chloramphenicol for use in the assay. To the outer wells of a white 96-well assay plate with clear bottom was added sterile water, to minimize evaporation from the remaining 60 wells during incubation. Peptide stocks in DMSO (5 mM) were diluted in TSB/BHI to the appropriate concentration (10× the final assay concentration) and 20 μL was added to the respective wells (20 μL TSB added to control wells without peptide). The optical density at 600 nm (OD_600_) of overnight cultures was measured and cultures were diluted to an OD_600_ = 0.01 in fresh TSB containing flavin mononucleotide (11.1 mg/L, final assay concentration; ∼10 mg/L, ∼26 μM). Subsequently, 180 μL of diluted overnight culture was added to the relevant wells of the assay plate. The assay plate was incubated in a BioTek Synergy H1 microplate reader at 37 °C with continuous double orbital shaking. Luminescence (gain = 255) and OD_600_ was measured every 20 min. for 14 hours.

### General procedures for automated solid-phase peptide synthesis (SPPS)

The following Fmoc-protected amino acids with side chain protecting groups (Fmoc-AA-OH) were used for the automated synthesis of peptides: Fmoc-Ala-OH, Fmoc-Met-OH, Fmoc-Phe-OH, and Fmoc-Val-OH. Automated peptide synthesis was carried out on a Biotage SyroWave™ synthesizer using iterative coupling and deprotection steps on 3-4-amino-(methylamino)benzoic acid (MeDbz)-Gly-ChemMatrix resin (0.45 mmol/g) or pre-loaded 2-chlorotrityl chloride (Cl-Trt)-polystyrene resin (0.7–0.9 mmol/g). *Loading of an amino acid residue to MeDbz-resin*. The coupling reaction was performed for 90 min at room temperature with short vortexing intervals (10 s) using stock solutions of Fmoc-AA-OH in DMF (5.00 equivalents to the resin loading, 0.5 M), HATU in DMF (4.90 equivalents, 0.5 M) and *i-* Pr_2_NEt in NMP (10.0 equivalents, 2.0 M) at a final concentration of 0.2 M for Fmoc-AA-OH. The coupling was followed by washing of the resin with DMF (5 × 1 min) and the procedure was repeated to achieve a double coupling. *Standard coupling of an amino acid residue*. The coupling reaction was performed for 40 min at room temperature with short vortexing intervals (10 s) using stock solutions of Fmoc-AA-OH in DMF (5.00 equivalents to the resin loading, 0.5 M), HBTU in DMF (4.90 equivalents, 0.5 M) and *i-*Pr_2_NEt in NMP (10.0 equivalents, 2.0 M) at a final concentration of 0.2 M for Fmoc-AA-OH. The coupling reaction was followed by washing of the resin with DMF (5 × 1 min) and the procedure was repeated to achieve a double coupling. Fmoc removal was performed in two stages: 1) piperidine in DMF (2:3, v/v) for 3 min and 2) piperidine in DMF (1:4, v/v) for 12 min with short vortexing intervals (10 s). The deprotection was followed by washing of the resin with DMF (3 × 1 min), CH_2_Cl_2_ (1 × 1 min) and DMF (3 × 1 min).

### General procedure for manual coupling step of *N*-terminal amino acids

The following amino acids were used in the manual coupling step as *N*-terminal amino acid: *N,N*-dimethyl-Cys(Trt)-OH, Fmoc-Dap(Alloc)-OH, *N,N*-dimethyl-Dap(Fmoc)-OH, Boc-Cys(S*t*-Bu)-OH, *N,N*-dimethyl-Cys[Val(Fmoc)]-OH, and Fmoc-Ser(TBDMS)-OH. After automated peptide elongation, the resin was transferred into a polypropylene syringe equipped with a fritted disk using CH_2_Cl_2_ and the resin was then washed with DMF (3 × 1 min). The coupling reaction was performed using amino acid (2.00 equivalents to the resin loading), HATU (2.00 equivalents), and *i-*Pr_2_NEt (4.00 equivalents) in DMF (final concentration = 0.08 M for Fmoc-AA-OH) at room temperature under light shaking. After 2 h, the coupling mixture was removed by suction and the resin was washed with DMF (3 × 1 min) and CH_2_Cl_2_ (3 × 1 min).

### General procedure for the synthesis of *N,N*-dimethyl-thiolactone peptides

*N-acyl-benzimidazolinone (Nbz) formation:* After completed peptide elongation on 20.0 μmol MeDbz-Gly-ChemMatrix resin (see Supplementary Fig. 7), the peptidyl-MeDbz resin (1.00 equivalents) was washed with CH_2_Cl_2_ (5 × 1 min). A solution of 4-nitrophenyl-chloroformate (5.00 equivalents) in CH_2_Cl_2_ (conc: 0.1 M) was added to the resin and the suspension was agitated for 30 min. The resin was then washed with CH_2_Cl_2_ (2 × 1 min) and the procedure was repeated. The resin was then washed with CH_2_Cl_2_ (3 × 1 min) and DMF (3 × 1 min) and a solution of *i-*Pr_2_NEt (25.0 equivalents, 0.5 M) in DMF was added to the resin. After 15 min, the resin was washed with DMF (3 × 1 min) and the procedure was repeated. The peptidyl-MeNbz-resin was then washed with DMF (3 × 1 min), *i-* Pr_2_NEt in DMF (5%, v/v) (3 × 1 min), DMF (3 × 1 min), MeOH (3 × 1 min), and CH_2_Cl_2_ (3 × 1 min) and dried under high vacuum. *On-resin cleavage-inducing cyclization:* The dried peptidyl-MeNbz-Gly-ChemMatrix resin (20.0 μmol, 1.00 equivalents; Supplementary Fig. 13) was treated with a deprotection cocktail (2.0 mL, TFA–*i-*Pr_3_SiH–water, 94:3:3, v/v/v) for 1 h and the TFA cocktail was subsequently removed from the resin. The resin was washed with CH_2_Cl_2_ (3 × 1 min), DMF (3 × 1 min), and CH_2_Cl_2_ (3 × 1 min) and dried under suction for several minutes. Cyclization buffer (5.0 mL, phosphate buffer (0.2 M, pH 6.8)–MeCN, 1:1, v/v) was added to the resin (final concentration = 4.0 mM) and the suspension was agitated at 50 °C. After 2 h, the solution was collected, and the resin was rinsed with fresh cyclization buffer. The combined peptide-containing washings were pooled and purified by preparative HPLC to afford the *N,N*-dimethyl-thiolactone peptides **1**–**5** after lyophilization.

## Data availability

The authors declare that the data supporting the findings of this study are available within the paper and its supplementary information files.

## Acknowledgements

We gratefully acknowledge the Independent Research Fund Denmark−Natural Sciences (Grant No. 0135-00427B; C.A.O.) and the LEO Foundation Open Competition Grant program (LF-OC-20-000517; M.S.B. and H.I., LF-OC-19-000039; C.A.O., and LF-OC-21-000901; C.A.O.) for financial support. We thank Prof. Christian Riedel (University of Ulm) for generously providing the bacterial strains used in the study.

## Author contributions

**Benjamin S. Bejder:** conceptualization, investigation, formal analysis, methodology, visualization, writing–original draft, writing–review, editing; **Fabrizio Monda:** investigation, formal analysis, methodology, writing–review & editing; **Bengt H. Gless:** conceptualization, investigation, formal analysis, methodology, supervision, visualization, writing–review & editing; **Martin S. Bojer:** investigation, formal analysis, methodology, supervision, writing–review & editing; **Hanne Ingmer:** funding acquisition, supervision, writing–review & editing; **Christian A. Olsen:** conceptualization, formal analysis, funding acquisition, project administration, resources, supervision, writing-original draft, writing–review & editing.

## Additional information

Supplementary figures, experimental methods, chemical synthesis and compound characterization data, as well as copies of HPLC traces, ^1^H and ^13^C NMR spectra is available for the manuscript.

