## Supplementary Information for "A short-lived peptide signal regulates cell-to-cell communication in *Listeria monocytogenes*"

### Supplementary Figures

#### Optimization of luminescence-based *agr* reporter assay of *Listeria monocytogenes* and evaluation of P1–P3

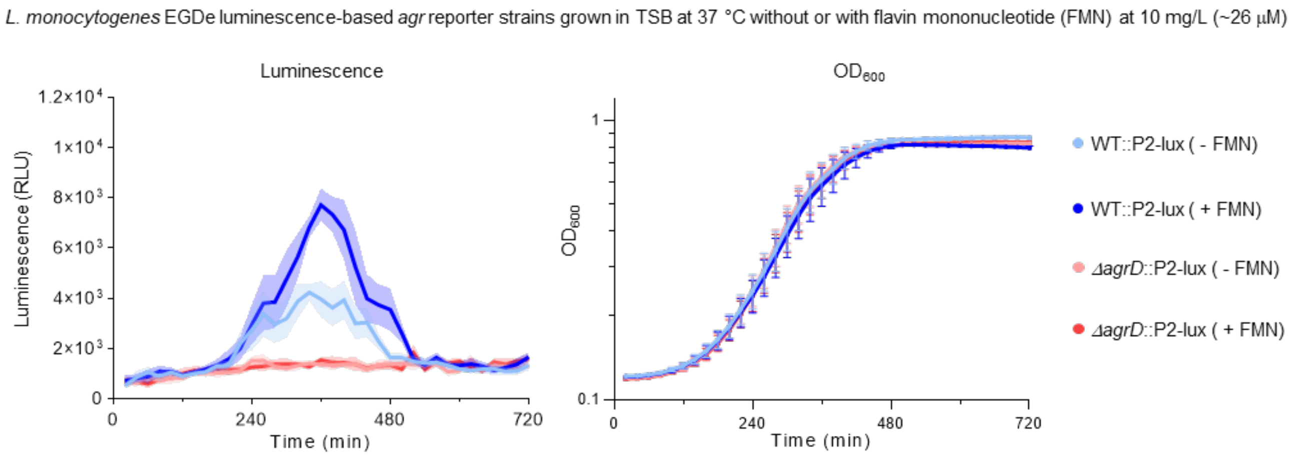

**Supplementary figure 1.** Reporter strain measurements from ***L. monocytogenes* EGDe WT::P2-lux** and ***L. monocytogenes* EGDe *ΔagrD*::P2-lux** grown in tryptic soy broth (**TSB**) medium, without or supplemented with flavin mononucleotide (10 mg/L, ~26 µM) at **37 °C**. Graphs represent transcriptional activity from the *agr* promoter measured in relative luminescence units (RLU), bacterial cell density measured as optical density at 600 nm ($OD_{600}$) and *agr* activity adjusted to cell density (RLU/$OD_{600}$). All curves represent the mean of two biological replicates with technical triplicates. Error bars and shaded areas represent the standard error of the mean (SEM).

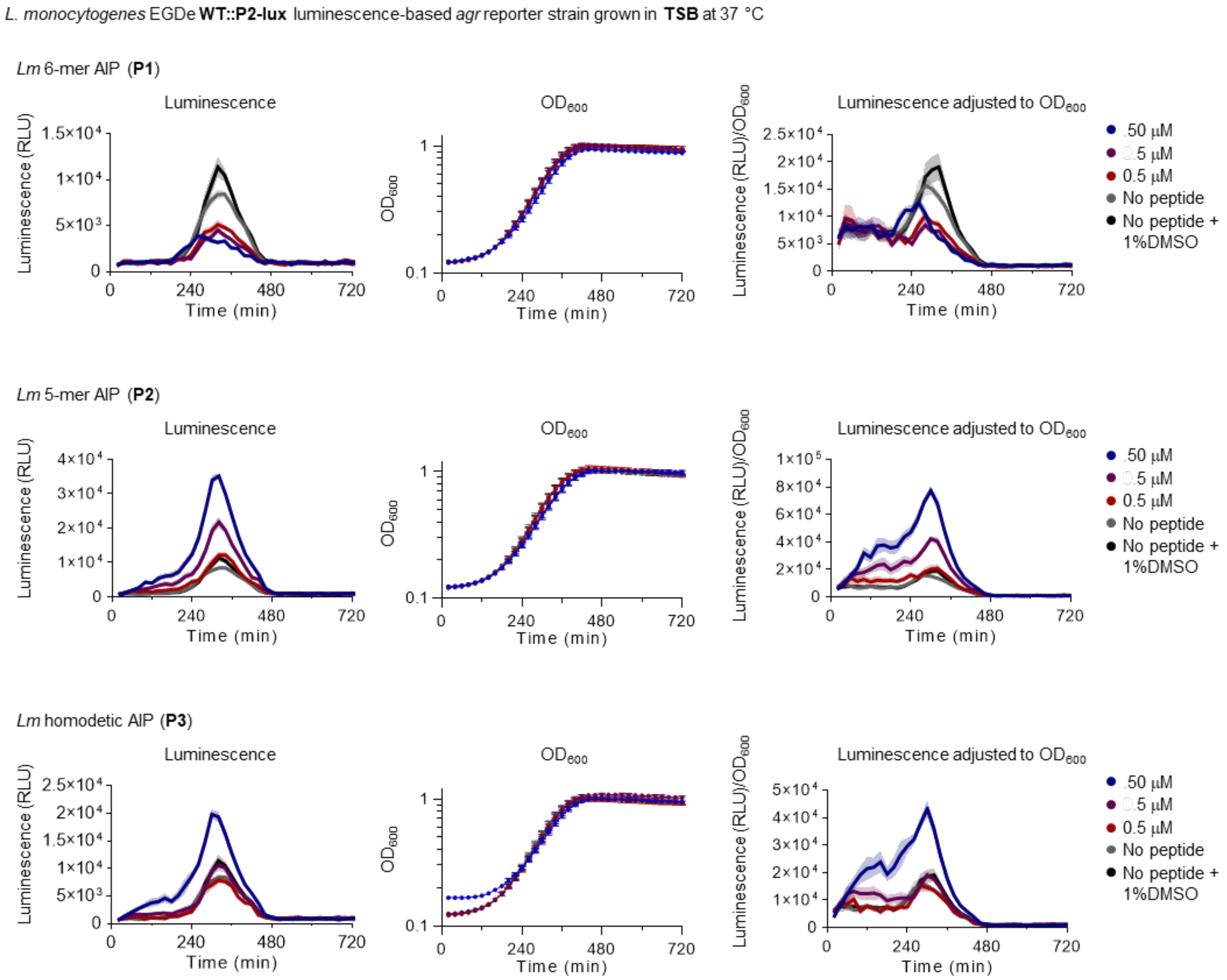

**Supplementary figure 2.** Measurements from the ***L. monocytogenes*** **EGDe WT::P2-lux** reporter strain treated with synthetic peptides (**P1**, **P2**, and **P3)** at selected concentrations (**50-0.5 µM**) and grown in tryptic soy broth (**TSB**) medium at **37 °C**. Graphs represent transcriptional activity from the *agr* promoter measured in relative luminescence units (RLU), bacterial cell density measured as optical density at 600 nm ($OD_{600}$) and *agr* activity adjusted to cell density (RLU/$OD_{600}$). All curves represent the mean of three biological replicates with at least technical duplicates. Error bars and shaded areas represent the standard error of the mean (SEM).

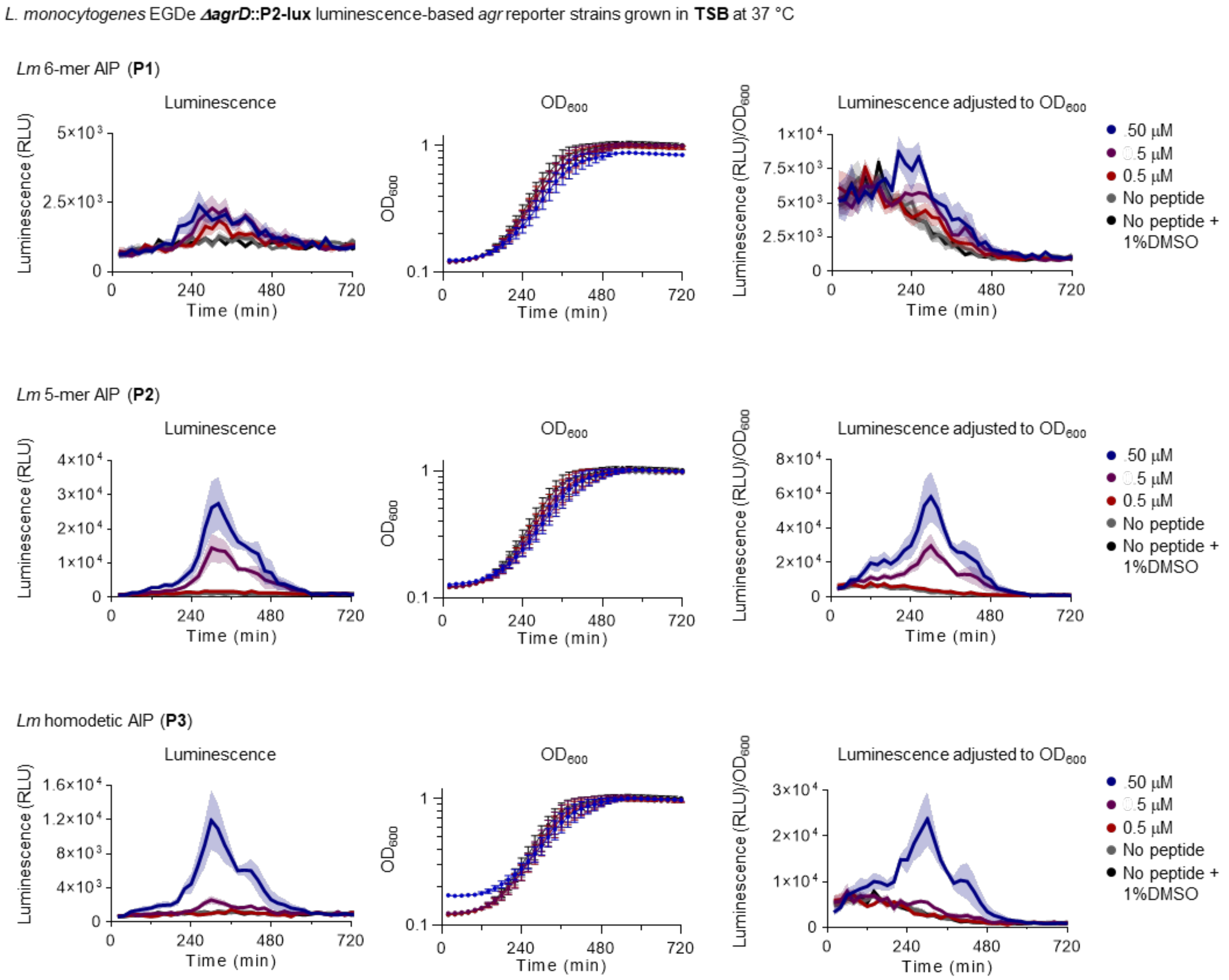

**Supplementary figure 3.** Measurements from the ***L. monocytogenes* EGDe Δ*agrD*::P2-lux** reporter strain treated with synthetic peptides (**P1**, **P2**, and **P3)** at selected concentrations (**50-0.5 µM**) and grown in tryptic soy broth (**TSB**) medium at **37 °C**. Graphs represent transcriptional activity from the *agr* promoter measured in relative luminescence units (RLU), bacterial cell density measured as optical density at 600 nm ($OD_{600}$) and *agr* activity adjusted to cell density ($RLU/OD_{600}$). All curves represent the mean of three biological replicates with at least technical duplicates. Error bars and shaded areas represent the standard error of the mean (SEM).

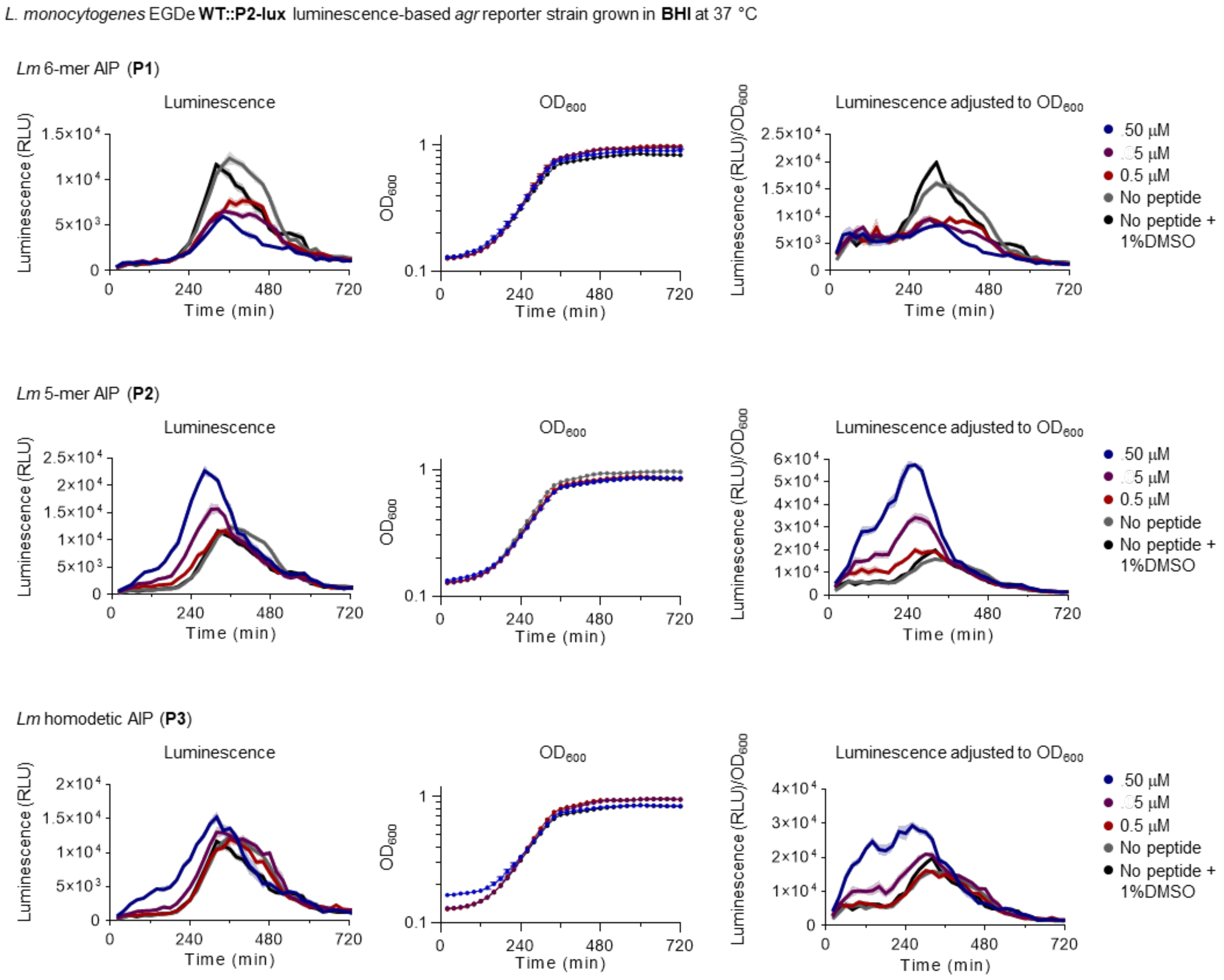

**Supplementary figure 4.** Measurements from the ***L. monocytogenes*** **EGDe WT::P2-lux** reporter strain treated with synthetic peptides (**P1**, **P2**, and **P3)** at selected concentrations (**50-0.5 µM**) and grown in brain heart infusion (**BHI**) medium at **37 °C**. Graphs represent transcriptional activity from the *agr* promoter measured in relative luminescence units (RLU), bacterial cell density measured as optical density at 600 nm ($OD_{600}$) and *agr* activity adjusted to cell density (RLU/$OD_{600}$). All curves represent the mean of three biological replicates with at least technical duplicates. Error bars and shaded areas represent the standard error of the mean (SEM).

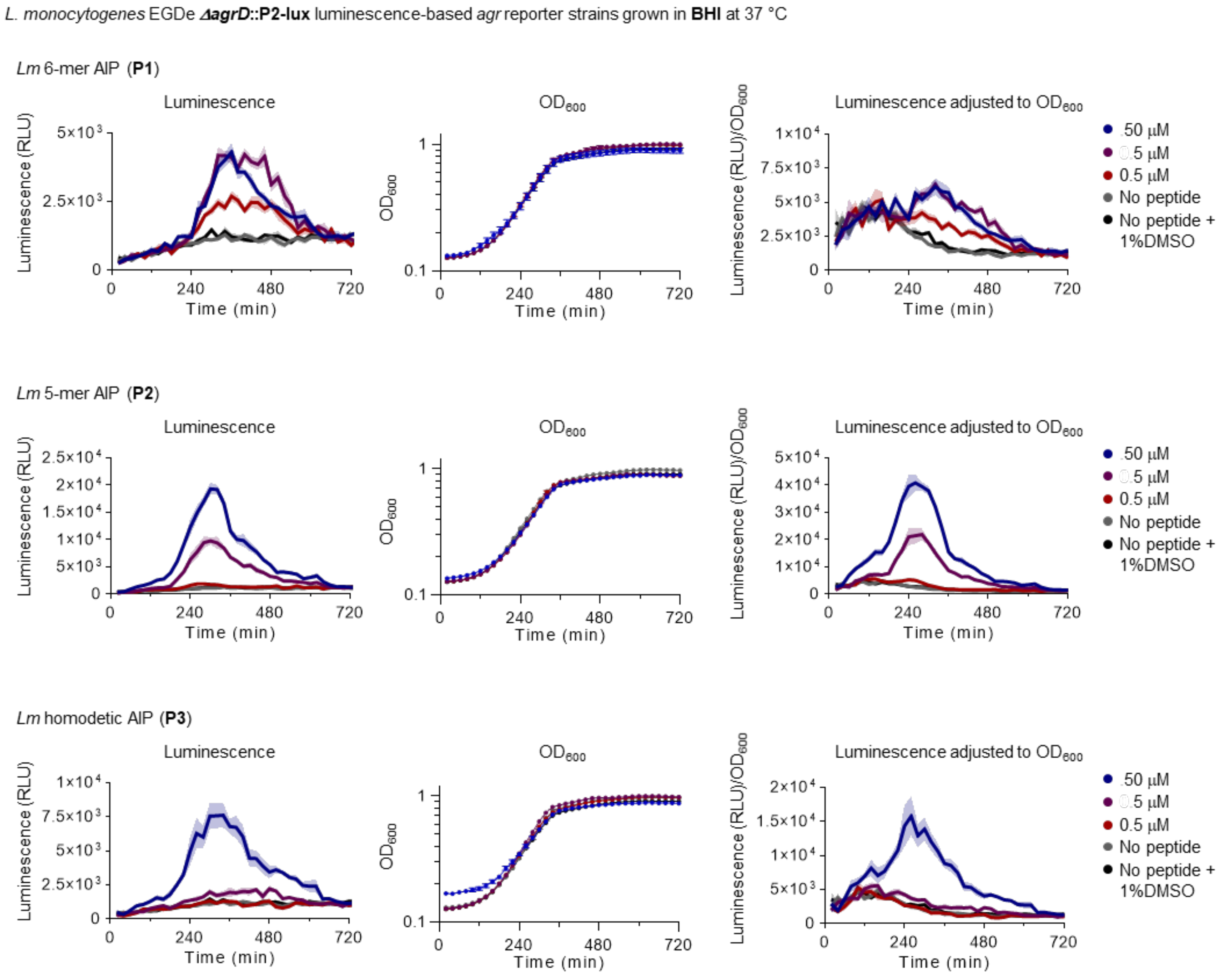

**Supplementary figure 5.** Measurements from the ***L. monocytogenes*** **EGDe *ΔagrD*::P2-lux** reporter strain treated with synthetic peptides (**P1**, **P2**, and **P3)** at selected concentrations (**50–0.5 µM**) and grown in brain heart infusion (**BHI**) medium at **37 °C**. Graphs represent transcriptional activity from the *agr* promoter measured in relative luminescence units (RLU), bacterial cell density measured as optical density at 600 nm ($OD_{600}$) and *agr* activity adjusted to cell density (RLU/$OD_{600}$). All curves represent the mean of three biological replicates with at least technical duplicates. Error bars and shaded areas represent the standard error of the mean (SEM).

#### Additional discussion of assay conditions

While previously observing a bimodal luminescence output for WT::P2-lux grown in BHI medium, we did not observe such a trend when supplementing with flavin mononucleotide and assaying at 37 °C (Supplementary Figs 4–5). The trends previously observed for **P1** and **P3** at 30 °C were recapitulated under the optimized conditions: WT::P2-lux with **P1** resulted in inhibition of the signal in both BHI and TSB media, while giving rise to a slight induction in the *ΔagrD*::P2-lux reporter strain after 240 min, which was most pronounced in BHI medium (Supplementary Figs 2–5); the homodetic peptide **P3** did not cause inhibition of WT::P2-lux, but produced early induction of *agr* in both TSB and BHI media at 50 µM, and also at 5 µM when adjusting luminescence measurements to the cell density ($OD_{600}$) (Supplementary Figs 2 and 4). Though, previous results at 30 °C demonstrated similar trends, the improved assay conditions uncovered the ability of **P3** to increase peak luminescence measurements (t ~330 min) by 2-fold compared to the untreated WT::P2-lux control in TSB (Supplementary Fig 2). The effects of **P3** against WT::P2-lux were reflected in assays with *ΔagrD*::P2-lux, where significant potency was only observed at 50 µM (Supplementary Fig 3). At this concentration, **P3** was capable of reaching untreated WT::P2-lux *agr* activity in TSB, but not fully in BHI medium (Supplementary Figs 3 and 5).

#### Assay data for *N*-acetyl-P2 (SP1)

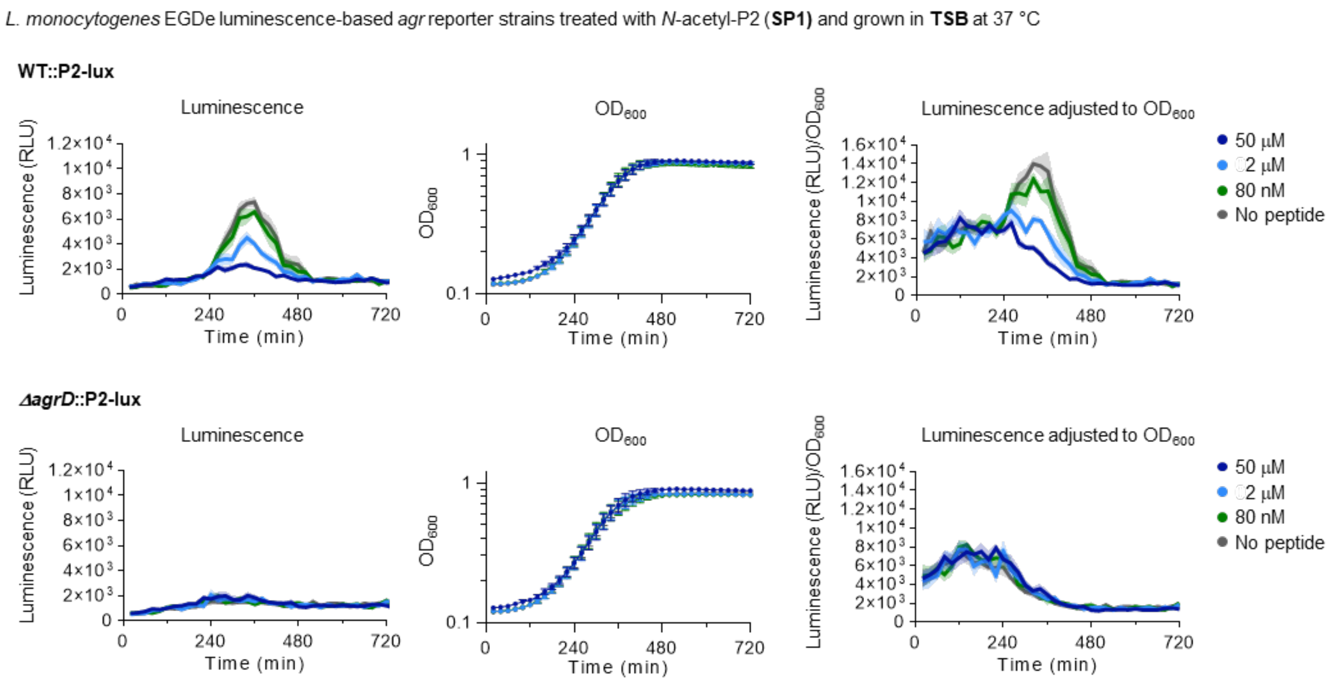

**Supplementary figure 6.** Measurements from the ***L. monocytogenes*** **EGDe WT::P2-lux** and ***L. monocytogenes*** **EGDe *ΔagrD*::P2-lux** reporter strains treated with *N*-acetyl-P2 (**SP1)** at selected concentrations (**50–0.08 µM**) and grown in tryptic soy broth (**TSB**) medium at **37 °C**. Graphs represent transcriptional activity from the *agr* promoter measured in relative luminescence units (RLU), bacterial cell density measured as optical density at 600 nm ($OD_{600}$) and *agr* activity adjusted to cell density ($RLU/OD_{600}$). All curves represent the mean of three biological replicates with at least technical duplicates. Error bars and shaded areas represent the standard error of the mean (SEM).

#### Synthesis of *N,N*-dimethyl thiolactone peptides

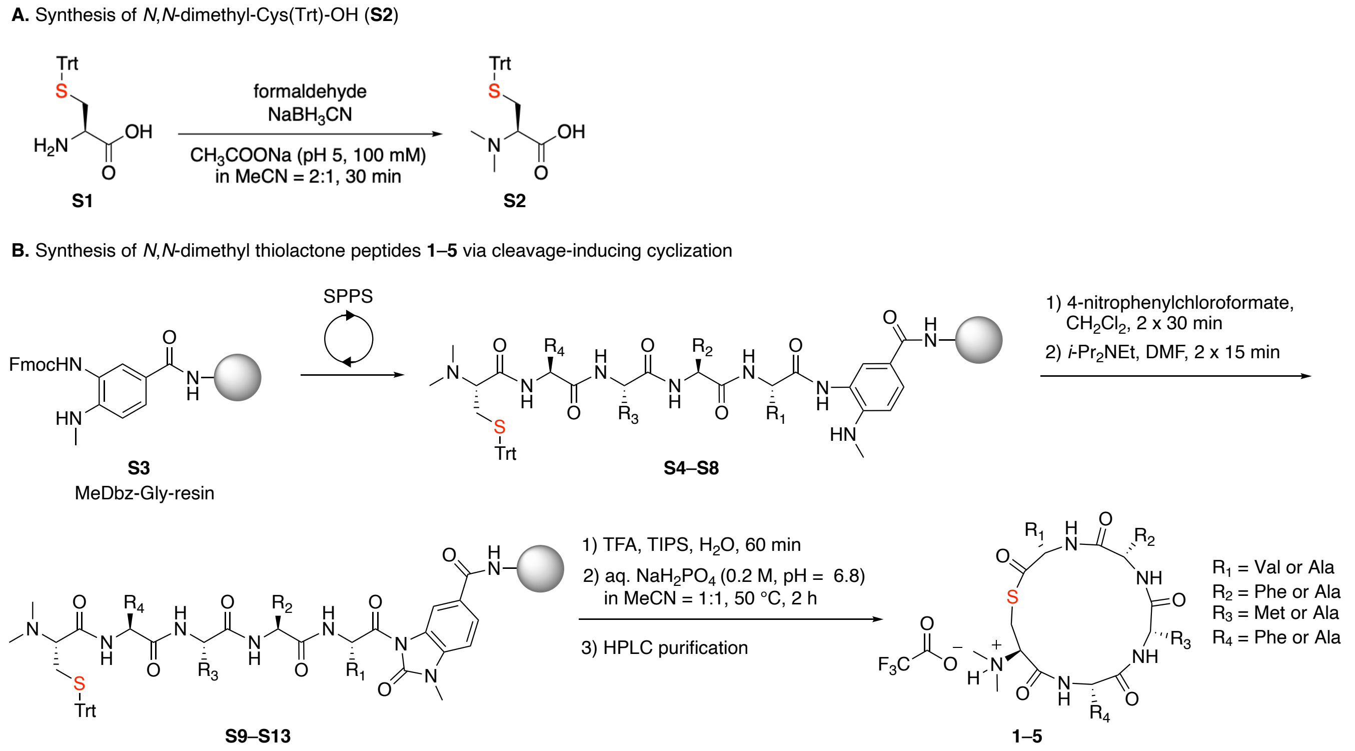

**Supplementary figure 7.** (**A**) Synthesis of *N,N*-dimethyl-Cys(Trt)-OH (**S2**) by reductive amination of H-Cys(Trt)-OH (**S1**). (**B**) Synthesis of *N,N*-dimethyl thiolactone peptides **1–5** using a cleavage-inducing cyclization protocol.[1] Protected linear peptides **S4–S8** were synthesized on 3-4-amino-(methylamino)benzoic acid (MeDbz)-Gly-resin and the last coupling steps was performed using **S2**. The MeDbz linker was activated to its corresponding *N*-acyl-benzimidazolinone (MeNbz) form and resin-bound MeNbz-peptides **S9–S13** were deprotected and followed by a cleavage-induced cyclization step, which afforded the *N,N*-dimethyl thiolactone peptides **1–5**.

#### Hydrolytic stability of *N,N*-dimethyl-P2 (1) and assay data for compounds 1–5

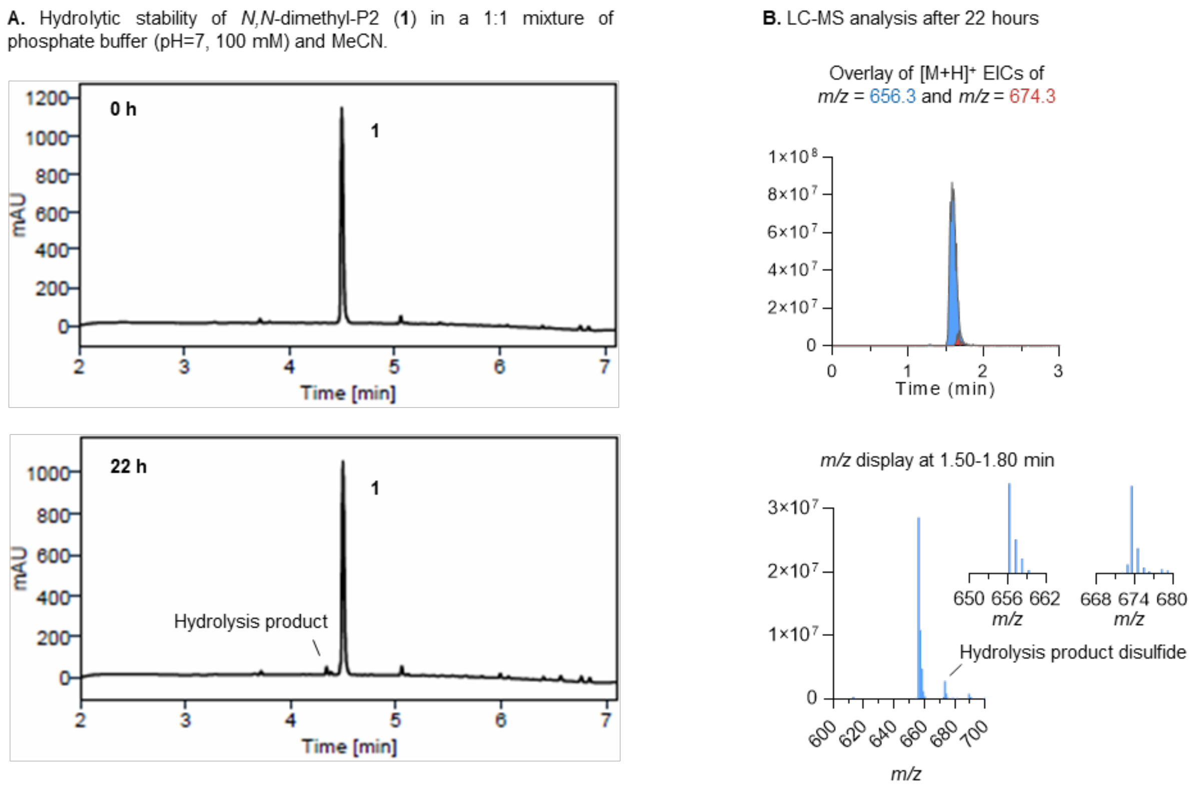

**Supplementary figure 8.** (**A**) Ultra-performance liquid chromatography (UPLC) traces of *N,N*-dimethyl-P2 (**1**) before and after being stirred for 22 hours at 37 °C in a 1:1 mixture of phosphate buffer (pH=7, 100 mM) and MeCN, displaying limited extent of hydrolysis (Purity >90% after 22 hours). (**B**) Extracted ion chromatogram (EIC) overlay of the $[M+H]^+$ adducts of **1** (*m/z* = 656.3, shown in blue) and hydrolyzed **1** (*m/z* = 674.3, shown in red), along with a mass spectrum displaying their presence after 22 hours.

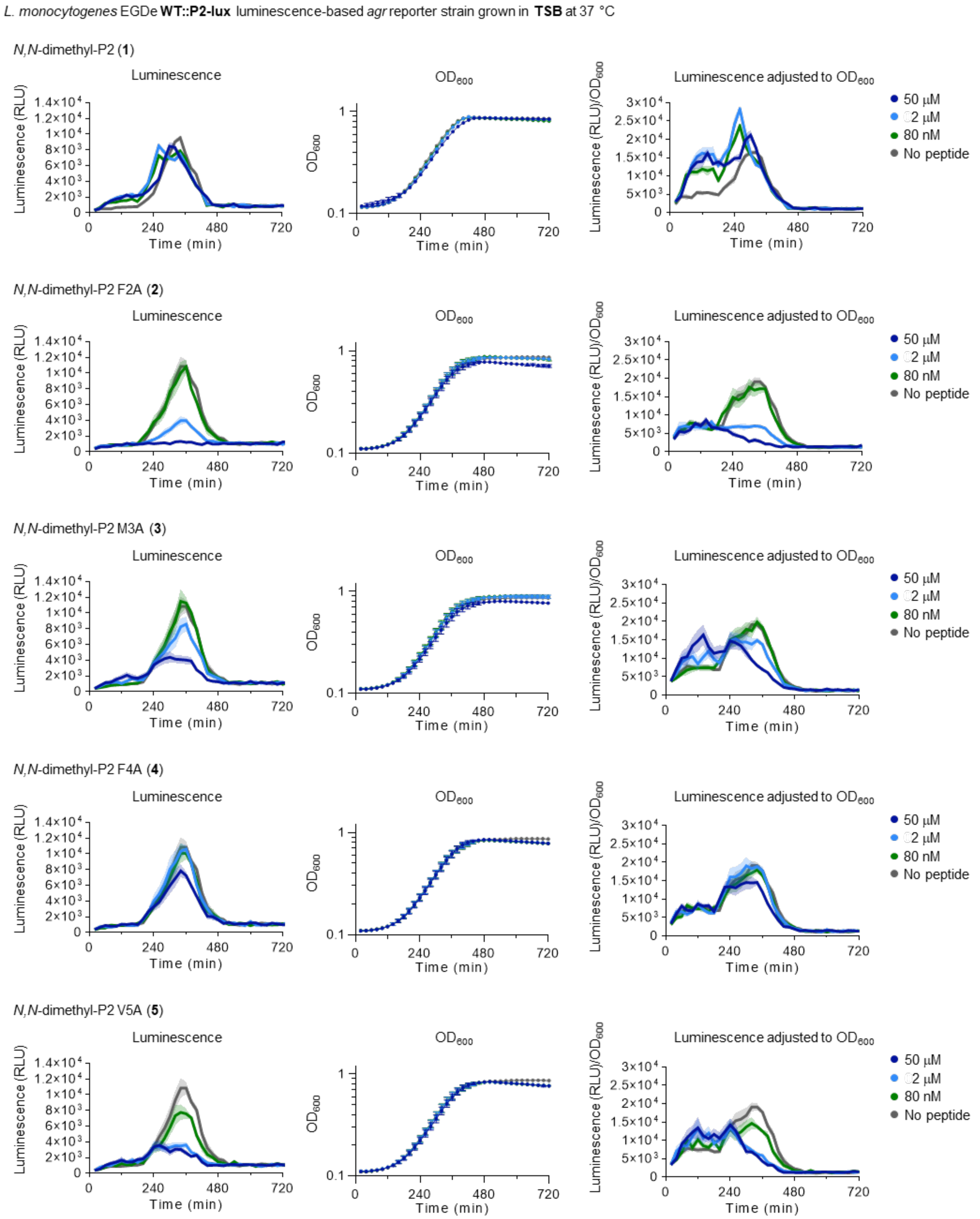

**Supplementary figure 9.** Measurements from the ***L. monocytogenes*** **EGDe WT::P2-lux** reporter strain treated with synthetic peptides (**1–5**) at selected concentrations (**50–0.08 µM**) and grown in tryptic soy broth (**TSB**) medium at **37 °C**. Graphs represent transcriptional activity from the *agr* promoter measured in relative luminescence units (RLU), bacterial cell density measured as optical density at 600 nm ($OD_{600}$) and *agr* activity adjusted to cell density ($RLU/OD_{600}$). All curves represent the mean of three biological replicates with at least technical duplicates. Error bars and shaded areas represent the standard error of the mean (SEM).

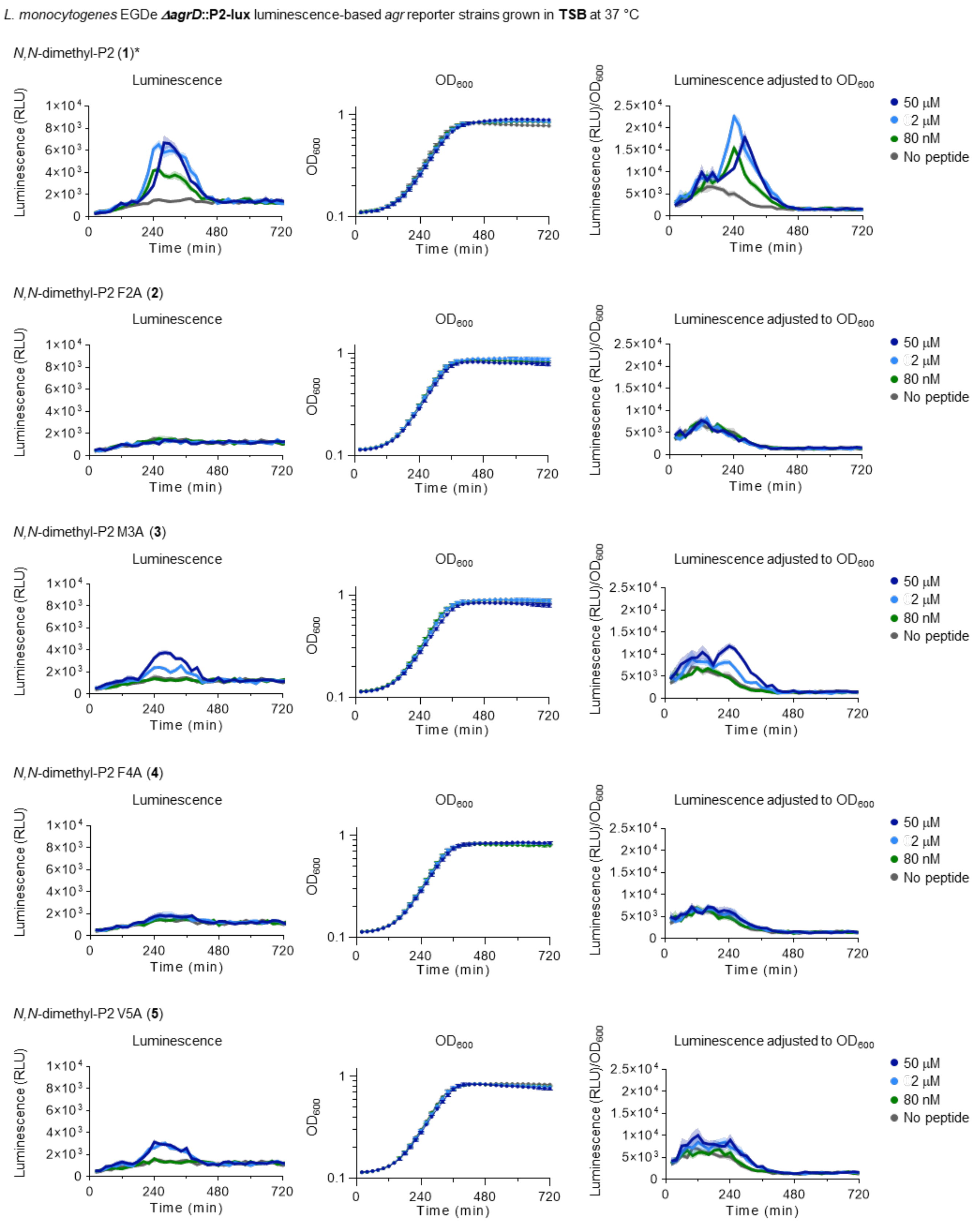

**Supplementary figure 10.** Measurements from the ***L. monocytogenes*** **EGDe *ΔagrD*::P2-lux** reporter strain treated with synthetic peptides (**1–5**) at selected concentrations (**50–0.08 µM**) and grown in tryptic soy broth (**TSB**) medium at **37 °C**. Graphs represent transcriptional activity from the *agr* promoter measured in relative luminescence units (RLU), bacterial cell density measured as optical density at 600 nm ($OD_{600}$) and *agr* activity adjusted to cell density ($RLU/OD_{600}$). All curves represent the mean of three biological replicates with at least technical duplicates. Error bars and shaded areas represent the standard error of the mean (SEM). *Two biological replicates.

### Synthesis and biological evaluation of lactam analogues 6 and 7

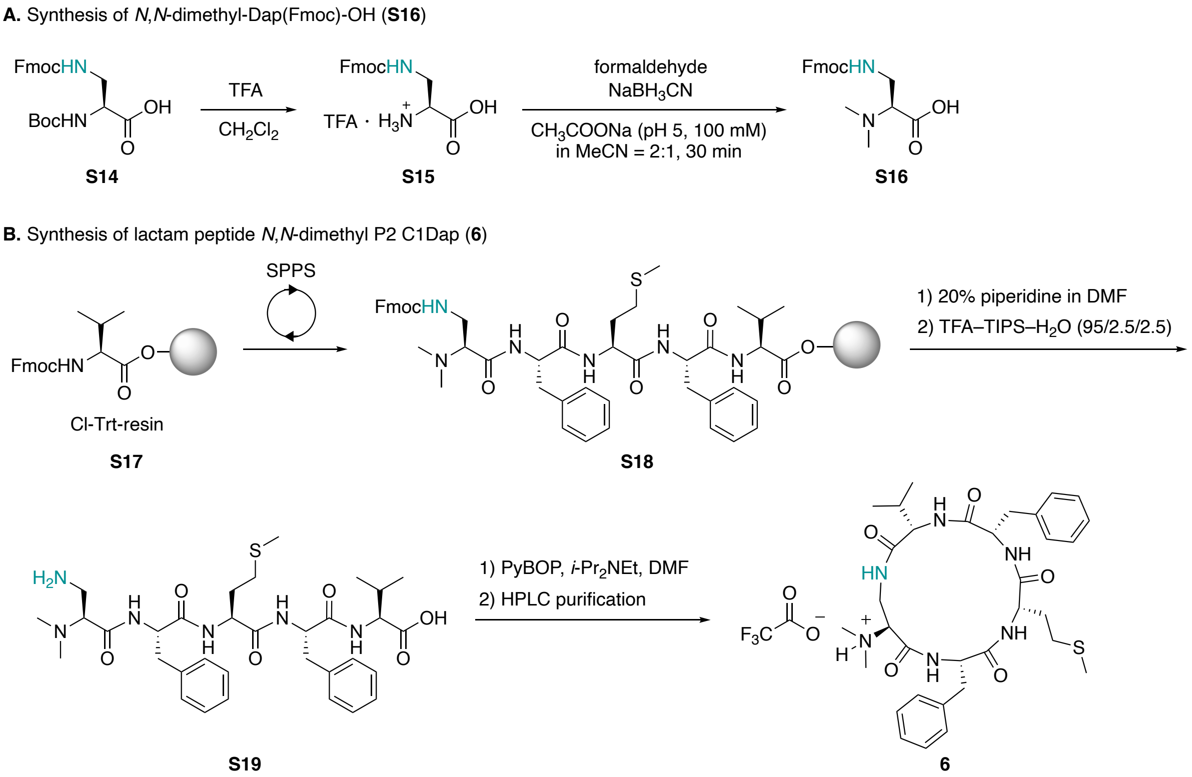

**Supplementary figure 11.** (**A**) Synthesis of *N,N*-dimethyl-Dap(Fmoc)-OH (**S16**) starting from Boc-Dap(Fmoc)-OH (**S14**) followed by reductive amination of deprotected **S15**. (**B**) Synthesis of lactam peptide *N,N*-dimethyl P2 C1Dap (**6**). The linear peptide **S18** was synthesized on Fmoc-Val-loaded 2-chlorotrityl chloride (Cl-Trt)-resin **S17** and the last coupling step was performed using **S16**. Resin-bound peptide **S18** was treated with piperidine and subsequently released from the resin to yield the linear peptide **S19**, which was cyclized via amide bond formation using PyBOP to afford the lactam peptide **6**.

Synthesis of lactam peptide P2 C1Dap (**7**)

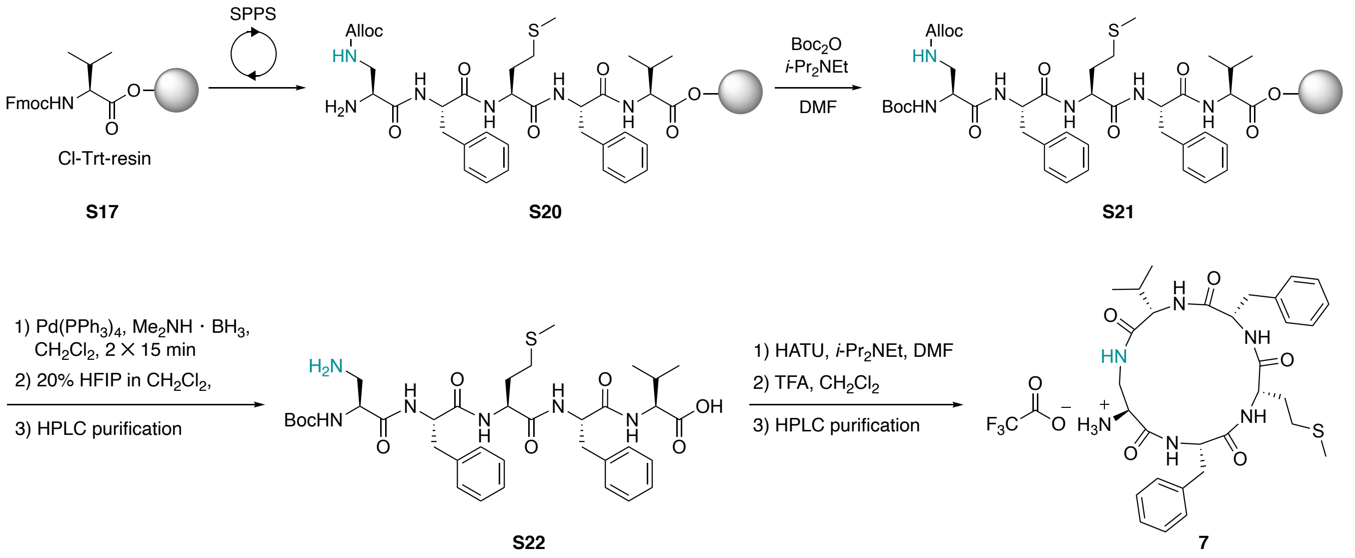

**Supplementary figure 12.** Synthesis of lactam peptide P2 C1Dap (**7**). Partially-protected linear peptide **S20** was synthesized on Fmoc-Val-loaded Cl-Trt-resin **S17** and the last coupling step was performed using Fmoc-Dap(Alloc)-OH followed by an Fmoc removal step. The *N*-terminus was Boc protected to give fully-protected peptide **S21**, which was treated to with $Pd(PPh_3)_4$ and subsequently released from the resin to yield the partially protected peptide **S22**. The linear peptide **S22** was cyclized via amide bond formation using HATU and treated with trifluoroacetic acid (TFA) to afford the lactam peptide **7**.

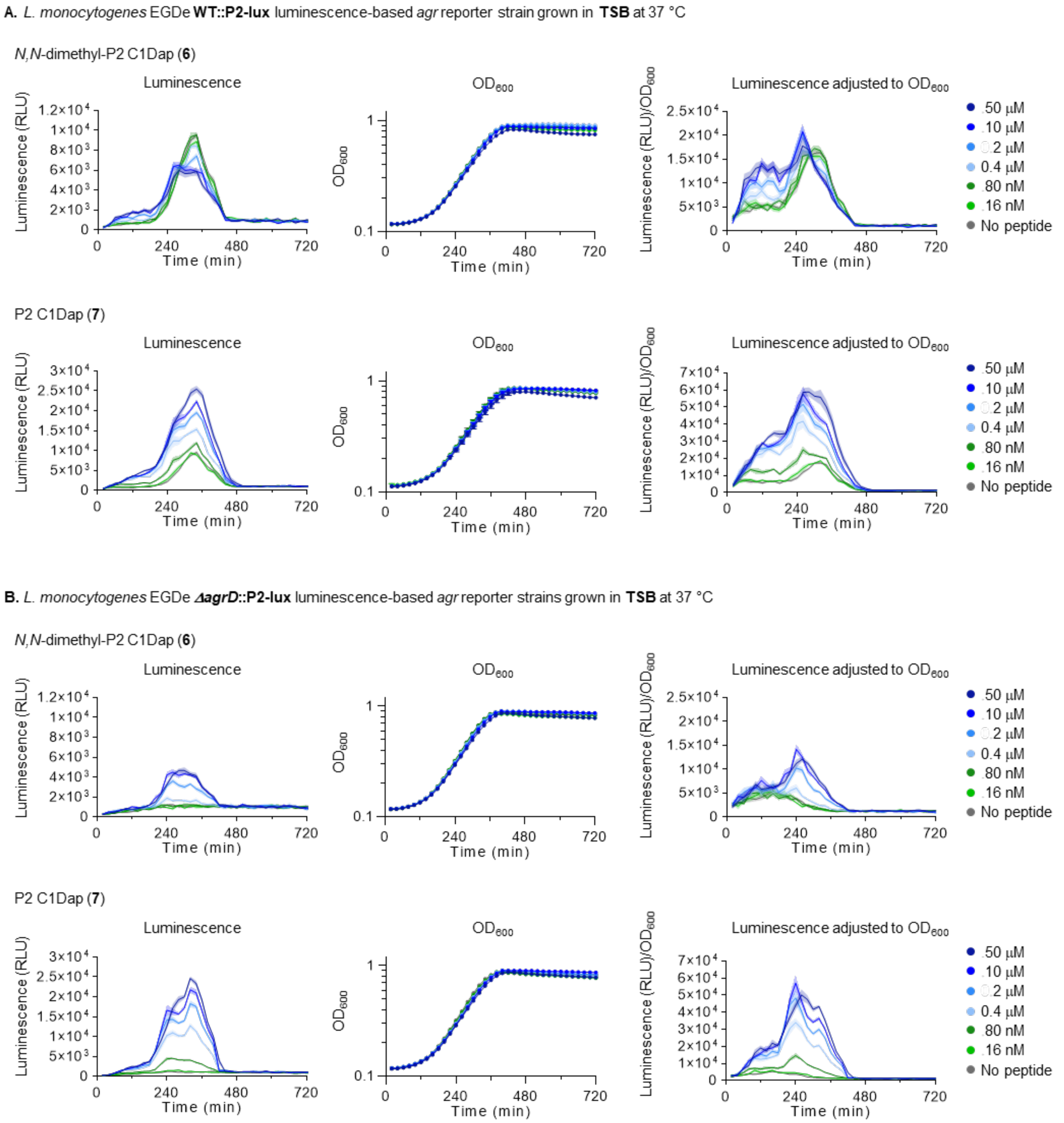

**Supplementary figure 13.** Reporter strain measurements from (**A**) ***L. monocytogenes*** **EGDe WT::P2-lux** and (**B**) ***L. monocytogenes*** **EGDe** ***ΔagrD*****::P2-lux** treated with synthetic peptides (**6** and **7**) at selected concentrations (**50–0.016 µM**) and grown in tryptic soy broth (**TSB**) medium at **37 °C**. Graphs represent transcriptional activity from the *agr* promoter measured in relative luminescence units (RLU), bacterial cell density measured as optical density at 600 nm ($OD_{600}$) and *agr* activity adjusted to cell density ($RLU/OD_{600}$). All curves represent the mean of three biological replicates with at least technical duplicates. Error bars and shaded areas represent the standard error of the mean (SEM).

#### Synthesis of lactone-containing peptide (8)

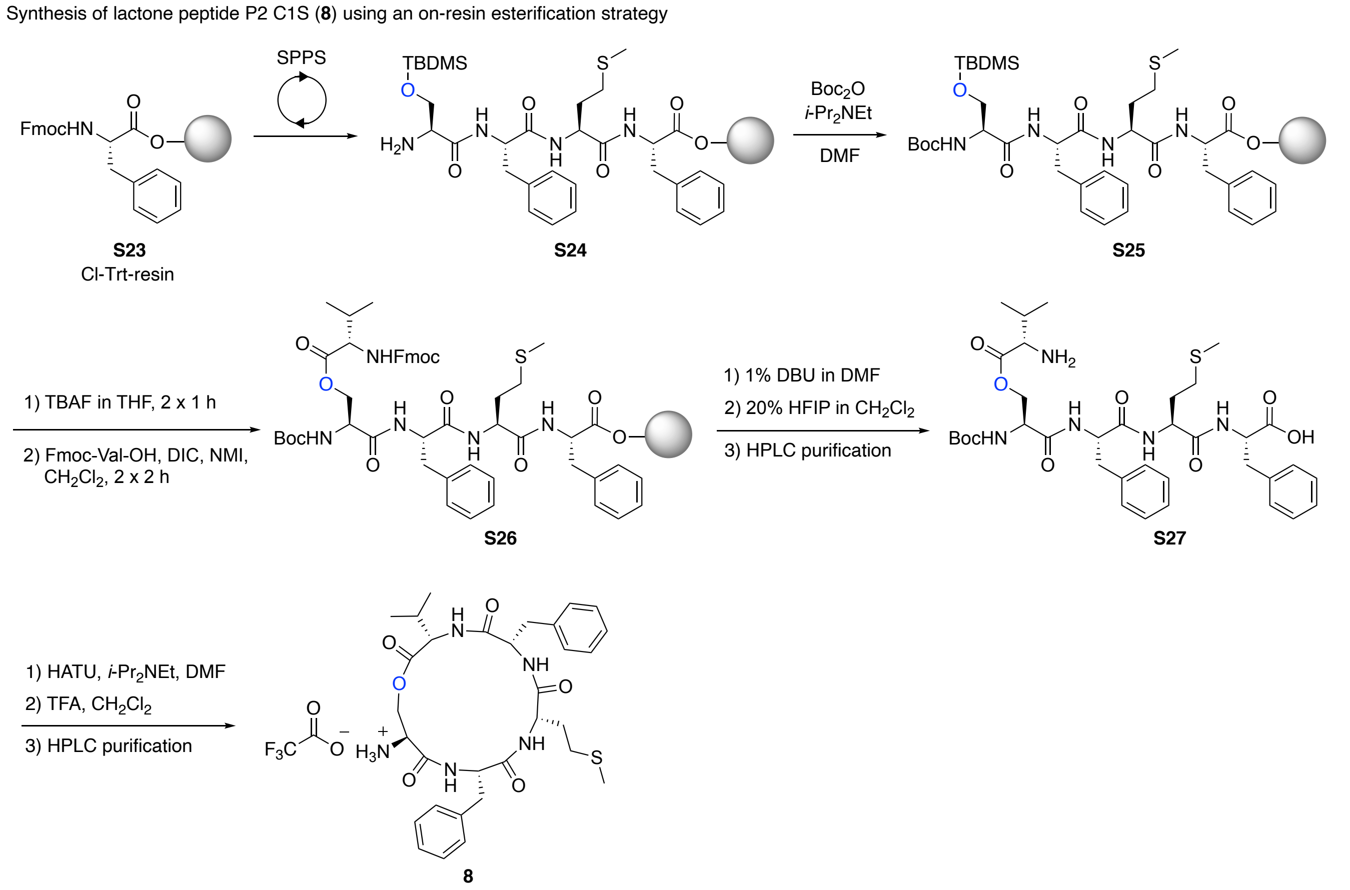

**Supplementary figure 14.** Synthesis of lactone peptide P2 C1Ser (**8**). The partially protected linear peptide **S24** was synthesized on Fmoc-Phe-loaded Cl-Trt-resin **S23** and the last coupling step was performed using Fmoc-Ser(TBDMS)-OH followed by an Fmoc removal step. The *N*-terminus was Boc protected to give fully-protected peptide **S25**, which was treated to with tetrabutylammonium fluoride (TBAF) and subsequently esterified using a previously reported anhydride method[2] to afford resin-bound depsipeptide **S26**. The peptide **S26** was deprotected and released from the solid support to yield the partially protected linear peptide **S27**, which was cyclized via amide bond formation using HATU and treated with TFA to afford the lactone peptide **8**.

### Stability and bioactivity of P2 C1Ser (8)

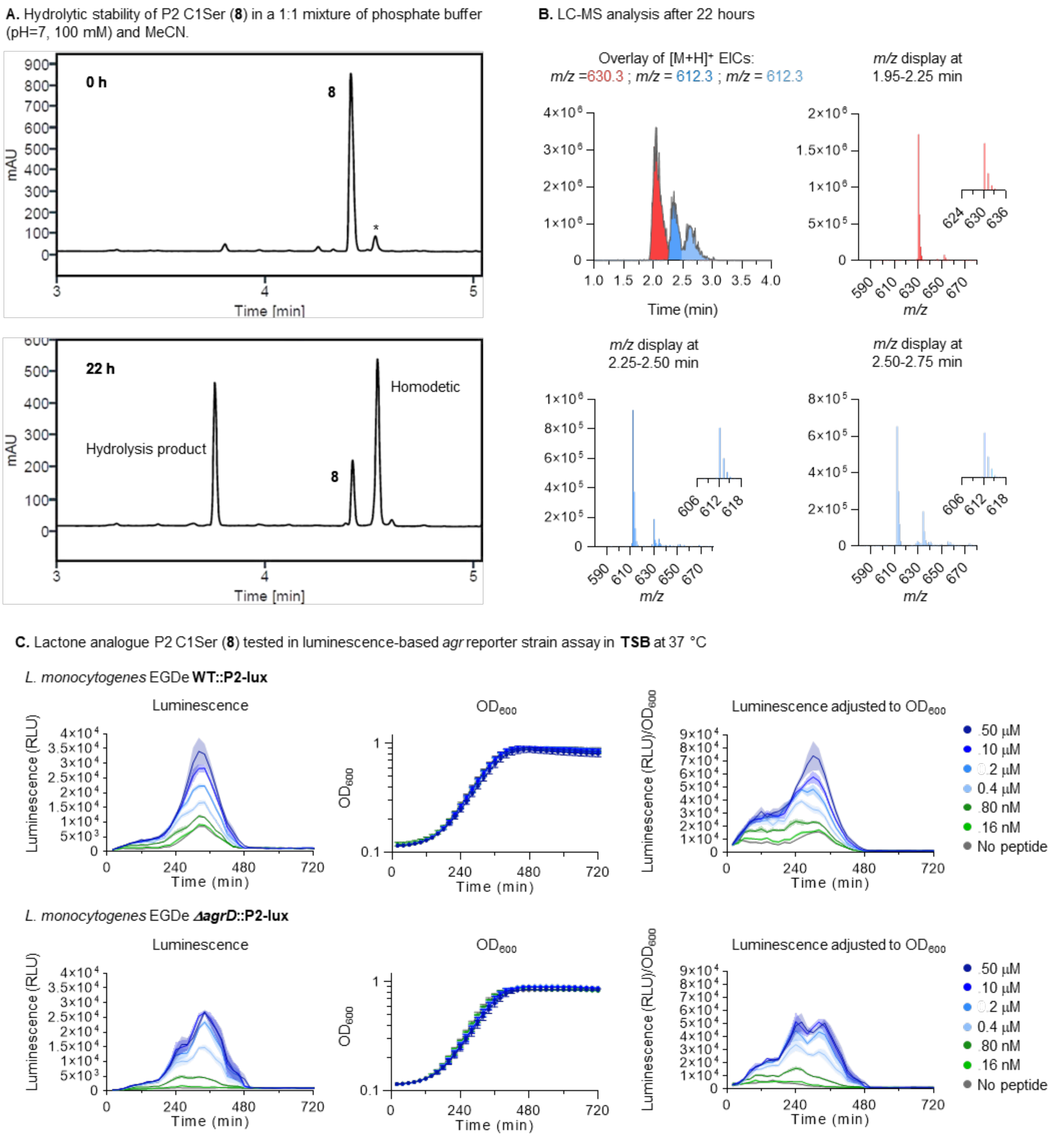

**Supplementary figure 15.** (**A**) Ultra-performance liquid chromatography (UPLC) traces of P2 C1Ser (**8**) at 0 and 22 hours stirred at 37 °C in a 1:1 mixture of phosphate buffer (pH=7, 100 mM) and MeCN, displaying significant hydrolysis, in addition to O→N acyl shift, resulting in the corresponding homodetic peptide form of **8**. *Homodetic peptide formed in the stock. (**B**) Extracted ion chromatogram (EIC) overlay of the $[M+H]^+$ adducts of **8** (*m/z* = 612.3, in blue), homodetic form of **8** (*m/z* = 612.3, in pale blue) and hydrolyzed **8** (*m/z* = 630.3, in red), and mass spectra displaying their presence after 22 hours. (**C**) *L. monocytogenes* EGDe reporter strains treated with synthetic peptide (**8**) at selected concentrations (50-0.016 µM). Graphs represent transcriptional activity from the *agr* promoter measured in relative luminescence units (RLU), bacterial cell density measured as optical density at 600 nm ($OD_{600}$) and *agr* activity adjusted to cell density ($RLU/OD_{600}$). All curves represent the mean of three biological replicates with at least technical duplicates. Error bars and shaded areas represent the standard error of the mean (SEM).

#### Synthesis and bioactivity of thioether-containing peptides 9 and 10

**A.** Synthesis of Fmoc-Val-iodide (**S30**)

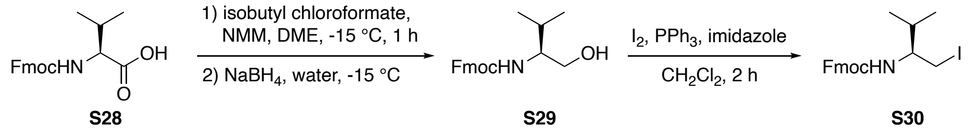

**B.** Synthesis of *N*,*N*-dimethyl thioether Cys-Val building block **S35** for SPPS

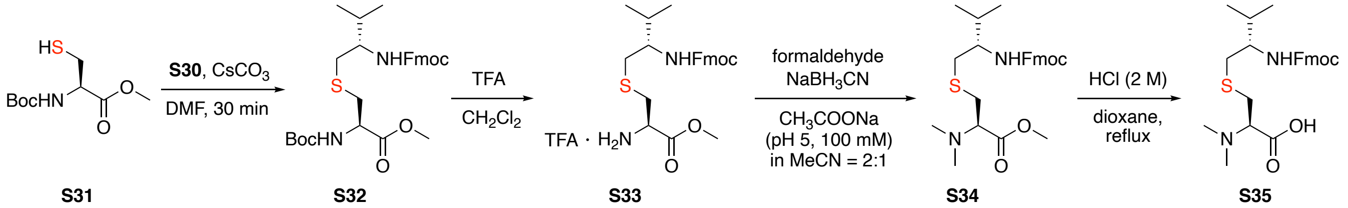

**C.** Synthesis of *N*,*N*-dimethyl P2 thioether (**9**) using a thioether building block strategy

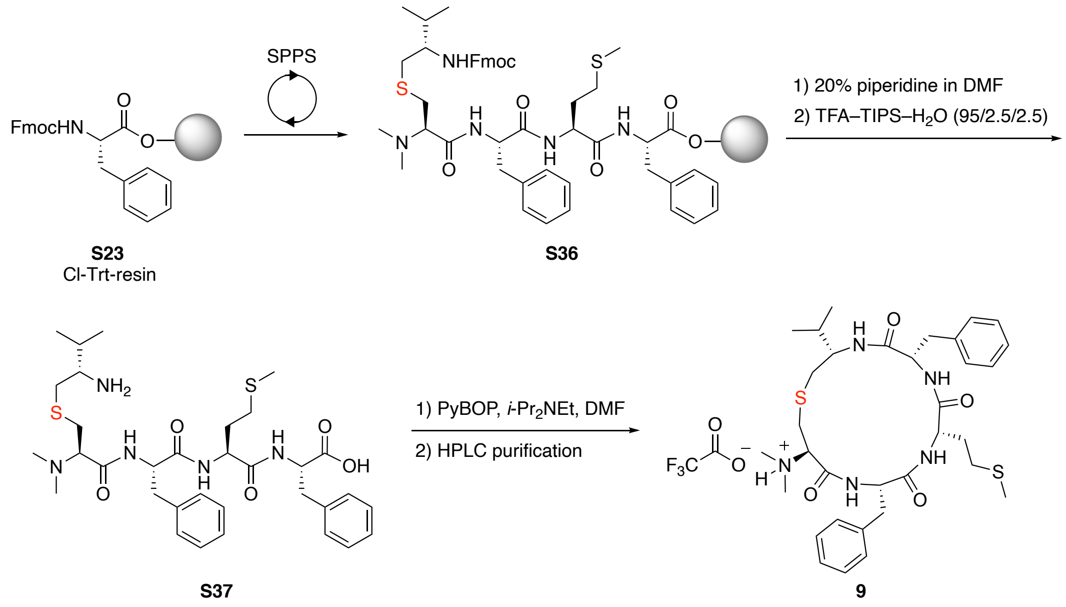

**Supplementary figure 16.** (**A**) Synthesis of Fmoc-Val-iodide (**S30**) starting from Fmoc-Val-OH (**S28**). (**B**) Synthesis of *N,N*-dimethyl-thioether Cys-Val building block **S35** suitable for SPPS. Boc-Cys-OMe (**S31**) was alkylated with Fmoc-Val-iodide (**S30**) to afford **S32**, which was further treated with TFA to yield the TFA salt **S33**. Reductive amination of **S33** gave the *N,N*-dimethyl methyl ester **S34**, which was hydrolyzed under acidic conditions to afford **S35.** (**C**) Synthesis of *N,N*-dimethyl P2 thioether (**9**). The thioether containing protected linear peptide **S36** was synthesized on Fmoc-Phe-loaded Cl-Trt-resin **S23** and the last coupling steps were performed using **S35**. Resin-bound peptide **S36** was deprotected and released from the solid support to yield the linear peptide **S37**, which was cyclized via amide bond formation using PyBOP to afford the *N,N*-dimethyl thioether peptide **9**.

Synthesis of P2 thioether (**10**) via on-resin cysteine alkylation

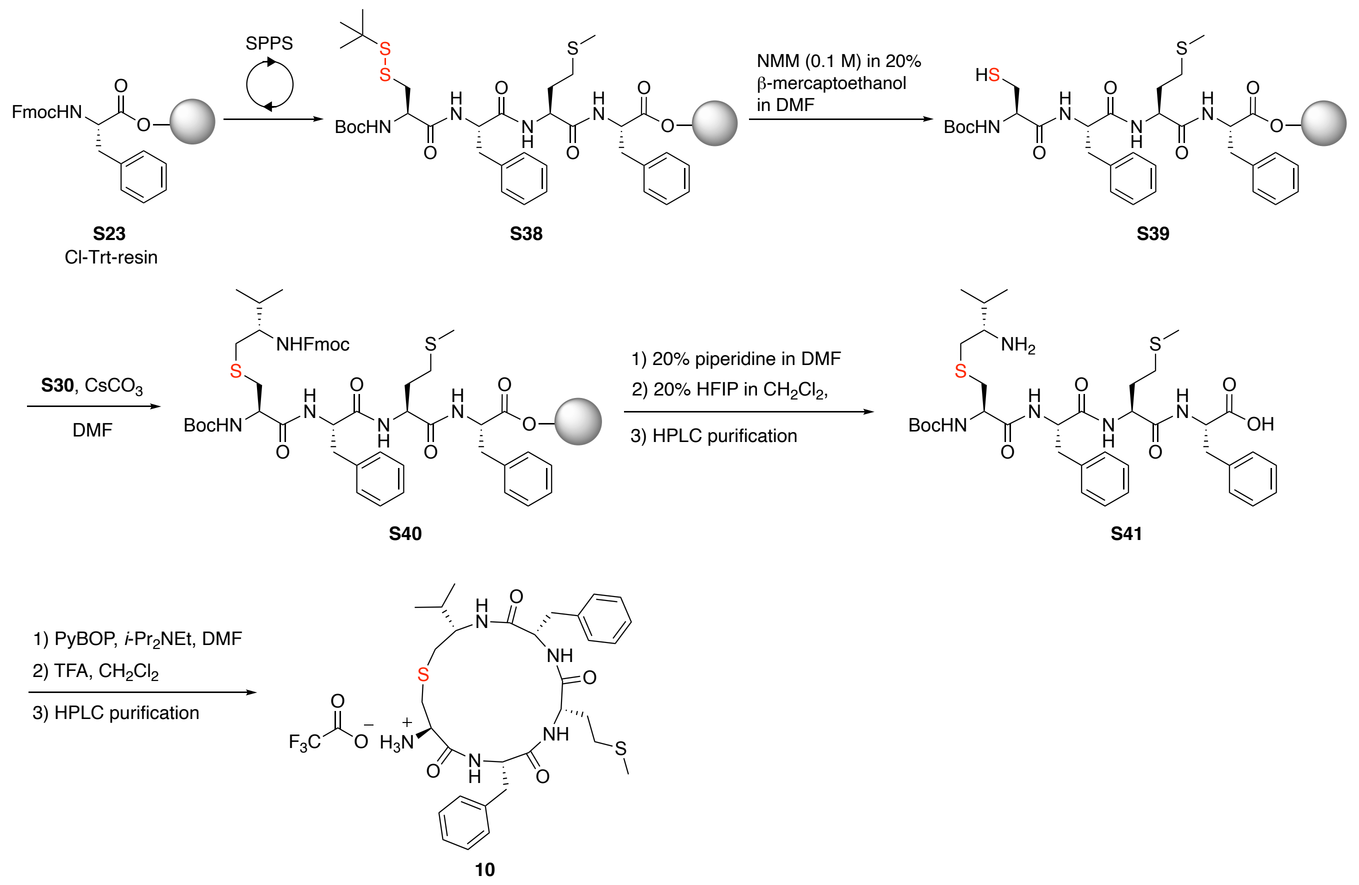

**Supplementary figure 17.** Synthesis of P2 thioether (**10**). The disulfide protected linear peptide **S38** was synthesized on Fmoc-Phe-loaded Cl-Trt-resin **S23** and the cysteine protecting group was removed using β-mercaptoethanol to afford resin-bound peptide **S39**. Cysteine alkylation with **S30** in the presence of $CsCO_3$ gave the thioether-containing pentapeptide **S40**, which was deprotected and released from the solid support to yield the partially protected peptide **S41**. The linear peptide **S41** was cyclized via amide bond formation using PyBOP and treated with TFA to afford the thioether peptide **10**.

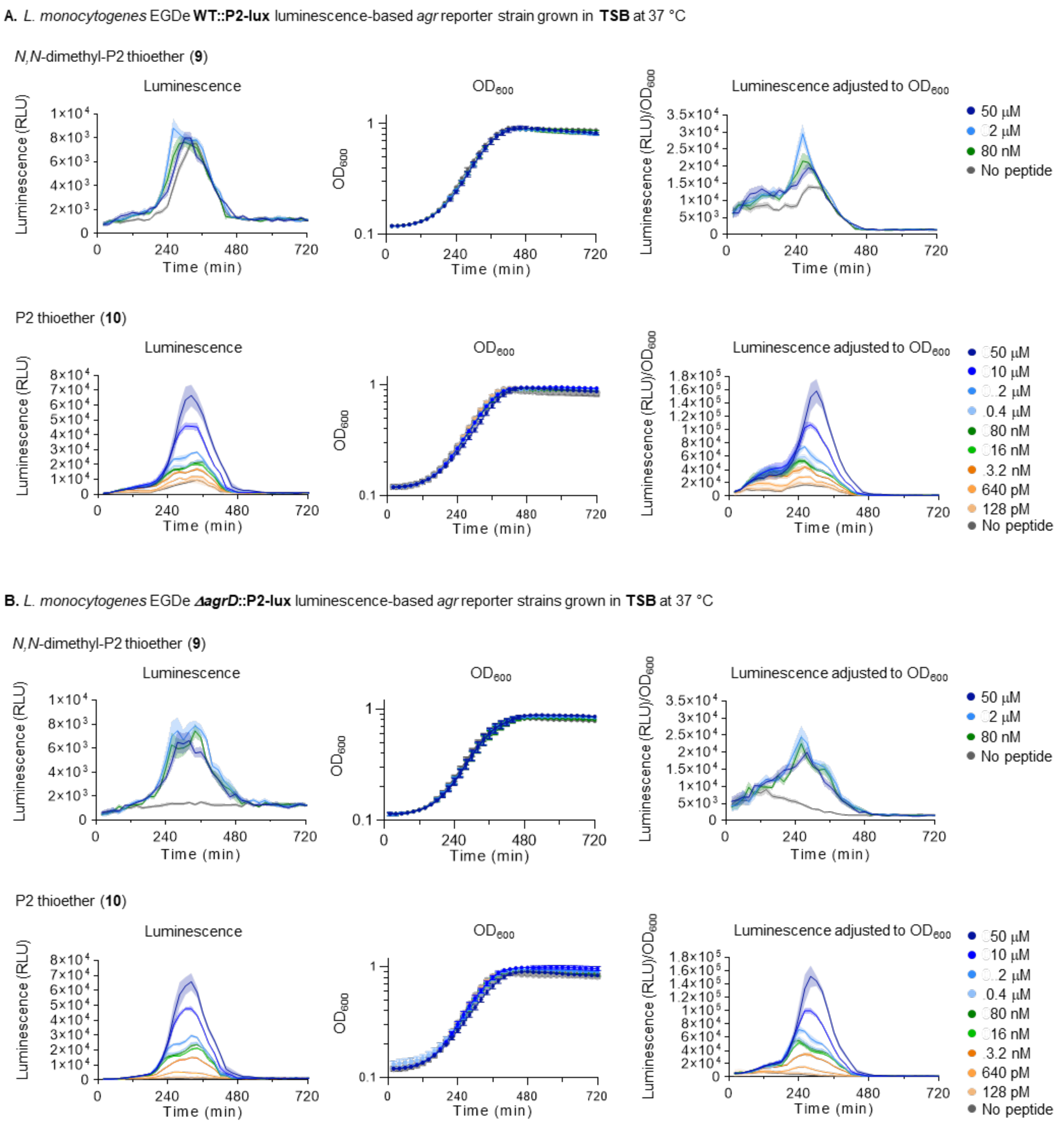

**Supplementary figure 18.** Reporter strain measurements from (**A**) ***L. monocytogenes*** **EGDe WT::P2-lux** and (**B**) ***L. monocytogenes*** **EGDe *ΔagrD*::P2-lux** treated with synthetic peptides (**9** and **10)** at selected concentrations (**50-0.000128 µM**) and grown in tryptic soy broth (**TSB**) medium at **37 °C**. Graphs represent transcriptional activity from the *agr* promoter measured in relative luminescence units (RLU), bacterial cell density measured as optical density at 600 nm ($OD_{600}$) and *agr* activity adjusted to cell density ($RLU/OD_{600}$). All curves represent the mean of three biological replicates with at least technical duplicates. Error bars and shaded areas represent the standard error of the mean (SEM).

### Materials and Methods

#### General Information

##### Reagents and materials

Amino acid building blocks for solid-phase peptide synthesis (SPPS) were purchased from CombiBlocks and 2-(1*H*-benzotriazol-1-yl)-1,1,3,3-tetramethyluronoium hexafluorophosphate (HBTU), 2-(1*H*-7-Azabenzotriazol-1-yl)-1,1,3,3-tetramethyluronoium hexafluorphosphate (HATU) and benzotriazole-1-yl-oxy-tris-pyrrolidino-phosphonium hexafluorophosphate (PyBOP) were purchased from PepChem. Aminomethyl ChemMatrix resin was obtained from PCAS BioMatrix. All other chemicals were obtained in the highest available purity from Merck or CombiBlocks and used without further purification. All solvents used were of analytical grade and purchased from Fisher Scientific. Manual solid-phase peptide synthesis was performed in polypropylene syringes equipped with fritted disks purchased from Torviq. Thermo Scientific™ white 96-well assay plates with clear bottom for luminescence measurements were purchased from Fisher Scientific. Flash column chromatography was performed on silica gel 60 (particle size 35–70 μm) purchased from VWR. Tryptic soy agar (TSA), tryptic soy broth (TSB) and brain heart infusion (BHI) media for bacterial culturing were purchased from Oxoid. Chloramphenicol purchase was purchased from Sigma-Aldrich. Riboflavin 5'-monophosphate sodium salt hydrate (flavin mononucleotide) was purchased from abcr.

##### Analytical ultra-performance liquid chromatography (UPLC)

Analytical UPLC analyses were performed on a C18 Agilent InfinityLab Poroshell 120 column (2.7 μm, 100 × 3.0 mm) using an Agilent 1260 Infinity II series system equipped with a diode array UV detector. Various gradients with eluent A (water–MeCN–TFA, 95:5:0.1, v/v/v) and eluent B (0.1% TFA in MeCN) were applied at a flow rate of 1.2 mL $min^{-1}$ to monitor reactions or to determine the purity of synthesized compounds ($\lambda$ = 215 nm).

##### UPLC mass spectrometry (MS)

UPLC-MS analyses were performed on a Phenomenex Kinetex column (1.7 μm, 100 Å, 50 × 2.10 mm) using a Waters Acquity UPLC system. A gradient with eluent C (0.1% HCOOH in water) and eluent D (0.1% HCOOH in MeCN) rising linearly from 5 to 100% of D over 2.90 min at a flow rate of 0.6 mL $min^{-1}$ was applied to monitor reactions mixtures and to confirm compound identities.

##### Preparative high-performance liquid chromatography (HPLC)

Preparative HPLC purifications were performed on a C18 Phenomenex Luna column (5 μm, 100 Å, 250 × 21.2 mm) or a C8 Phenomenex Luna column (5 μm, 100 Å, 250 × 21.2 mm) using an Agilent 1260 LC system equipped with a diode array UV detector and an evaporative light-scattering detector (ELSD). Various gradients with eluent A (water–MeCN–TFA, 95:5:0.1, v/v/v) and eluent B (0.1% TFA in MeCN) at a flow rate of 20 mL $min^{-1}$ were applied for purification. Fractions containing the purified target compound were identified using UPLC-MS and assessed for purities >95% using analytical UPLC. Selected fractions were pooled and lyophilized. Used gradients for purification are stated in the given experiment.

**Nuclear magnetic resonance (NMR) spectroscopy**

NMR spectra were recorded at 298 K using a Bruker Avance III ($^1$H NMR and $^{13}$C NMR recorded at 400 and 101 MHz, respectively) or a Bruker Avance III HD ($^1$H NMR and $^{13}$C NMR recorded at 600 and 150 MHz, respectively). Chemical shifts are reported in parts per million (ppm) relative to the deuterated solvent peak of DMSO-$d_6$ ($\delta_H$ = 2.50 ppm; $\delta_C$ = 39.52 ppm) or $CDCl_3$ ($\delta_H$ = 7.26 ppm; $\delta_C$ = 77.16 ppm) as internal standard.

**Plate reader**

Luminescence-based reporter strain assay were performed using a BioTek Synergy H1 microplate reader with Gen5™ software. Plates were incubated at 37 °C with continuous double orbital shaking (283 rpm). Luminescence (gain = 255) and optical density at $\lambda$ = 600 nm ($OD_{600}$) was measured every 20 minutes.

**List of used bacterial strains**

| **Strain ID** | **Details** | **Reference** |
|---|---|---|
| *Listeria monocytogenes* EGDe::pPL2*lux*$P_{II}$ (WT::P2-lux) | Chromosomal integration of the *luxABCDEP* operon under regulation of the *agr* promoter into the $tRNA^{Arg}$ locus of *L. monocytogenes* EGDe wild type, $Cm^r$ | [4] |
| *Listeria monocytogenes* EGDe*ΔagrD*::pPL2*lux*$P_{II}$ (*Δagr*D::P2-lux) | Chromosomal integration of the *luxABCDEP* operon under regulation of the *agr* promoter into the $tRNA^{Arg}$ locus of *L. monocytogenes* EGDe*ΔagrD*, $Cm^r$ | [4] |

#### Synthesis of amino acid building blocks

##### *N,N*-Dimethyl-*S*-trityl-*L*-cysteine (S2)

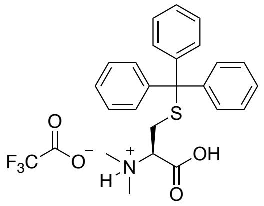

To a solution of H-Cys(Trt)-OH (**S1**) (500 mg, 1.38 mmol, 1.00 equiv) in sodium acetate buffer (pH 5, 100 mM)–MeCN (21 mL, 1:2, v/v) was added 37% formaldehyde solution (1.03 mL, 13.8 mmol, 10.0 equiv) followed by $NaCNBH_3$ (867 mg, 13.8 mmol, 10.0 equiv) and the reaction mixture was stirred at room temperature. After 30 min, the reaction mixture was acidified to pH 3 using HCl (2 M) and extracted with EtOAc (2 × 30 mL). The organic layer was dried over $Na_2SO_4$, filtered and evaporated to dryness under reduced pressure. The crude compound **S2** was purified by flash column chromatography ($CH_2Cl_2$–MeOH = 9:1 + 0.25% AcOH) to afford the acetate salt of **S2**. To enable the use in SPPS, the acetate salt of **S2** was lyophilized in water containing 0.1% TFA to afford the trifluoroacetate salt of **S2** (260 mg, 0.51 mmol, 37%). **$^1$H NMR** (600 MHz, $CDCl_3$) δ = 7.37–7.28 (m, 6H), 7.25–7.15 (m, 6H), 7.15–7.10 (m, 3H), 3.12 (d, *J* = 6.7 Hz, 1H), 2.98 (dd, *J* = 13.0, 7.5 Hz, 1H), 2.33 (s, 6H), 2.23 (dd, *J* = 13.1, 5.8 Hz, 1H). **$^{13}$C NMR** (151 MHz, $CDCl_3$) δ = 169.2, 144.4, 129.8, 128.2, 127.0, 69.4, 67.9, 41.1, 28.0. **UPLC-MS** (ESI) *m/z* calcd for $[M+H]^+$ $C_{24}H_{26}NO_2S^+$: 392.17, found 392.21.

##### *(S)*-3-((((9*H*-Fluoren-9-yl)methoxy)carbonyl)amino)-2-(dimethylamino)propanoic acid (S16)

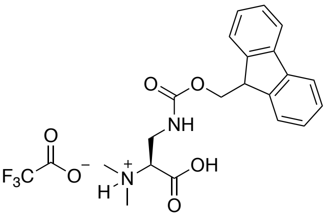

Boc-Dap(Fmoc)-OH (**S14**) (1.00 g, 2.34 mmol, 1.00 equiv) was dissolved in $CH_2Cl_2$ (5.0 mL) and TFA (5.0 mL) was added to the solution and the reaction mixture was stirred at room temperature. After 30 min, the solvents were removed under reduced pressure and crude product **S15** was co-evaporated three times with toluene to remove excess TFA affording trifluoroacetate salt **S15** (1.03 g, 2.34 mmol, quant.) which was used without further purification. Trifluoroacetate salt **S15** (150 mg, 0.32 mmol, 1.00 equiv) was dissolved in sodium acetate buffer (pH 5, 100 mM)–MeCN (5 mL, 1:2, v/v) and 37% formaldehyde solution (171 µL, 2.24 mmol, 7.00 equiv) followed by $NaCNBH_3$ (141 mg, 2.24 mmol, 7.00 equiv) was added the solution and the reaction mixture was stirred at room temperature. After 30 min, the reaction mixture was purified by reverse-phase automated column chromatography to afford the title compound **S16** as trifluoroacetate salt (144 mg, 0.31 mmol, 96%) after lyophilization. **$^1$H NMR** (600 MHz, DMSO-*d*$_6$) δ = 7.89 (d, *J* = 7.5 Hz, 2H), 7.75–7.62 (m, 3H), 7.42 (td, *J* = 7.5, 1.1 Hz, 2H), 7.33 (td, *J* = 7.4, 1.1 Hz, 2H), 4.39–4.33 (m, 2H), 4.23 (t, *J* = 6.7 Hz, 1H), 4.10 (t, *J* = 5.4 Hz, 1H), 3.65 (t, *J* = 5.8 Hz, 2H), 2.85 (s, 6H). **$^{13}$C NMR** (151 MHz, DMSO-*d*$_6$) δ = 168.1, 156.4, 143.8, 143.8, 140.8, 140.8, 127.7, 127.2, 125.2, 120.2, 65.9, 65.8, 46.7, 41.5, 38.3. **UPLC-MS** (ESI) *m/z* calcd for $[M+H]^+$ $C_{20}H_{23}N_2O_4^+$: 355.17, found 355.09.

**(9*H*-Fluoren-9-yl)methyl (*S*)-(1-iodo-3-methylbutan-2-yl)carbamate (S30)**

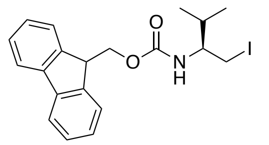

$C_{20}H_{22}INO_2$
MW: 435.31

To a solution of Fmoc-Val-OH (**S28**) (2.00 g, 5.89 mmol, 1.00 equiv) in 1,2-dimethoxyethane (DME) (12.0 mL) cooled to -15 °C using a salt/ice bath under nitrogen atmosphere was added *N*-methylmorpholine (NMM) (0.65 mL, 5.89 mmol, 1.00 equiv) followed by isobutyl chloroformate (0.77 mL, 5.89 mmol, 1.00 equiv) and the reaction mixture was stirred at -15 °C. After 1 h, the reaction mixture was filtered and the filtrate was cooled to -15 °C before a solution of $NaBH_4$ (334 mg, 8.84 mmol, 1.50 equiv) in water (3.0 mL) was added. After 5 min, the reaction mixture was diluted with water (150 mL) leading to precipitation of the desired Fmoc-Val alcohol **S29** (1.60 g, 4.92 mmol, 84%), which was collected by filtration and used without further purification.

Fmoc-Val iodide (**S30**) was prepared according to a previously published procedure.[3] $PPh_3$ (1.28 g, 4.90 mmol, 3.00 equiv), iodine (1.24 g, 4.90 mmol, 3.00 equiv) and imidazole (554 mg, 8.15 mmol, 5.00 equiv) were dissolved in anhydrous $CH_2Cl_2$ (16.0 mL) under nitrogen atmosphere and stirred for 5 min before a solution of Fmoc-Val alcohol (530 mg, 163 mmol, 1.00 equiv) in anhydrous $CH_2Cl_2$ (4.0 mL) was added and the reaction mixture was stirred at room temperature. After 1.5 h, the solvent was removed under reduced pressure and the remaining crude residue purified by flash column chromatography (heptane–EtOAc = 9:1) to afford the title compound **S30** (557 mg, 1.28 mmol, 75%). **$^1$H NMR** (600 MHz, DMSO-*d*$_6$) δ = 7.89 (d, *J* = 7.5 Hz, 2H), 7.73 (dd, *J* = 10.2, 7.5 Hz, 2H), 7.44–7.36 (m, 3H), 7.36–7.28 (m, 2H), 4.41–4.28 (m, 2H), 4.23 (t, *J* = 7.1 Hz, 1H), 3.43 (dd, *J* = 10.1, 3.7 Hz, 1H), 3.39–3.28 (m, 1H), 3.23 (dd, *J* = 10.0, 8.5 Hz, 1H), 1.82–1.73 (m, *J* = 6.8 Hz, 1H), 0.85 (t, *J* = 6.2 Hz, 6H). %). **$^{13}$C NMR** (151 MHz, DMSO-*d*$_6$) δ = 155.9, 143.9, 143.8, 140.7, 127.6, 127.0, 127.0, 125.3, 125.2, 120.1, 65.2, 57.7, 46.8, 31.8, 19.2, 18.0, 11.7.

***S*-((*S*)-2-((((9*H*-Fluoren-9-yl)methoxy)carbonyl)amino)-3-methylbutyl)-*N*,*N*-dimethyl-*L*-cysteine (S35)**

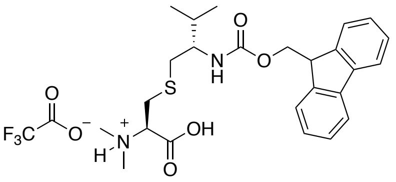

$C_{25}H_{32}N_2O_4S$
MW: 456.60
TFA salt: 570.62

A suspension of $CsCO_3$ (113 mg, 0.35 mmol, 1.50 equiv) in anhydrous DMF (2.0 mL) was sonicated for 15 min and subsequently Boc-Cys-OMe (**S31**) (81.0 mg, 0.35 mmol, 1.50 equiv) and Fmoc-Val iodide (**S30**) (100 mg, 0.23 mmol, 1.00 equiv) were added and the reaction mixture stirred at room temperature. After 30 min, the reaction mixture was diluted with water (10 mL) and extracted with EtOAc (3 × 10 mL). The organic layer was dried over $Na_2SO_4$, filtered and evaporated to dryness under reduced pressure to yield the crude product **S32**, which was dissolved in TFA–$CH_2Cl_2$ (5.0 mL, 1:1, v/v) and stirred at room temperature. After 30 min, the solvents were removed under reduced pressure and crude product **S33** was co-evaporated three times with toluene to remove excess TFA affording trifluoroacetate salt **S33**, which was used without further purification. Trifluoroacetate salt **S33** was dissolved in sodium acetate buffer (pH 5, 100 mM)–MeCN (15 mL, 1:2, v/v) and 37% formaldehyde solution (344 µL, 4.60 mmol, 20.0 equiv) followed by $NaCNBH_3$ (290 mg, 4.60 mmol, 20.0 equiv) was added the solution and the reaction mixture was stirred at room temperature. After 30 min, MeCN was removed under reduced pressure and the reaction mixture was extracted with EtOAc (3 × 10 mL). The organic layer was dried over $Na_2SO_4$, filtered and evaporated to dryness under reduced pressure to yield the crude dimethyl product **S34**, which was dissolved in dioxane–HCl (2 M) (5.0 mL, 1:1, v/v) and heated under reflux for 48 h. The reaction mixture was purified by reverse-phase automated column chromatography to afford the title compound **S35** as trifluoroacetate salt (74.2 mg, 0.13 mmol, 57%) after lyophilization. **$^1$H NMR** (600 MHz, DMSO-$d_6$) δ = 7.89 (d, $J$ = 7.6 Hz, 2H), 7.71 (t, $J$ = 7.0 Hz, 3H), 7.42 (t, $J$ = 7.4 Hz, 2H), 7.33 (td, $J$ = 7.5, 3.0 Hz, 2H), 7.22 (d, $J$ = 9.3 Hz, 1H), 4.38–4.32 (m, 1H), 4.30–4.20 (m, 3H), 3.56–3.40 (m, 1H), 3.19–3.04 (m, 2H), 2.85–2.77 (m, 7H), 2.57 (dd, $J$ = 13.3, 10.0 Hz, 1H), 1.74 (tq, $J$ = 13.3, 6.6 Hz, 1H), 0.88–0.82 (m, 6H). **$^{13}$C NMR** (151 MHz, DMSO-$d_6$) δ = 168.3, 156.3, 144.0, 143.8, 140.7, 127.6, 127.0, 127.0, 125.2, 120.1, 120.1, 65.9, 65.3, 55.4, 46.8, 41.3, 34.9, 31.6, 28.6, 19.3, 17.9. **UPLC-MS** (ESI) *m/z* calcd for $[M+H]^+$ $C_{25}H_{33}N_2O_4S^+$: 457.22, found 457.21.

#### Characterization data for synthetic peptides

##### *N,N*-Dimethyl-P2 (1)

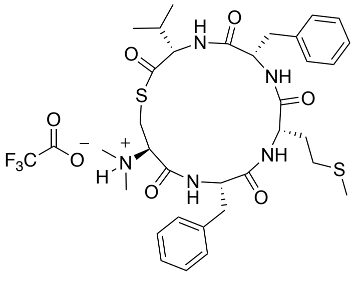

$C_{33}H_{45}N_5O_5S_2$
MW: 655.87
TFA salt: 769.90

Peptide **1** was synthesized on MeDbz-resin on 20 µmol scale using the general procedures for automated SPPS, manual coupling of *N,N*-dimethyl-Cys(Trt)-OH (**S2**) and the synthesis of *N,N*-dimethyl-thiolactone peptides. Preparative RP-HPLC purification (5–95% B over 30 min) afforded the trifluoroacetate salt of peptide **1** as a white solid (4.7 mg, 6.1 µmol, 31%). Purity 95% determined by UPLC ($\lambda$ = 215 nm). **$^1$H NMR** (600 MHz, DMSO-$d_6$) δ = 10.51 (br s, 1H), 8.92 (d, *J* = 7.7 Hz, 1H), 8.77 (d, *J* = 9.1 Hz, 1H), 8.35 (d, *J* = 9.9 Hz, 1H), 7.31–7.15 (m, 11H), 4.69–4.62 (m, 1H), 4.60 (dd, *J* = 9.9, 4.4 Hz, 1H), 4.39 (ddd, *J* = 12.3, 8.3, 4.3 Hz, 1H), 3.76 (td, *J* = 8.2, 6.1 Hz, 1H), 3.72–3.68 (m, 1H), 3.47 (dd, *J* = 12.9, 4.3 Hz, 1H), 3.30–3.15 (m, 3H), 3.10 (dd, *J* = 13.6, 5.1 Hz, 1H), 2.82 (dd, *J* = 13.6, 10.9 Hz, 1H), 2.77 (dd, *J* = 12.9, 11.7 Hz, 1H), 2.69 (s, 3H), 2.43 (pd, *J* = 6.9, 4.5 Hz, 1H), 2.34 (s, 3H), 2.18–2.10 (m, 2H), 1.98 (s, 0H), 1.96 (s, 3H), 1.94–1.81 (m, 2H), 0.99 (d, *J* = 6.9 Hz, 3H), 0.95 (d, *J* = 6.8 Hz, 3H). **UPLC-MS** (ESI) *m/z* calcd for $[M+H]^+$ $C_{33}H_{46}N_5O_5S_2^+$: 656.29, found 656.35.

##### *N,N*-Dimethyl-P2 F2A (2)

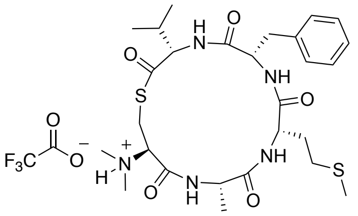

$C_{27}H_{41}N_5O_5S_2$
MW: 579.78
TFA salt: 693.80

Peptide **2** was synthesized on MeDbz-resin on 20 µmol scale using the general procedures for automated SPPS, manual coupling of *N,N*-dimethyl-Cys(Trt)-OH (**S2**) and the synthesis of *N,N*-dimethyl-thiolactone peptides. Preparative RP-HPLC purification (5–95% B over 30 min) afforded the trifluoroacetate salt of peptide **2** as a white solid (4.3 mg, 6.2 µmol, 31%). Purity 97% determined by UPLC ($\lambda$ = 215 nm). **$^1$H NMR** (600 MHz, DMSO-$d_6$) δ = 10.49 (s, 1H), 8.64–8.38 (m, 3H), 8.24 (d, *J* = 9.9 Hz, 1H), 7.57–7.04 (m, 5H), 4.58 (dd, *J* = 9.9, 4.2 Hz, 1H), 4.39 (p, *J* = 7.4 Hz, 1H), 4.34–4.21 (m, 1H), 3.88–3.75 (m, 2H), 3.67 (q, *J* = 7.3 Hz, 1H), 3.48 (dd, *J* = 13.0, 4.5 Hz, 1H), 3.29–3.16 (m, 2H), 2.92–2.84 (m, 1H), 2.79 (s, 4H), 2.41 (pt, *J* = 6.6, 3.3 Hz, 1H), 2.24 (t, *J* = 7.5 Hz, 2H), 1.98 (s, 4H), 1.92–1.83 (m, 1H), 1.24 (d, *J* = 7.2 Hz, 3H), 0.94 (t, *J* = 6.7 Hz, 6H). **UPLC-MS** (ESI) *m/z* calcd for $[M+H]^+$ $C_{27}H_{42}N_5O_5S_2^+$: 580.26, found 580.21.

##### *N,N*-Dimethyl-P2 M3A (3)

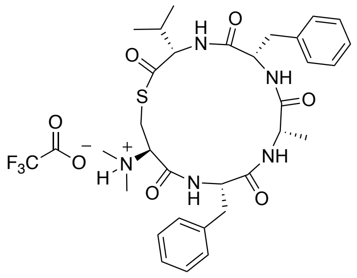

$C_{31}H_{41}N_5O_5S$
MW: 595.76
TFA salt: 709.78

Peptide **3** was synthesized on MeDbz-resin on 20 μmol scale using the general procedures for automated SPPS, manual coupling of *N,N*-dimethyl-Cys(Trt)-OH (**S2**) and the synthesis of *N,N*-dimethyl-thiolactone peptides. Preparative RP-HPLC purification (5–95% B over 30 min) afforded the trifluoroacetate salt of peptide **3** as a white solid (3.8 mg, 5.4 μmol, 27%). Purity 98% determined by UPLC ($\lambda$ = 215 nm). **$^1$H NMR** (600 MHz, DMSO-$d_6$) δ = 10.30 (s, 1H), 8.74 (s, 1H), 8.49 (d, *J* = 9.3 Hz, 1H), 8.41 (d, *J* = 8.0 Hz, 1H), 8.24 (d, *J* = 10.0 Hz, 1H), 7.38–7.06 (m, 10H), 4.69–4.64 (m, 1H), 4.62 (dd, *J* = 10.0, 3.9 Hz, 1H), 4.22 (q, *J* = 7.8 Hz, 1H), 3.74 (s, 1H), 3.71–3.63 (m, 0H), 3.46–3.41 (m, 9H), 3.26 (d, *J* = 7.8 Hz, 2H), 3.02 (dd, *J* = 13.6, 4.5 Hz, 1H), 2.78–2.66 (m, 2H), 2.48–2.40 (m, 1H), 1.26 (d, *J* = 6.9 Hz, 3H), 0.98 (d, *J* = 6.9 Hz, 3H), 0.94 (d, *J* = 6.9 Hz, 3H). **UPLC-MS** (ESI) *m/z* calcd for $[M+H]^+$ $C_{31}H_{42}N_5O_5S^+$: 596.29, found 596.23.

##### *N,N*-Dimethyl-P2 F4A (4)

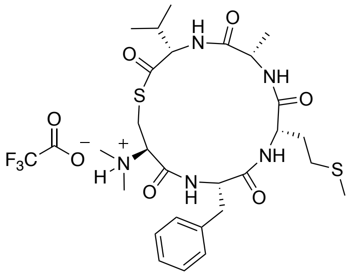

$C_{27}H_{41}N_5O_5S_2$
MW: 579.78
TFA salt: 693.80

Peptide **4** was synthesized on MeDbz-resin on 20 μmol scale using the general procedures for automated SPPS, manual coupling of *N,N*-dimethyl-Cys(Trt)-OH (**S2**) and the synthesis of *N,N*-dimethyl-thiolactone peptides. Preparative RP-HPLC purification (5–95% B over 30 min) afforded the trifluoroacetate salt of peptide **4** as a white solid (1.5 mg, 2.2 μmol, 11%). Purity 91% determined by UPLC ($\lambda$ = 215 nm). **$^1$H NMR** (600 MHz, DMSO-$d_6$) δ = 10.36 (s, 1H), 9.03 (s, 1H), 8.56 (d, *J* = 9.5 Hz, 1H), 8.42 (d, *J* = 7.8 Hz, 1H), 8.09 (d, *J* = 9.9 Hz, 1H), 7.33–7.16 (m, 5H), 4.74 (ddd, *J* = 11.2, 9.5, 4.6 Hz, 1H), 4.53 (dd, *J* = 9.9, 4.2 Hz, 1H), 4.19 (p, *J* = 7.4 Hz, 1H), 3.86–3.79 (m, 1H), 3.73 (s, 1H), 3.43 (dd, *J* = 12.9, 4.6 Hz, 1H), 3.09 (dd, *J* = 13.6, 4.6 Hz, 1H), 2.73 (ddd, *J* = 13.0, 11.4, 2.7 Hz, 2H), 2.47–2.29 (m, 3H), 2.19–2.09 (m, 1H), 2.05 (s, 4H), 1.43 (d, *J* = 7.3 Hz, 3H), 0.96 (d, *J* = 6.9 Hz, 3H), 0.91 (d, *J* = 6.8 Hz, 3H). **UPLC-MS** (ESI) *m/z* calcd for $[M+H]^+$ $C_{27}H_{42}N_5O_5S_2^+$: 580.26, found 580.24.

***N,N*-Dimethyl-P2 V5A (5)**

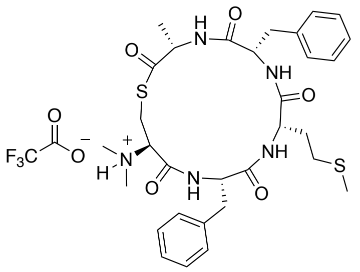

$C_{31}H_{41}N_5O_5S_2$
MW: 627.82
TFA salt: 741.84

Peptide **5** was synthesized on MeDbz-resin on 20 μmol scale using the general procedures for automated SPPS, manual coupling of *N,N*-dimethyl-Cys(Trt)-OH (**S2**) and the synthesis of *N,N*-dimethyl-thiolactone peptides. Preparative RP-HPLC purification (5–95% B over 30 min) afforded the trifluoroacetate salt of peptide **5** as a white solid (5.1 mg, 6.9 μmol, 35%). Purity 98% determined by UPLC ($\lambda$ = 215 nm). **$^1$H NMR** (600 MHz, DMSO-$d_6$) δ = 9.00 (d, $J$ = 7.6 Hz, 1H (d, $J$ = 9.3 Hz, 1H), 8.40 (d, $J$ = 9.1 Hz, 1H), 8.33 (d, $J$ = 8.4 Hz, 1H), 7.38–7.05 (m, 10H), 4.68 (td, $J$ = 9.9, 5.4 Hz, 1H), 4.61–4.53 (m, 1H), 4.40 (ddd, $J$ = 12.1, 8.4, 4.4 Hz, 1H), 3.71–3.64 (m, 1H), 3.62 (dd, $J$ = 11.6, 4.3 Hz, 1H), 3.44–3.38 (m, 10H), 3.18 (dd, $J$ = 13.6, 4.4 Hz, 1H), 3.09–3.00 (m, 2H), 2.84–2.71 (m, 2H), 2.14 (t, $J$ = 7.3 Hz, 2H), 1.96 (s, 4H), 1.95–1.87 (m, 1H), 1.37 (d, $J$ = 7.0 Hz, 3H). **UPLC-MS** (ESI) *m/z* calcd for $[M+H]^+$ $C_{31}H_{42}N_5O_5S_2^+$: 628.26, found 628.24.

***N,N*-Dimethyl-P2 C1Dap (6)**

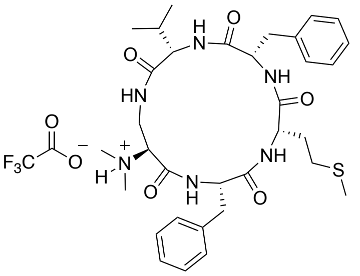

$C_{33}H_{46}N_6O_5S$
MW: 638.83
TFA salt: 752.85

The protected linear peptide **S18** was synthesized in 40.0 μmol scale on Cl-Trt polystyrene resin preloaded with Fmoc-Val-OH **S17** (0.90 mmol/g) using the general procedures for automated SPPS and manual coupling of *N,N*-dimethyl-Dap(Fmoc)-OH (**S16**). After completed peptide elongation, a solution of piperidine in DMF (2.0 mL, 1:4, v/v) was added to the resin and the resin agitated at room temperature for 15 min. The resin was subsequently washed with DMF (3 × 1 min), $CH_2Cl_2$ (3 × 1 min) and dried under suction for 15 min. The dried resin was treated with a cleavage cocktail (3.0 mL, TFA–*i*-$Pr_3SiH$–water, 95:2.5:2.5, v/v/v) for 30 min at room temperature. The cleavage solution was removed from the resin, collected and the resin rinsed with TFA (1.0 mL). The combined cleavage solution and the rinsing solution were concentrated under a stream of nitrogen and the cleaved peptide **S19** was triturated in cold $Et_2O$ (10 mL) and pelleted by centrifugation. The crude peptide **S19** (27 mg, 30.5 μmol) was obtained as TFA salt and used without further purification.

The linear peptide **S19** (27 mg, 30.5 μmol, 1.00 equiv) was dissolved in anhydrous DMF (3.0 mL) under nitrogen atmosphere and added dropwise to a solution of PyBOP (15.9 mg, 30.5 μmol, 1.00 equiv) and *i*-$Pr_2NEt$ (21.2 μL, 122 μmol, 4.00 equiv) in anhydrous DMF (28.0 mL). The reaction mixture was stirred overnight at room temperature and after full consumption of **S19** was confirmed by UPLC-MS, the reaction was reduced to dryness under reduced pressure. The remaining residue was purified by preparative RP-HPLC (20–70% B over 30 min) to afford the trifluoroacetate salt of peptide **6** as a white solid (9.0 mg, 12.0 µmol, 39% from peptide **S22**). Purity 98% determined by UPLC ($\lambda$ = 215 nm). **$^1$H NMR** (600 MHz, DMSO-$d_6$) δ = 10.03 (s, 1H), 9.20 (d, *J* = 6.8 Hz, 1H), 8.56 (d, *J* = 8.0 Hz, 1H), 8.01 (d, *J* = 7.4 Hz, 1H), 7.59 (d, *J* = 9.8 Hz, 1H), 7.31–7.17 (m, 11H), 4.67–4.60 (m, 1H), 4.29 (dd, *J* = 9.8, 5.1 Hz, 1H), 4.01 (ddd, *J* = 11.5, 7.3, 4.1 Hz, 1H), 3.94–3.86 (m, 2H), 3.55–3.50 (m, 1H), 3.38–3.19 (m, 3H), 3.08 (dd, *J* = 13.8, 5.3 Hz, 1H), 2.80–2.71 (m, 2H), 2.71–2.55 (broad signal for $(CH_3)_2N^+$, 6H), 2.38–2.30 (m, 1H), 2.29–2.21 (m, 2H), 2.02–1.98 (m, 4H), 1.97–1.88 (m, 1H), 0.96 (d, *J* = 6.9 Hz, 3H), 0.93 (d, *J* = 6.8 Hz, 3H). *The final peptide sample contains tri(pyrrolidin-1-yl)phosphine oxide as non-UV active (215–280 nm) impurity (340 µg in 9.0 mg, ~10 mol%) from the PyBOP cyclization step, which was inseparable by RP-HPLC. **UPLC-MS** (ESI) *m/z* calcd for $[M+H]^+$ $C_{33}H_{47}N_6O_5S^+$: 639.33, found 639.30.

**P2 C1Dap (7)**

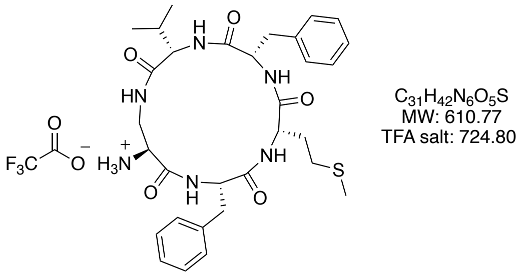

The protected linear peptide **S20** was synthesized in 80.0 µmol scale on Cl-Trt polystyrene resin preloaded with Fmoc-Val-OH **S17** (0.90 mmol/g) using the general procedures for automated SPPS and manual coupling of Fmoc-Dap(Alloc)-OH. After completed peptide elongation, a solution of di-*tert*-butyl dicarbonate ($Boc_2O$) (349 mg, 1.60 mmol, 20.0 equiv), *i*-$Pr_2NEt$ (0.42 mL, 2.40 mmol, 30.0 equiv) in DMF (5.0 mL) was added the peptidyl-resin **S20** with free *N*-terminal amine and the resin was agitated at room temperature. After 1 h, the solution was removed and the Boc-protected peptidyl-resin **S21** was washed with DMF (3 × 1 min) and $CH_2Cl_2$ (3 × 1 min) and dried under high vacuum. The resin **S21** was swelled in anhydrous $CH_2Cl_2$ (3.0 mL) for 15 min and subsequently a solution of $Pd(PPh_3)_4$ (18.5 mg, 16.0 µmol, 0.20 equiv), $Me_2NH·BH_3$ (23.6 mg, 0.40 mmol, 5.00 equiv) in anhydrous $CH_2Cl_2$ (3.0 mL) was added to the resin and the resin was agitated at room temperature. After 15 min, the solution was removed by suction and the resin treated with a fresh $Pd(PPh_3)_4$–$Me_2NH·BH_3$–$CH_2Cl_2$ solution. After 15 min, the solution was removed by suction and the resin washed with DMF (3 × 1 min) and $CH_2Cl_2$ (3 × 1 min) and dried overnight under vacuum. The dried resin was treated with a solution of (hexafluoroisopropanol) HFIP in $CH_2Cl_2$ (5.0 mL, 1:4, v/v) for 30 min at room temperature. The cleavage solution was removed from the resin, collected and a fresh HFIP–$CH_2Cl_2$ solution was added to the resin. After 30 min the cleavage solution was removed from the resin, collected and the resin rinsed with $CH_2Cl_2$ (5.0 mL). The combined cleavage solutions and the rinsing solution were evaporated to dryness under reduced pressure to yield the crude peptide **S22,** which was purified by preparative RP-HPLC (5–95% B over 30 min) to afford the trifluoroacetate salt of peptide **S22** as a white solid (24 mg, 28.5 µmol). The partially-protected peptide **S22** (15.0 mg, 17.8 µmol, 1.00 equiv) was dissolved in anhydrous DMF (3.0 mL) under nitrogen atmosphere and added dropwise to a solution of HATU (6.77 mg, 17.8 µmol, 1.00 equiv) and *i*-$Pr_2NEt$

(12.4 µL, 71.2 µmol, 4.00 equiv) in anhydrous DMF (15.0 mL). The reaction mixture was stirred overnight at room temperature and after full consumption of **S22** was confirmed by UPLC-MS, the reaction was reduced to dryness under reduced pressure. The remaining residue was treated with a solution of TFA in $CH_2Cl_2$ (4.0 mL, 1:1, v/v) for 1 h and subsequently concentrated under a stream of nitrogen and purified by preparative RP-HPLC (5–95% B over 30 min) to afford the trifluoroacetate salt of peptide **7** as a white solid (3.2 mg, 4.4 µmol, 25% from peptide **S22**). Purity 98% determined by UPLC ($\lambda$ = 215 nm). **$^1$H NMR** (600 MHz, DMSO-*d*$_6$) δ = 9.13 (d, *J* = 6.7 Hz, 1H), 8.47 (d, *J* = 7.0 Hz, 1H), 8.15 (s, 3H), 7.98 (d, *J* = 7.3 Hz, 1H), 7.61 (d, *J* = 9.6 Hz, 1H), 7.36–7.16 (m, 10H), 7.10 (t, *J* = 6.1 Hz, 1H), 4.56–4.48 (m, 1H), 4.25 (dd, *J* = 9.6, 5.4 Hz, 1H), 4.05–3.98 (m, 1H), 3.93–3.88 (m, 1H), 3.87–3.79 (m, 1H), 3.55–3.49 (m, 1H), 3.39–3.33 (m, 1H), 3.31–3.20 (m, 2H), 3.02 (dd, *J* = 14.0, 5.9 Hz, 1H), 2.84 (dd, *J* = 14.0, 9.3 Hz, 1H), 2.32–2.24 (m, 1H), 2.21–2.09 (m, 2H), 2.03–1.85 (m, 5H), 0.96–0.91 (m, 6H). **UPLC-MS** (ESI) *m/z* calcd for $[M+H]^+$ $C_{31}H_{43}N_6O_5S^+$: 611.30, found 611.24.

**P2 C1Ser (8)**

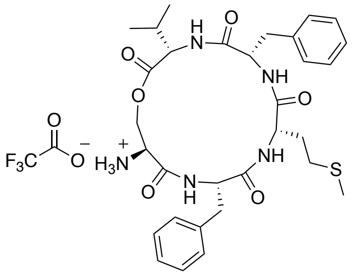

The partially protected linear peptide **S24** was synthesized in 80.0 µmol scale on Cl-Trt polystyrene resin preloaded with Fmoc-Phe-OH **S23** (0.87 mmol/g) using the general procedures for automated SPPS and manual coupling of Fmoc-Ser(TBDMS)-OH. After completed peptide elongation, a solution of $Boc_2O$ (349 mg, 1.60 mmol, 20.0 equiv), *i*-$Pr_2NEt$ (0.42 mL, 2.40 mmol, 30.0 equiv) in DMF (5.0 mL) was added the peptidyl-resin **S24** with free *N*-terminal amine and the resin was agitated at room temperature. After 1 h, the solution was removed and the Boc-protected peptidyl-resin **S25** was washed with DMF (3 × 1 min) and $CH_2Cl_2$ (3 × 1 min) and dried under high vacuum. The resin **S25** was swelled in anhydrous THF (5.0 mL) for 15 min and subsequently a solution of tetrabutylammonium fluoride (TBAF) in THF (1.0 M) (0.80 mL, 0.80 mmol, 10.0 equiv) in anhydrous THF (4.2 mL) was added to the resin and the resin was agitated at room temperature. After 1 h, the TBAF solution was removed by suction and the resin treated with a fresh TBAF–THF solution. After 1 h, the TBAF solution was removed by suction and the resin washed with DMF (3 × 1 min) and $CH_2Cl_2$ (3 × 1 min) and dried overnight under vacuum.

On-resin esterification was performed according to a previously published protocol.[2] The resin was swelled in anhydrous $CH_2Cl_2$ (3.0 mL) for 15 min and subsequently a solution of Fmoc-Val-OH (136 mg, 0.40 mmol, 5.00 equiv), *N*,*N*'-diisopropylcarbodiimide (DIC) (75.2 µL, 0.48 mmol, 6.00 equiv) and *N*-methylimidazole (NMI) (17.2 µL, 0.22 mmol, 5.40 equiv). in anhydrous $CH_2Cl_2$ (2.0 mL) was added. The resin was agitated at room temperature for 2 h and subsequently washed with anhydrous $CH_2Cl_2$ (3 × 1 min). A fresh Fmoc-Val-OH–DIC– NMI solution was added to resin and after 2 h of incubation, the resin was washed with DMF (3 × 1 min), $CH_2Cl_2$ (3 × 1 min) and DMF (3 × 1 min).

Fmoc-removal of the *O*-acylated peptidyl resin **S26** was performed by treatment of the resin with a solution of 1,8-biazabicyclo[5.4.0]undec-7-ene (DBU) in DMF (2.0 mL, 1:99, v/v) (8 × 30 s). The resin was subsequently washed with DMF (3 × 1 min), $CH_2Cl_2$ (3 × 1 min) and dried under suction for 15 min. The dried resin was treated with a solution of HFIP in $CH_2Cl_2$ (5.0 mL, 1:4, v/v) for 30 min at room temperature. The cleavage solution was removed from the resin, collected and a fresh HFIP–$CH_2Cl_2$ solution was added to the resin. After 30 min the cleavage solution was removed from the resin, collected and the resin rinsed with $CH_2Cl_2$ (5.0 mL). The combined cleavage solutions and the rinsing solution were evaporated to dryness under reduced pressure to yield the crude peptide **S27,** which was purified by preparative RP-HPLC (5–95% B over 30 min) to afford the trifluoroacetate salt of peptide **S27** as a white solid (30 mg, 35.6 μmol).

The partially protected peptide **S27** (15.0 mg, 17.8 μmol, 1.00 equiv) was dissolved in anhydrous DMF (3.0 mL) under nitrogen atmosphere and added dropwise to a solution of HATU (6.77 mg, 17.8 μmol, 1.00 equiv) and *i*-$Pr_2NEt$ (12.4 μL, 71.2 μmol, 4.00 equiv) in anhydrous DMF (15.0 mL). The reaction mixture was stirred overnight at room temperature and after full consumption of **S27** was confirmed by UPLC-MS, the reaction was reduced to dryness under reduced pressure. The remaining residue was treated with a solution of TFA in $CH_2Cl_2$ (4.0 mL, 1:1, v/v) for 1 h and subsequently concentrated under a stream of nitrogen and purified by preparative RP-HPLC (5–95% B over 30 min) to afford the trifluoroacetate salt of peptide **8** as a white solid (6.8 mg, 9.4 μmol, 53% from peptide **S27**). Purity 97% determined by UPLC ($\lambda$ = 215 nm). **$^1$H NMR** (600 MHz, DMSO-*d*$_6$) δ = 9.12 (d, *J* = 6.9 Hz, 1H), 8.86 (d, *J* = 4.5 Hz, 1H), 8.19 (s, 3H), 7.32 (d, *J* = 7.3 Hz, 1H), 7.30–7.25 (m, 7H), 7.25–7.18 (m, 2H), 7.14–7.09 (m, 2H), 4.58 (dd, *J* = 12.2, 3.1 Hz, 1H), 4.48 (dd, *J* = 9.6, 4.8 Hz, 1H), 4.37–4.29 (m, 2H), 4.24 (t, *J* = 2.5 Hz, 1H), 4.02 (ddd, *J* = 11.3, 7.1, 4.3 Hz, 1H), 3.50 (ddd, *J* = 10.4, 6.8, 3.7 Hz, 1H), 3.41–3.34 (m, 2H), 2.93 (dd, *J* = 14.1, 8.6 Hz, 1H), 2.86 (dd, *J* = 14.1, 6.8 Hz, 1H), 2.26–2.09 (m, 2H), 2.02–1.97 (m, 1H), 1.96 (s, 3H), 1.91–1.83 (m, 1H), 0.89 (d, *J* = 6.8 Hz, 3H), 0.85 (d, *J* = 6.8 Hz, 3H). **UPLC-MS** (ESI) *m/z* calcd for $[M+H]^+$ $C_{31}H_{42}N_5O_6S^+$: 612.29, found 612.21.

##### ***N,N*-Dimethyl-P2 thioether (9)**

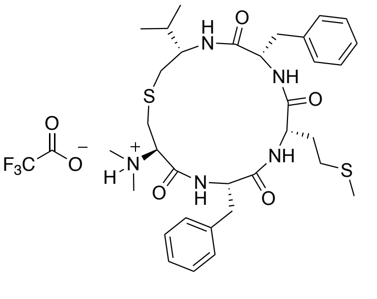

The protected linear peptide **S36** was synthesized in 25.0 μmol scale on Cl-Trt polystyrene resin preloaded with Fmoc-Phe-OH **S23** (0.87 mmol/g) using the general procedures for automated SPPS and manual coupling of *N,N*-dimethyl-Cys[Val(Fmoc)]-OH (**S35**). After completed peptide elongation, a solution of piperidine in DMF (2.0 mL, 1:4, v/v) was added to the resin and the resin agitated at room temperature for 15 min. The resin was subsequently washed with DMF (3 × 1 min), $CH_2Cl_2$ (3 × 1 min) and dried under suction for 15 min. The dried resin was treated with a cleavage cocktail (3.0 mL, TFA–*i*-$Pr_3SiH$–water, 95:2.5:2.5, v/v/v) for 30 min at room temperature. The cleavage solution was removed from the resin, collected and the resin rinsed with TFA (1.0 mL). The combined cleavage solution and the rinsing solution were concentrated under a stream of nitrogen and the cleaved peptide **S37**

was triturated in cold $Et_2O$ (10 mL) and pelleted by centrifugation. The crude peptide **S37** was obtained as TFA salt and used without further purification.

The linear peptide **S37** (25.0 μmol based on resin, 1.00 equiv) was dissolved in anhydrous DMF (3.0 mL) under nitrogen atmosphere and added dropwise to a solution of PyBOP (13.0 mg, 25.0 μmol, 1.00 equiv) and *i*-$Pr_2NEt$ (17.4 μL, 100 μmol, 4.00 equiv) in anhydrous DMF (22.0 mL). The reaction mixture was stirred overnight at room temperature and after full consumption of **S37** was confirmed by UPLC-MS, the reaction was reduced to dryness under reduced pressure. The remaining residue was purified by preparative RP-HPLC (20–70% B over 30 min) and two diastereomers of **9** were isolated due to partial loss of optical purity of building block **S35**. The major isomer was concluded to correspond to desired peptide and was obtained as the trifluoroacetate salt of peptide **9** as a white solid (4.3 mg, 5.7 μmol, 23% based on resin). Purity 95% determined by UPLC ($\lambda$ = 215 nm). **$^1$H NMR** (600 MHz, DMSO-$d_6$) δ = 10.01 (s, 1H), 9.00 (d, $J$ = 6.8 Hz, 1H), 8.87 (d, $J$ = 9.1 Hz, 1H), 8.17 (d, $J$ = 8.1 Hz, 1H), 7.31–7.23 (m, 7H), 7.23–7.16 (m, 4H), 4.86–4.79 (m, 1H), 4.06 (ddd, $J$ = 12.1, 8.8, 4.2 Hz, 1H), 3.87–3.83 (m, 1H), 3.77–3.69 (m, 1H), 3.54 (ddd, $J$ = 9.0, 7.1, 5.5 Hz, 1H), 3.25–3.08 (m, 5H), 2.75 (dd, $J$ = 13.7, 11.1 Hz, 1H), 2.62 (dd, $J$ = 11.0, 3.8 Hz, 1H), 2.53–2.47 (m, 2H), 2.28–2.23 (m, 2H), 2.08–1.93 (m, 5H), 1.83–1.74 (m, $J$ = 6.7 Hz, 1H), 0.95–0.90 (m, 6H). **UPLC-MS** (ESI) *m/z* calcd for $[M+H]^+$ $C_{33}H_{48}N_5O_4S_2^+$: 642.31, found 642.36.

**P2 thioether (10)**

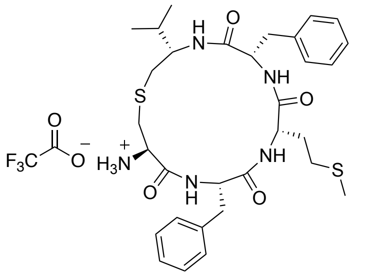

The fully protected linear peptide **S38** was synthesized in 80.0 μmol scale on Cl-Trt polystyrene resin preloaded with Fmoc-Phe-OH **S23** (0.87 mmol/g) using the general procedures for automated SPPS and manual coupling of Boc-Cys(S*t*-Bu)-OH. After completed peptide elongation, a solution of *N*-methylmorpholine (NMM) (44.2 μL, 0.40 mmol, final concentration = 0.1 M) in a mixture of β-mercaptoethanol–DMF (4.0 mL, 1:4, v/v) was added to the peptidyl resin **S38** and the resin was agitated overnight at room temperature. The next day, the thiol-containing solution was removed by suction and the resin **S39** was washed with DMF (3 × 1 min), $CH_2Cl_2$ (3 × 1 min) and dried dried overnight under vacuum.

The resin **S39** was swelled in anhydrous DMF (3.0 mL) for 15 min and in a separate flask, a suspension of $CsCO_3$ (130 mg, 0.40 mmol, 5.00 equiv) in anhydrous DMF (5.0 mL) was sonicated for 15 min and subsequently Fmoc-Val-iodide (**S30**) (174 mg, 0.40 mmol, 5.00 equiv) was added. The alkylation reaction mixture was added to the resin **S39** and the resin was agitated at room temperature overnight. The next day, the solution was removed by suction and the alkylated peptidyl resin **S40** was washed with DMF (3 × 1 min), MeOH (3 × 1 min) and DMF (3 × 1 min) and a solution of piperidine in DMF (3.0 mL, 1:4, v/v) was added to the resin and the resin agitated at room temperature for 15 min. The resin was subsequently washed with DMF (3 × 1 min), $CH_2Cl_2$ (3 × 1 min) and dried under suction for 15 min. The dried resin was treated with a solution of HFIP in $CH_2Cl_2$ (5.0 mL, 1:4, v/v) for 30 min at room temperature. The cleavage solution was removed from the resin, collected and a fresh HFIP–$CH_2Cl_2$ solution was added to the resin. After 30 min the cleavage solution was removed from the resin, collected and the resin rinsed with $CH_2Cl_2$ (5.0 mL). The combined cleavage solutions and the rinsing solution were evaporated to dryness under reduced pressure to yield the crude peptide **S41**, which was purified by preparative RP-HPLC (5–95% B over 30 min) to afford the trifluoroacetate salt of peptide **S41** as a white solid (11.2 mg, 13.2 μmol).

The partially-protected peptide **S41** (11.2 mg, 13.2 μmol, 1.00 equiv) was dissolved in anhydrous DMF (2.0 mL) under nitrogen atmosphere and added dropwise to a solution of PyBOP (6.87 mg, 13.2 μmol, 1.00 equiv) and *i*-$Pr_2NEt$ (9.20 μL, 52.8 μmol, 4.00 equiv) in anhydrous DMF (11.0 mL). The reaction mixture was stirred overnight at room temperature and after full consumption of **S41** was confirmed by UPLC-MS, the reaction was reduced to dryness under reduced pressure. The remaining residue was treated with a solution of TFA in $CH_2Cl_2$ (3.0 mL, 1:1, v/v) for 1 h and subsequently concentrated under a stream of nitrogen and purified by preparative RP-HPLC (5–95% B over 30 min) to afford the trifluoroacetate salt of peptide **10** as a white solid (4.0 mg, 5.5 μmol, 42% from peptide **S41**). Purity 97% determined by UPLC ($\lambda$ = 215 nm). **$^1$H NMR** (600 MHz, DMSO-$d_6$) δ = 8.95 (d, $J$ = 7.1 Hz, 1H), 8.86 (d, $J$ = 5.6 Hz, 1H), 8.15 (s, 3H), 7.85 (d, $J$ = 7.5 Hz, 1H), 7.32–7.20 (m, 8H), 7.20–7.16 (m, 3H), 4.54–4.41 (m, 1H), 4.08–4.00 (m, 1H), 3.98–3.92 (m, 1H), 3.58–3.53 (m, 1H), 3.50–3.44 (m, 1H), 3.33–3.23 (m, 2H), 3.07 (dd, $J$ = 14.0, 3.8 Hz, 1H), 2.99–2.86 (m, 3H), 2.83 (dd, $J$ = 14.0, 7.6 Hz, 1H), 2.78 (dd, $J$ = 12.4, 3.6 Hz, 1H), 2.05–1.96 (m, 2H), 1.97–1.90 (m, 4H), 1.90–1.81 (m, 2H), 0.89 (d, $J$ = 6.7 Hz, 3H), 0.87 (d, $J$ = 6.8 Hz, 3H). **UPLC-MS** (ESI) *m/z* calcd for $[M+H]^+$ $C_{31}H_{44}N_5O_4S_2^+$: 614.28, found 614.25.

### Copies of UPLC traces of synthesized peptides

*N,N*-dimethyl P2 (**1**)

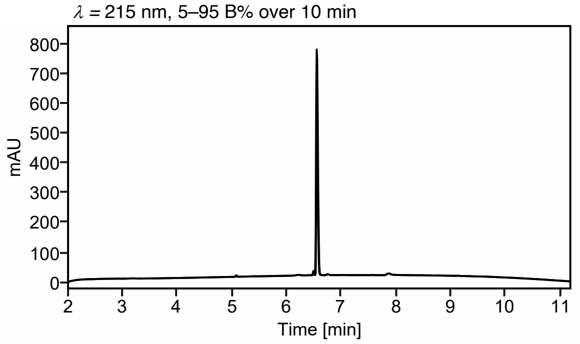

*N,N*-dimethyl P2 F2A (**2**)

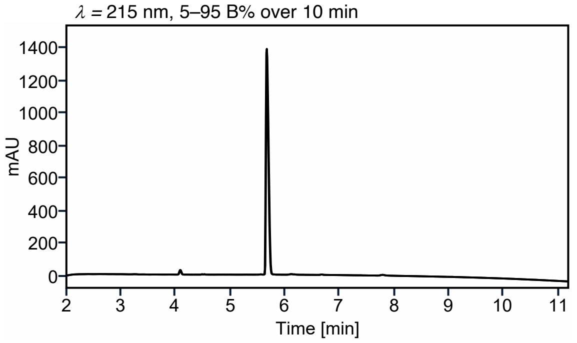

*N,N*-dimethyl P2 M3A (**3**)

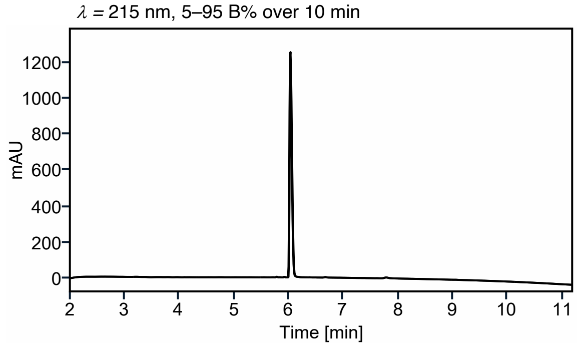

*N,N*-dimethyl P2 F4A (**4**)

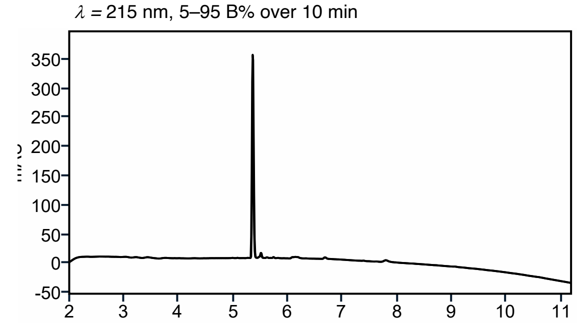

*N,N*-dimethyl P2 V5A (**3**)

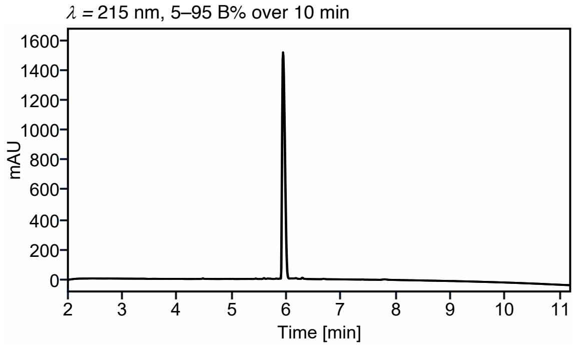

*N,N*-dimethyl P2 C1Dap (**6**)

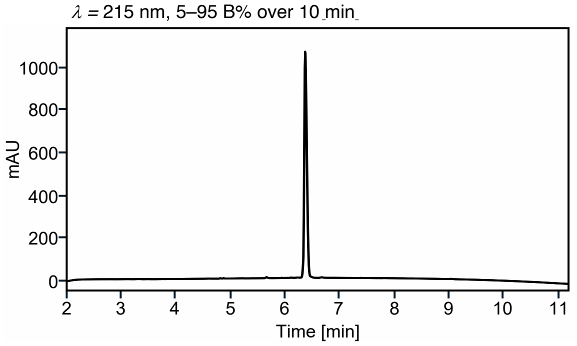

**Figure S13.** Copies of UPLC traces of synthesized peptides.

P2 C1Dap (**7**)

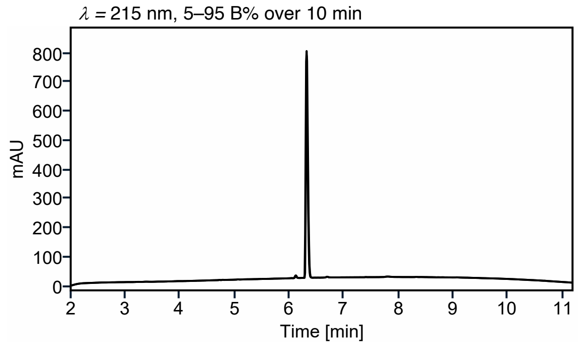

P2 C1Ser (**8**)

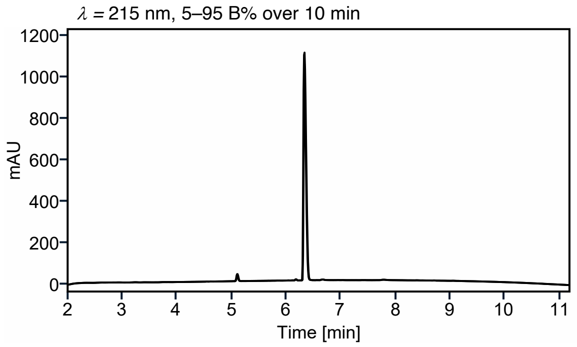

*N,N*-dimethyl P2 thioether (**9**)

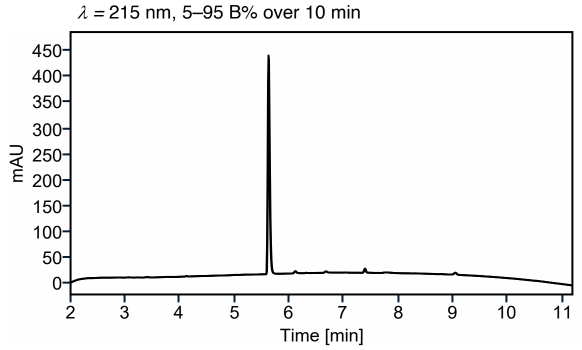

P2 thioether (**10**)

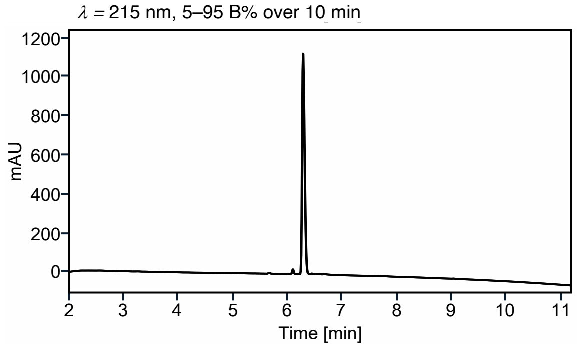

**Figure S14.** Copies of UPLC traces of synthesized peptides.

### Copies of NMR spectra of synthesized compounds

##### *N*,*N*-Dimethyl-*S*-trityl-*L*-cysteine (S2)

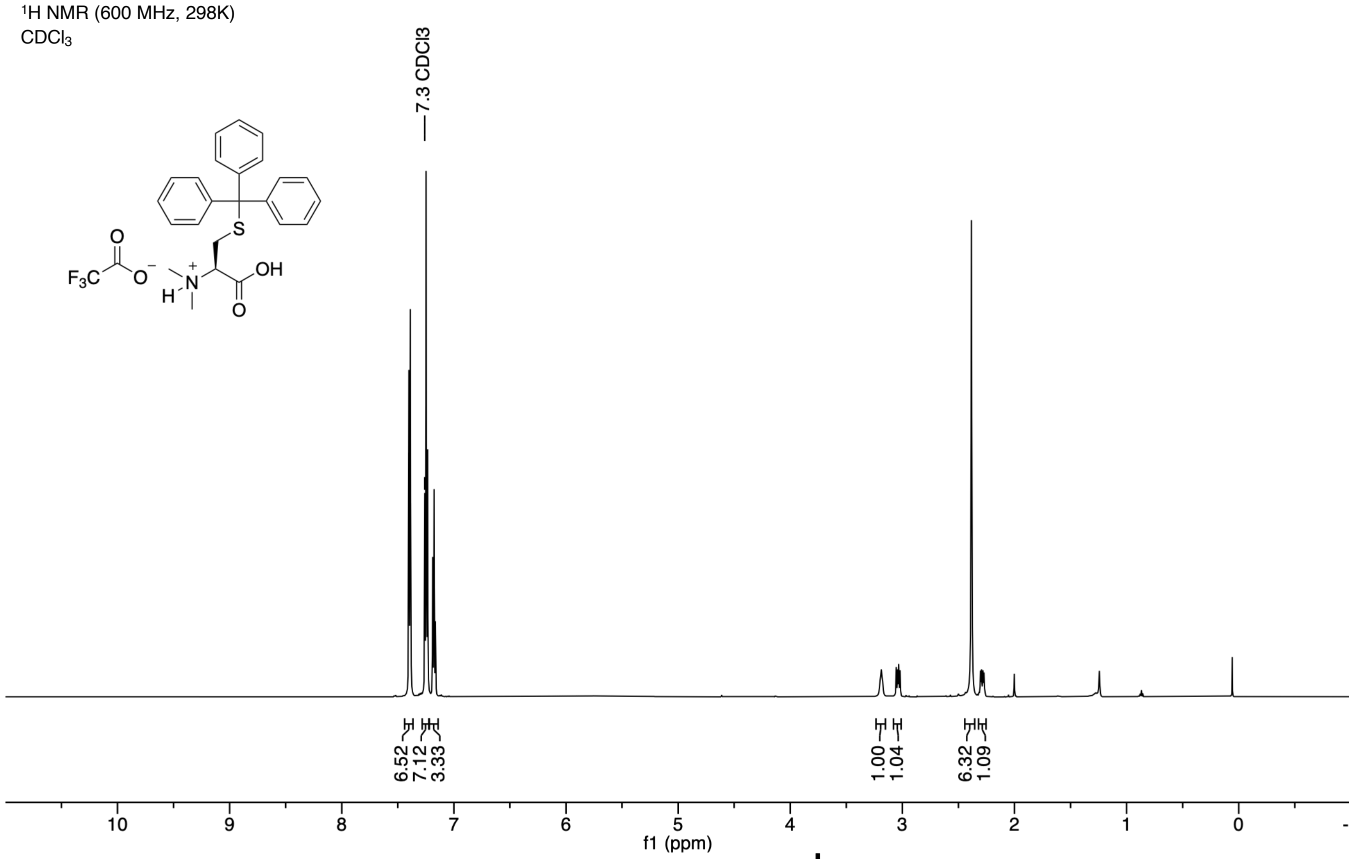

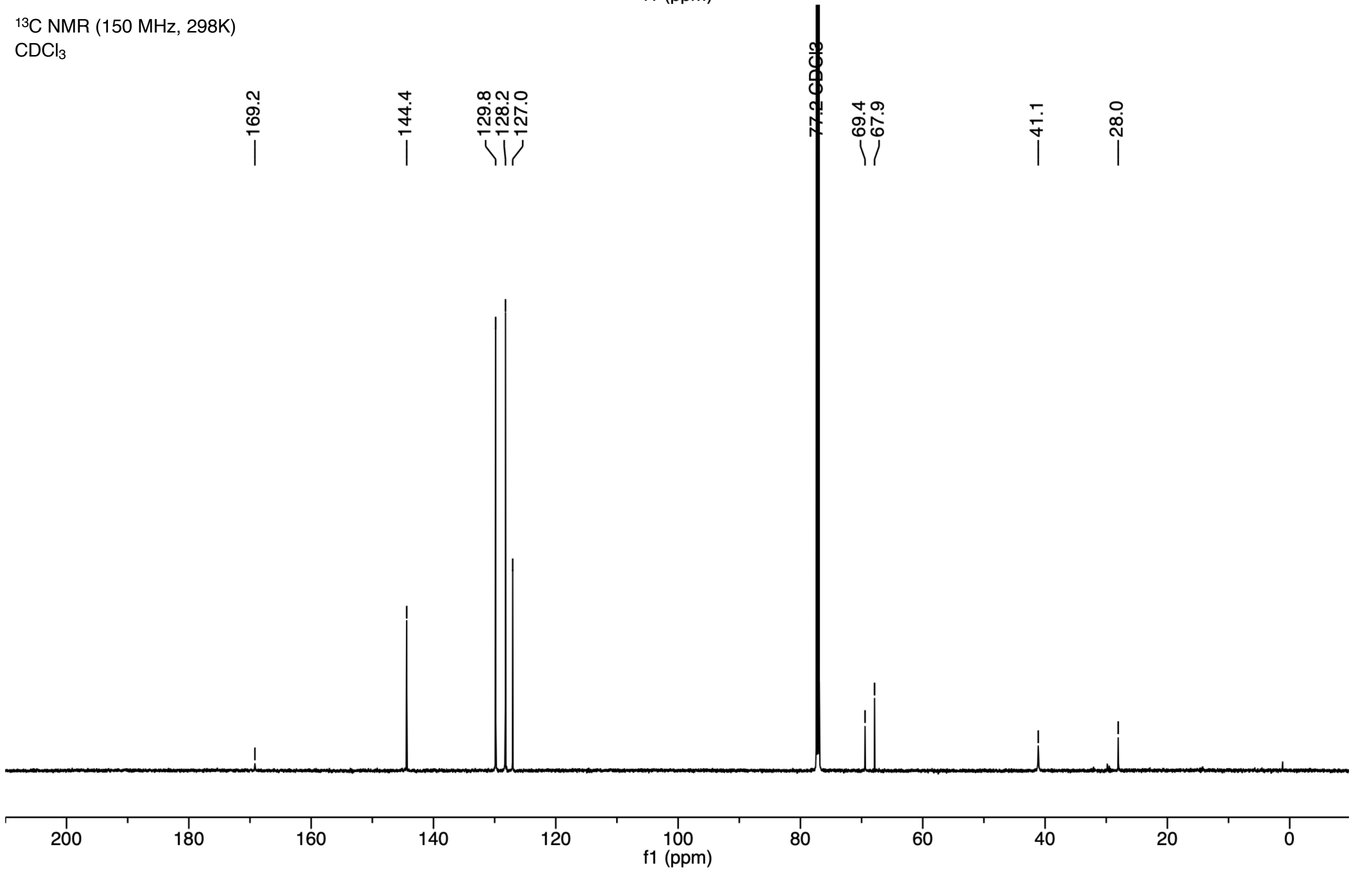

**(*S*)-3-((((9*H*-Fluoren-9-yl)methoxy)carbonyl)amino)-2-(dimethylamino)propanoic acid (S16)**

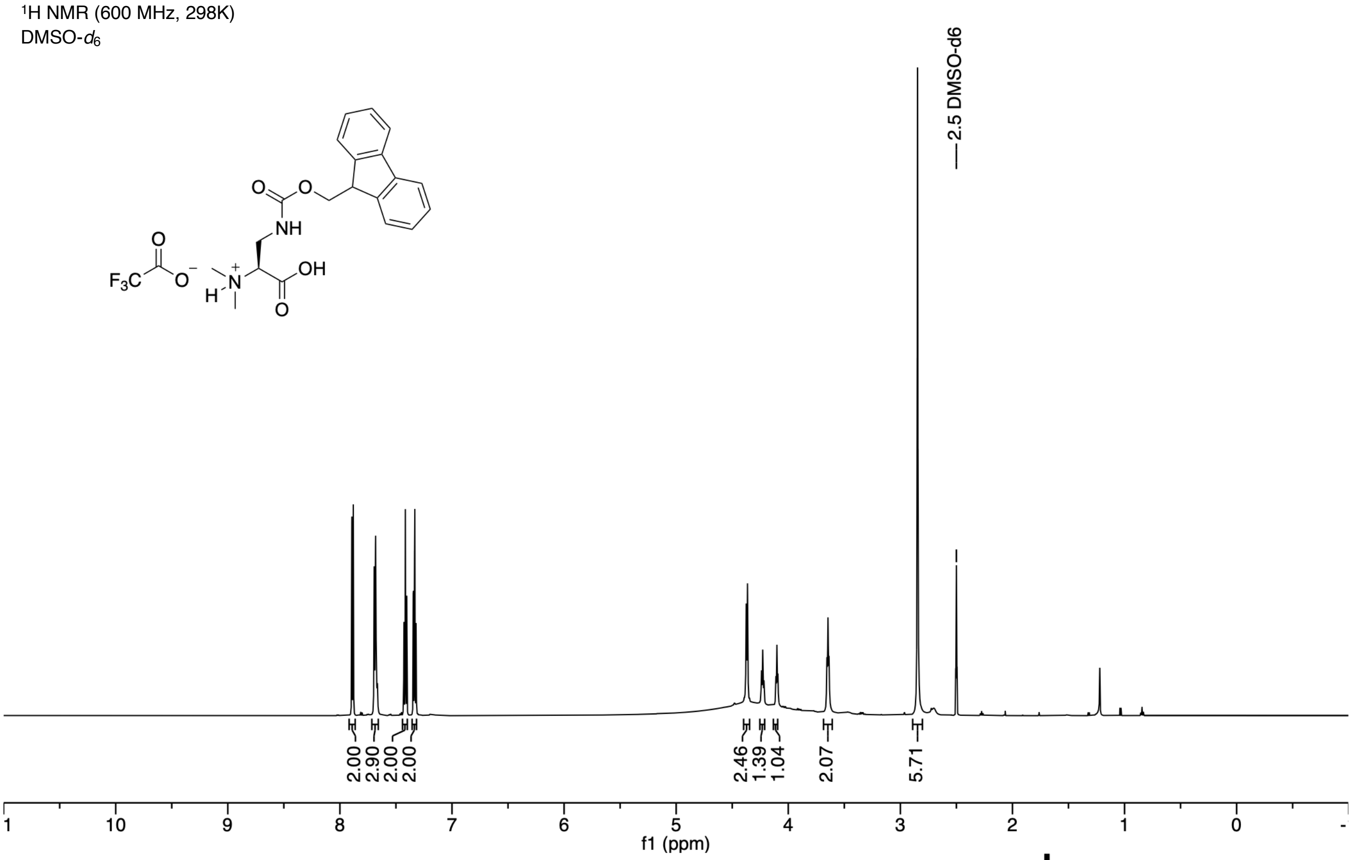

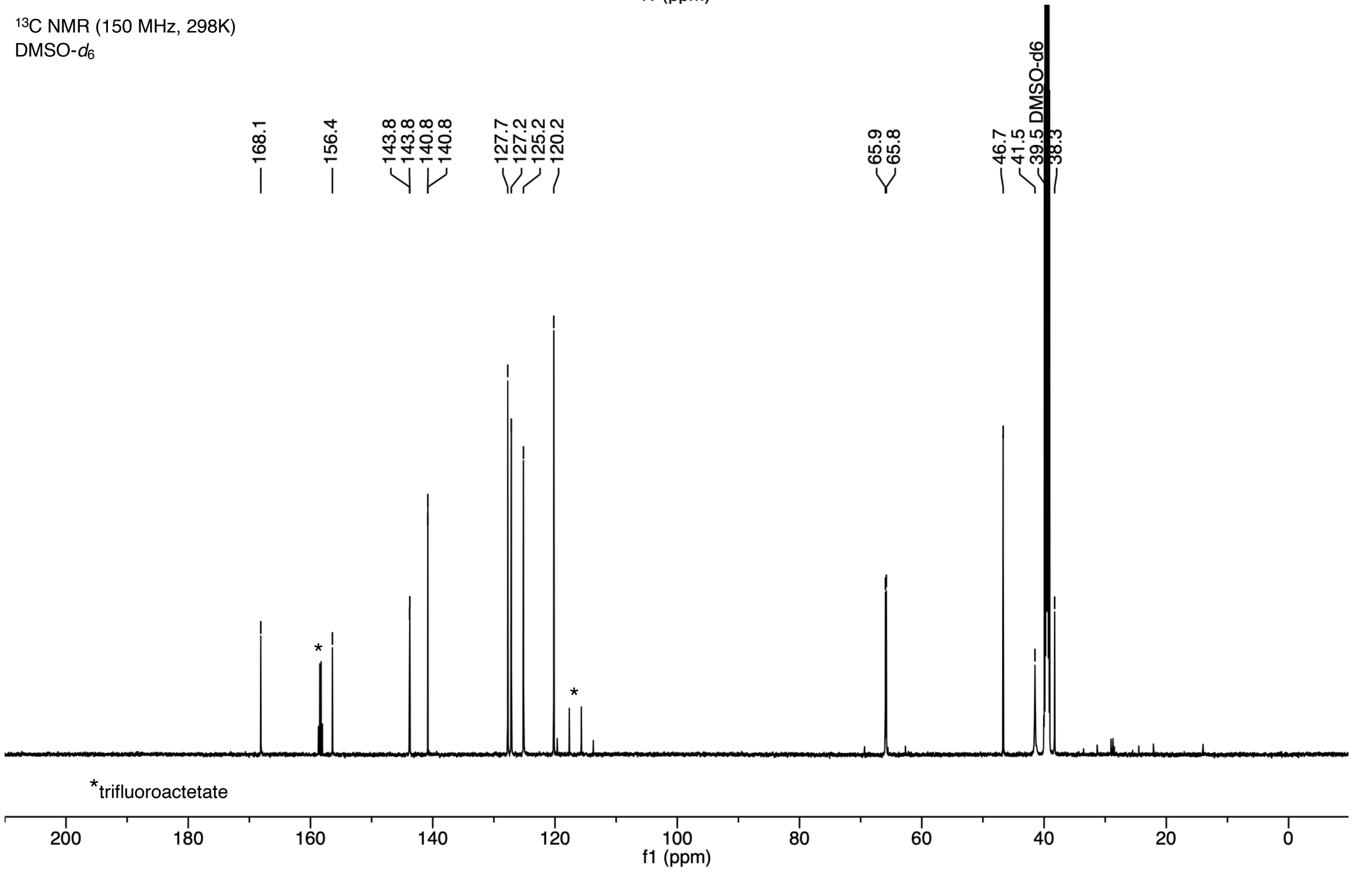

#### (9*H*-Fluoren-9-yl)methyl (*S*)-(1-iodo-3-methylbutan-2-yl)carbamate (S30)

$^{1}$H NMR (600 MHz, 298K)
DMSO-$d_6$

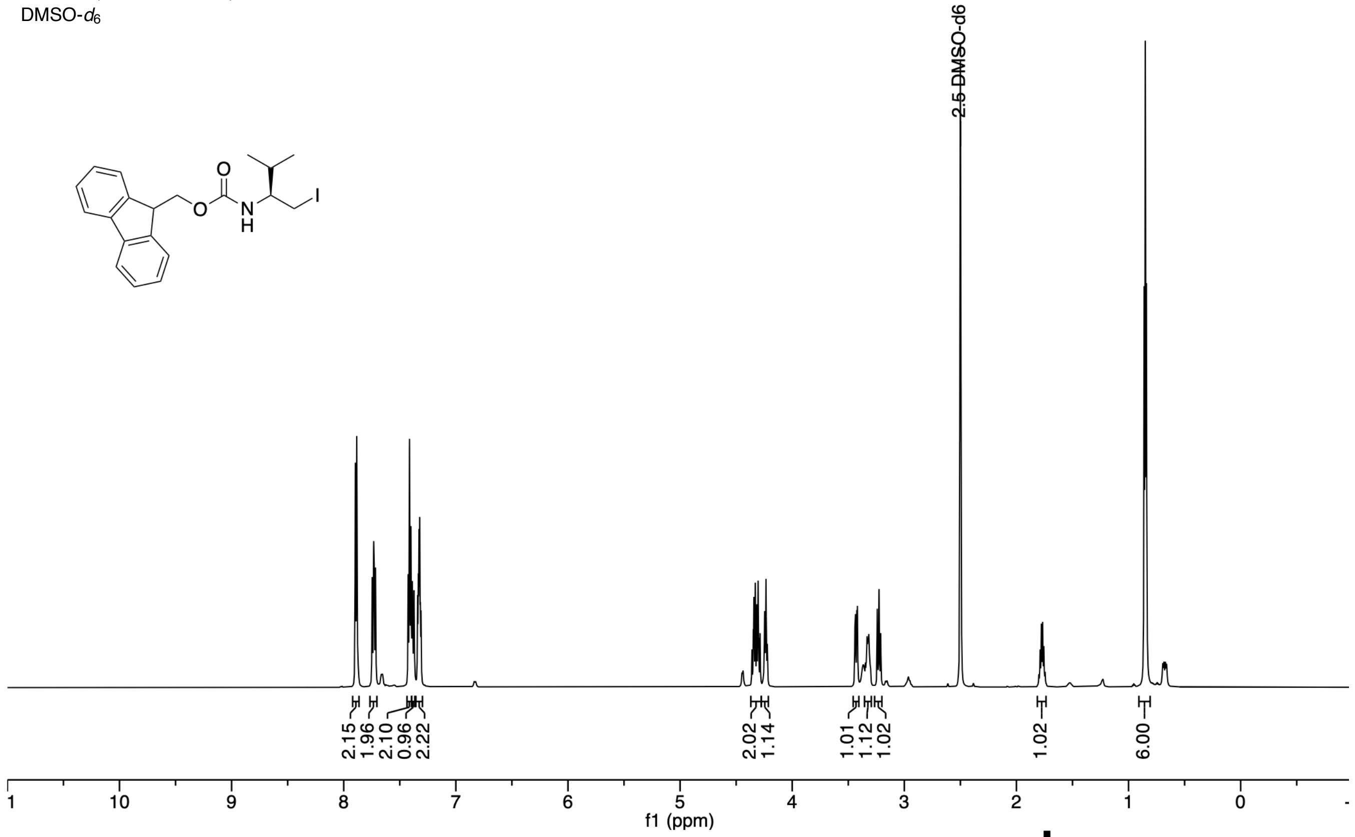

$^{13}$C NMR (150 MHz, 298K)
DMSO-$d_6$

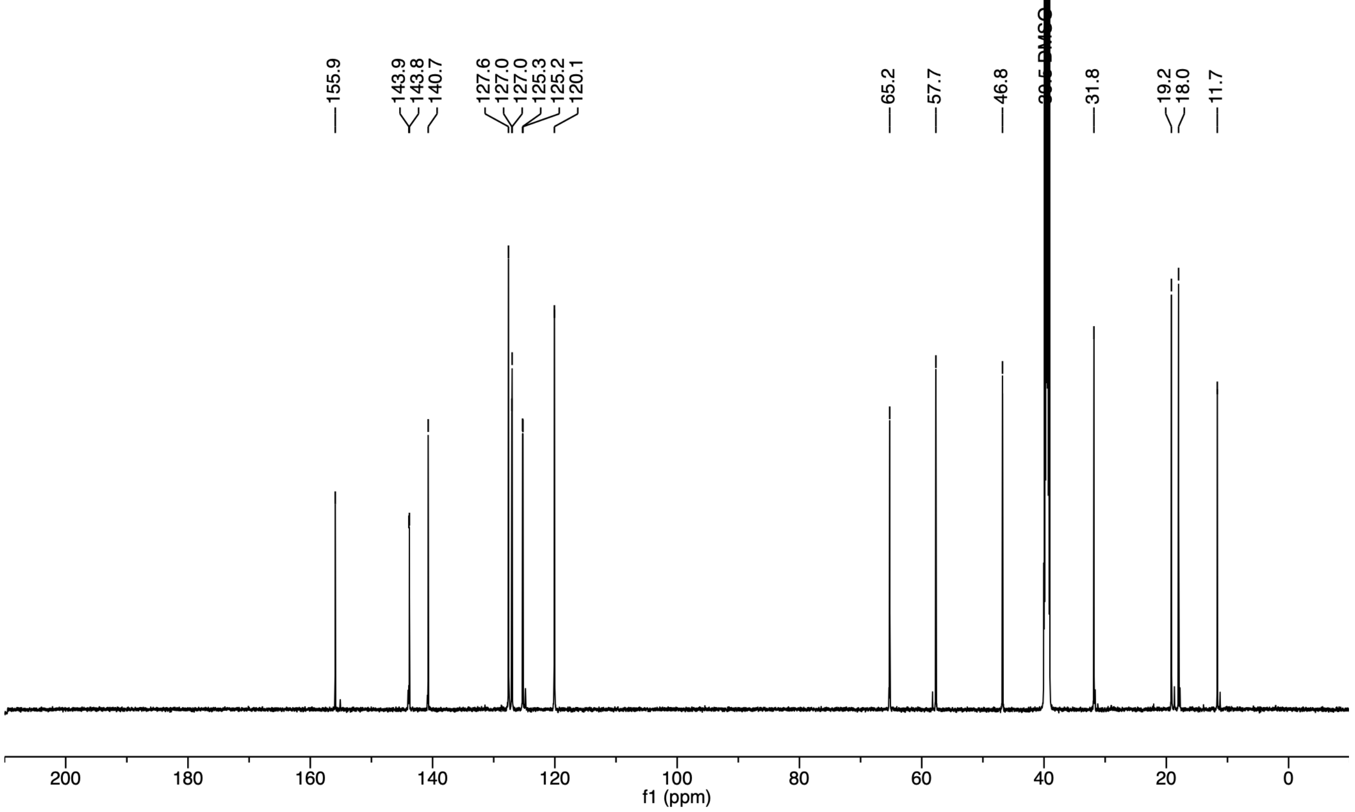

**_S_-((_S_)-2-((((9_H_-Fluoren-9-yl)methoxy)carbonyl)amino)-3-methylbutyl)-_N_,_N_-dimethyl-_L_-cysteine (S35)**

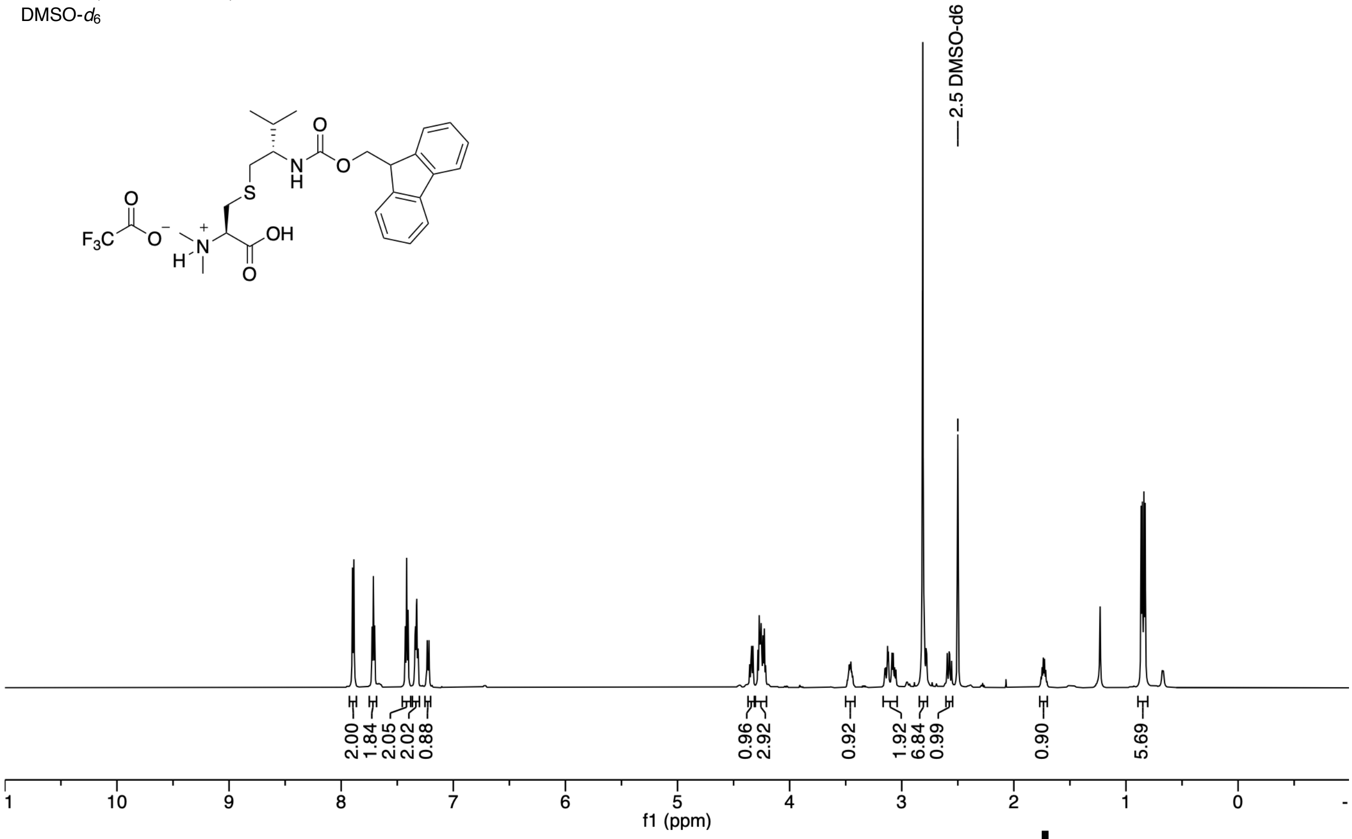

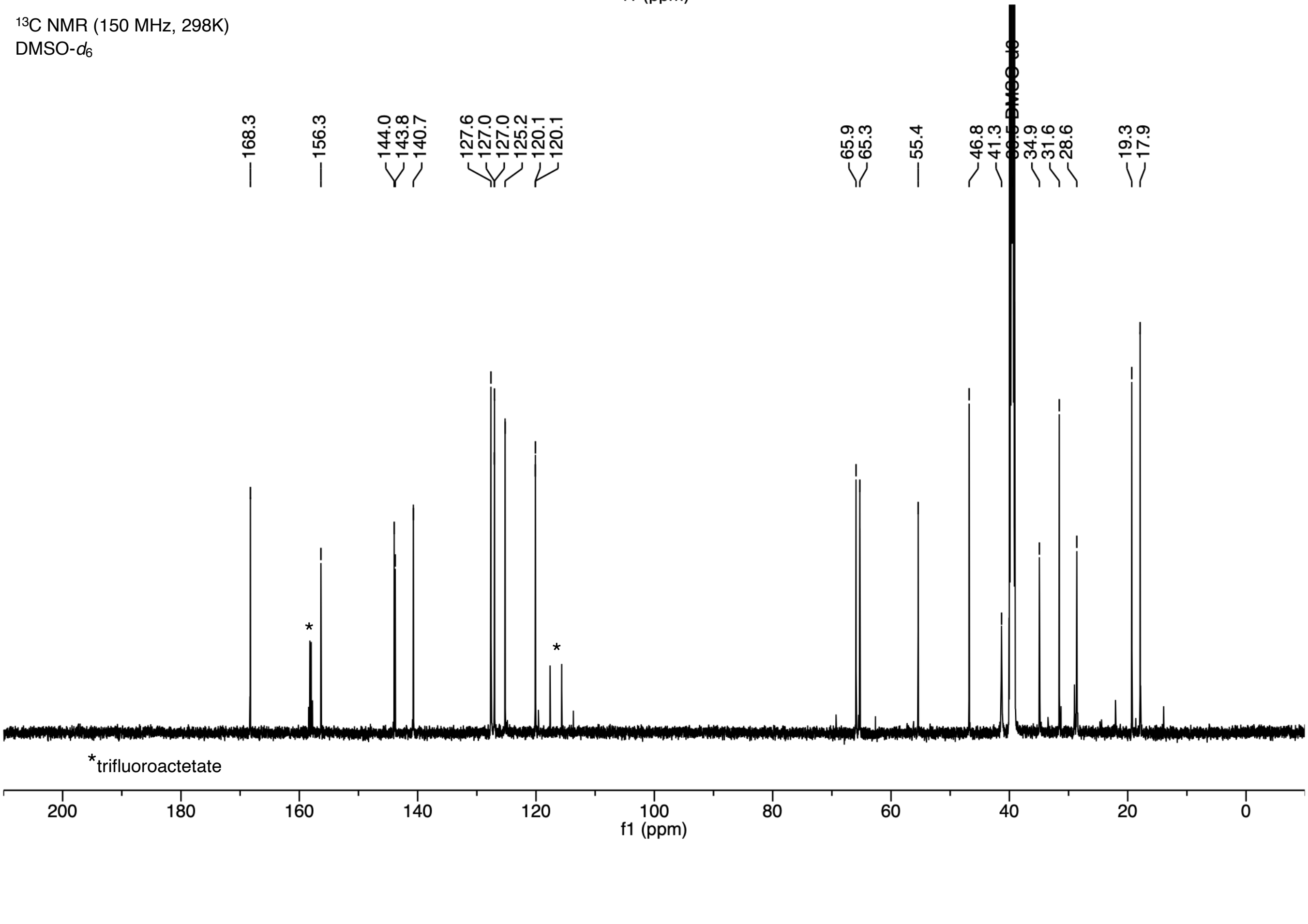

#### *N,N*-Dimethyl-P2 (1)

$^{1}$H NMR (600 MHz, 298K)
DMSO-$d_6$

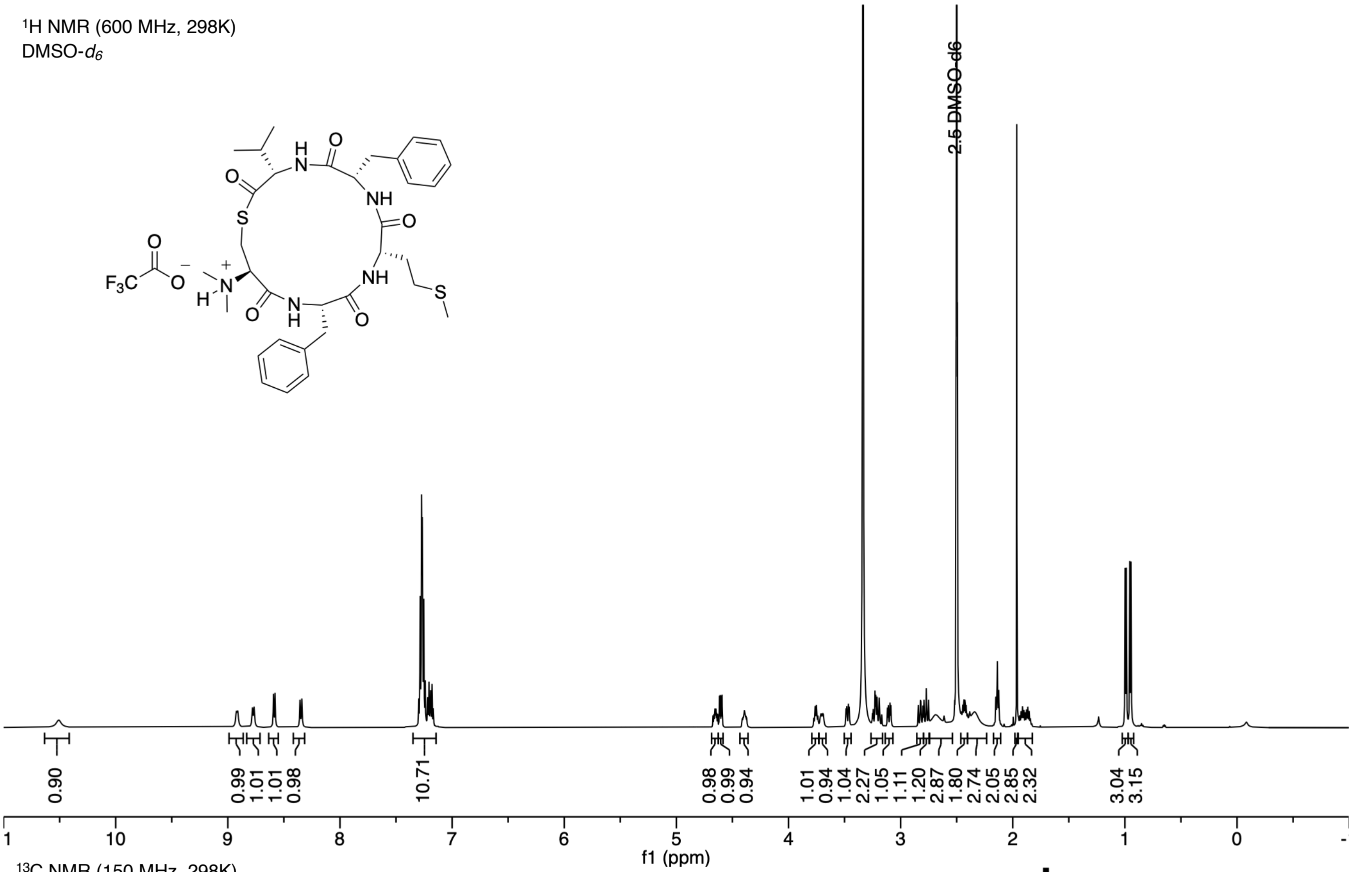

$^{13}$C NMR (150 MHz, 298K)
DMSO-$d_6$

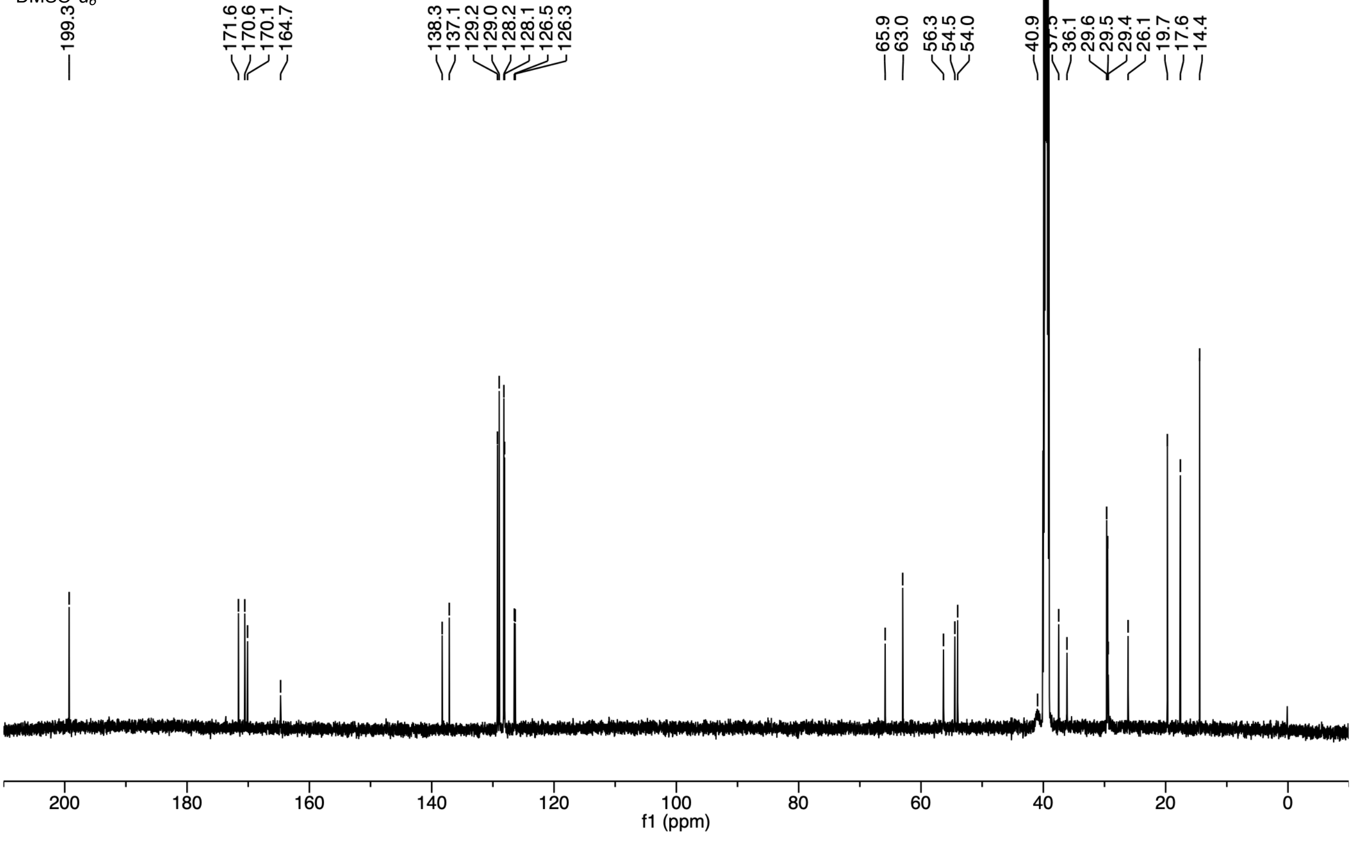

#### *N,N*-Dimethyl-P2 F2A (2)

$^{1}$H NMR (600 MHz, 298K)
DMSO-*d*$_6$

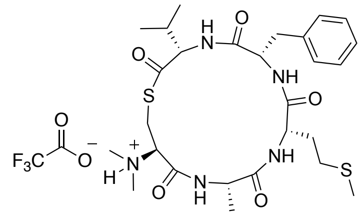

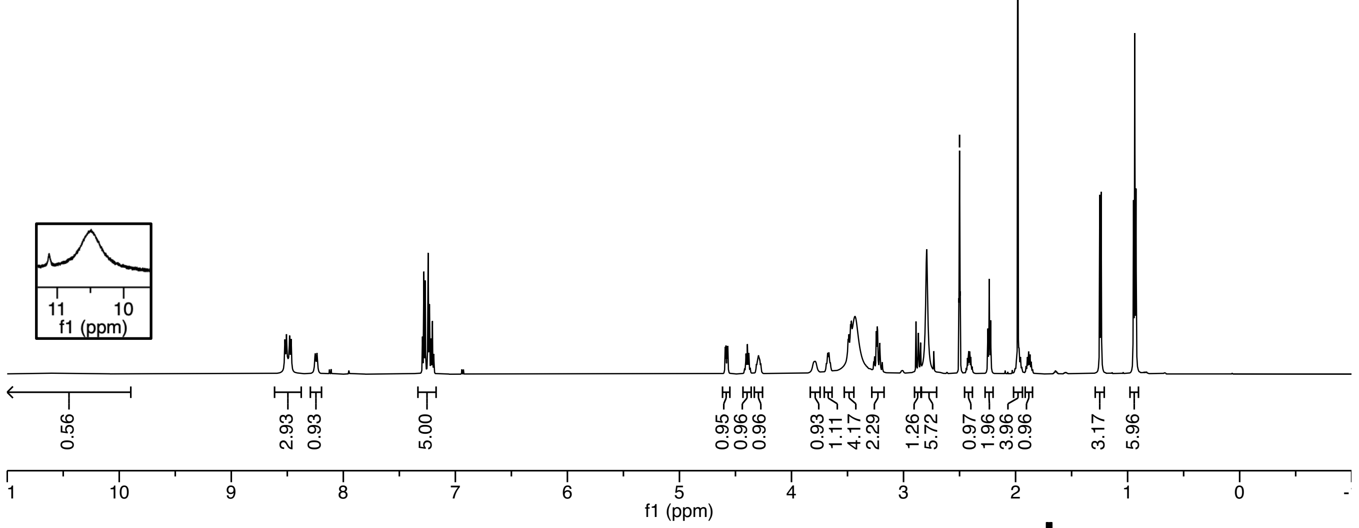

$^{13}$C NMR (150 MHz, 298K)
DMSO-*d*$_6$

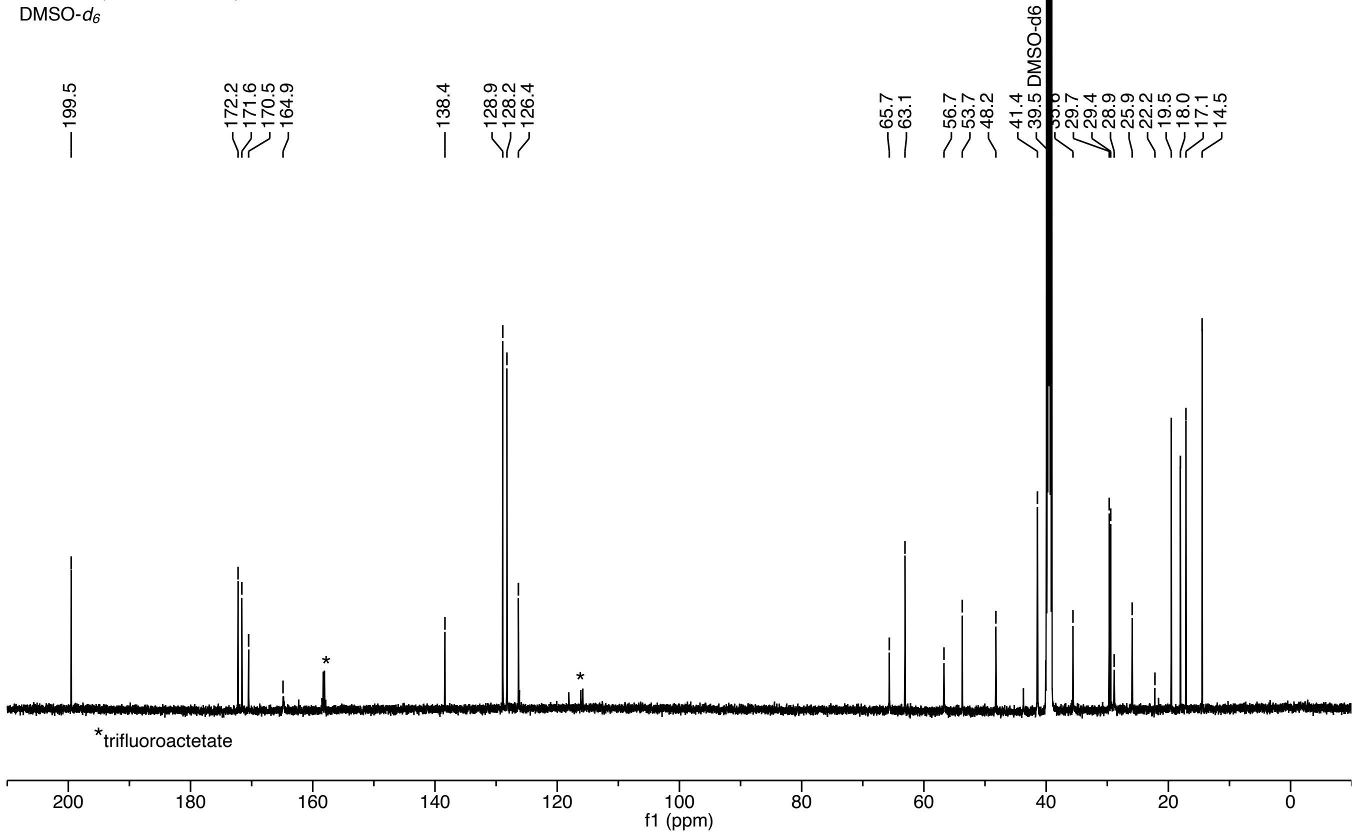

#### *N*,*N*-Dimethyl-P2 M3A (3)

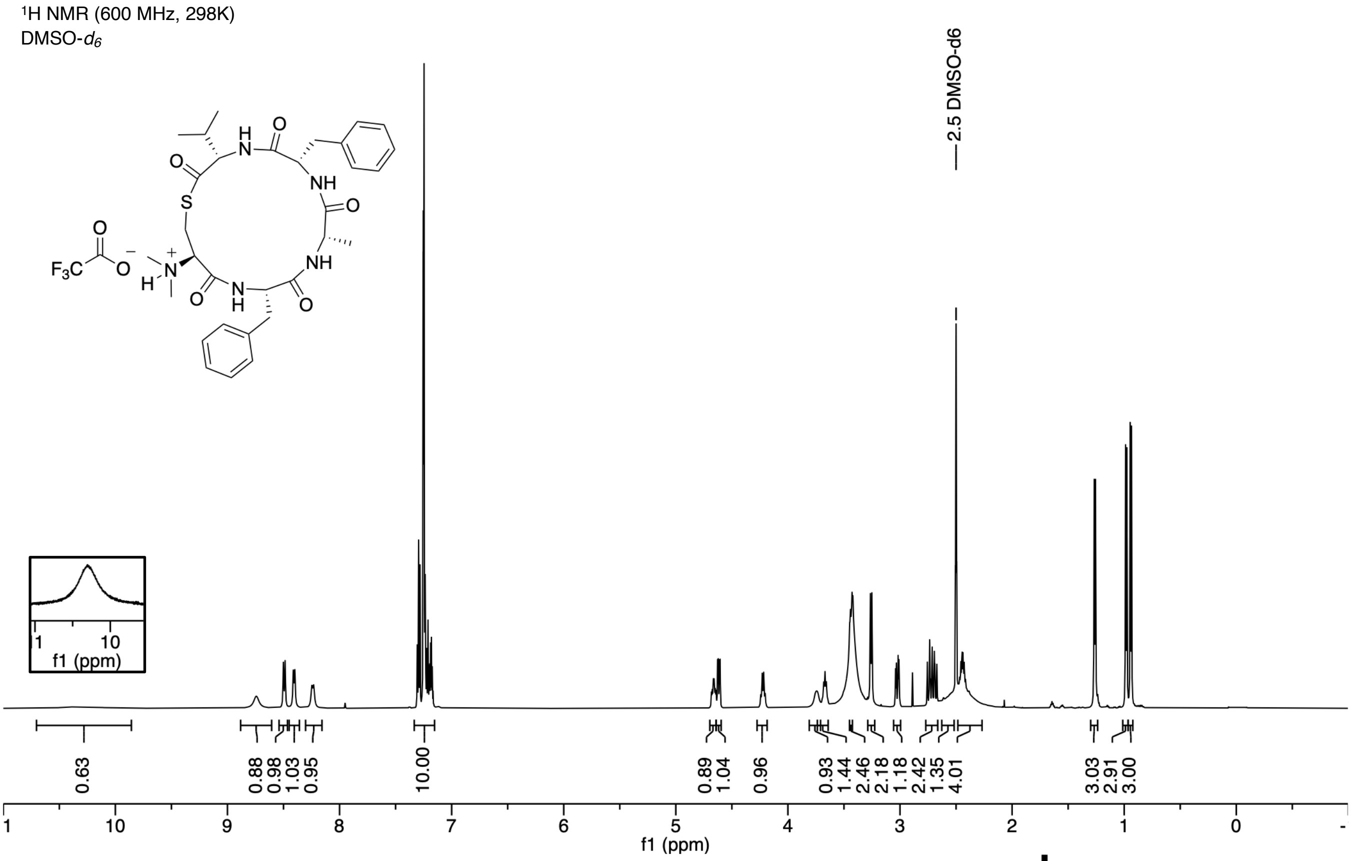

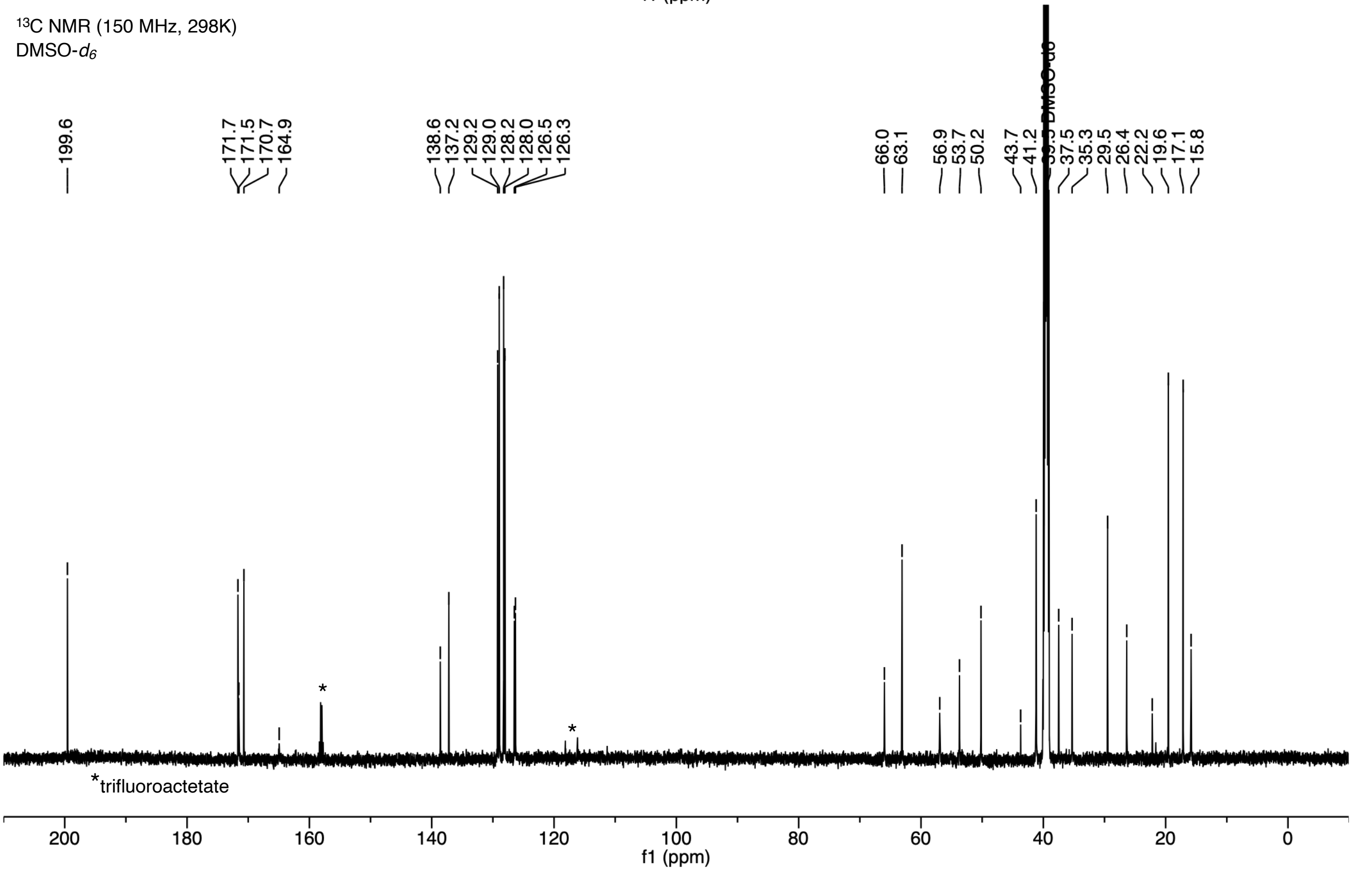

#### *N*,*N*-Dimethyl-P2 F4A (4)

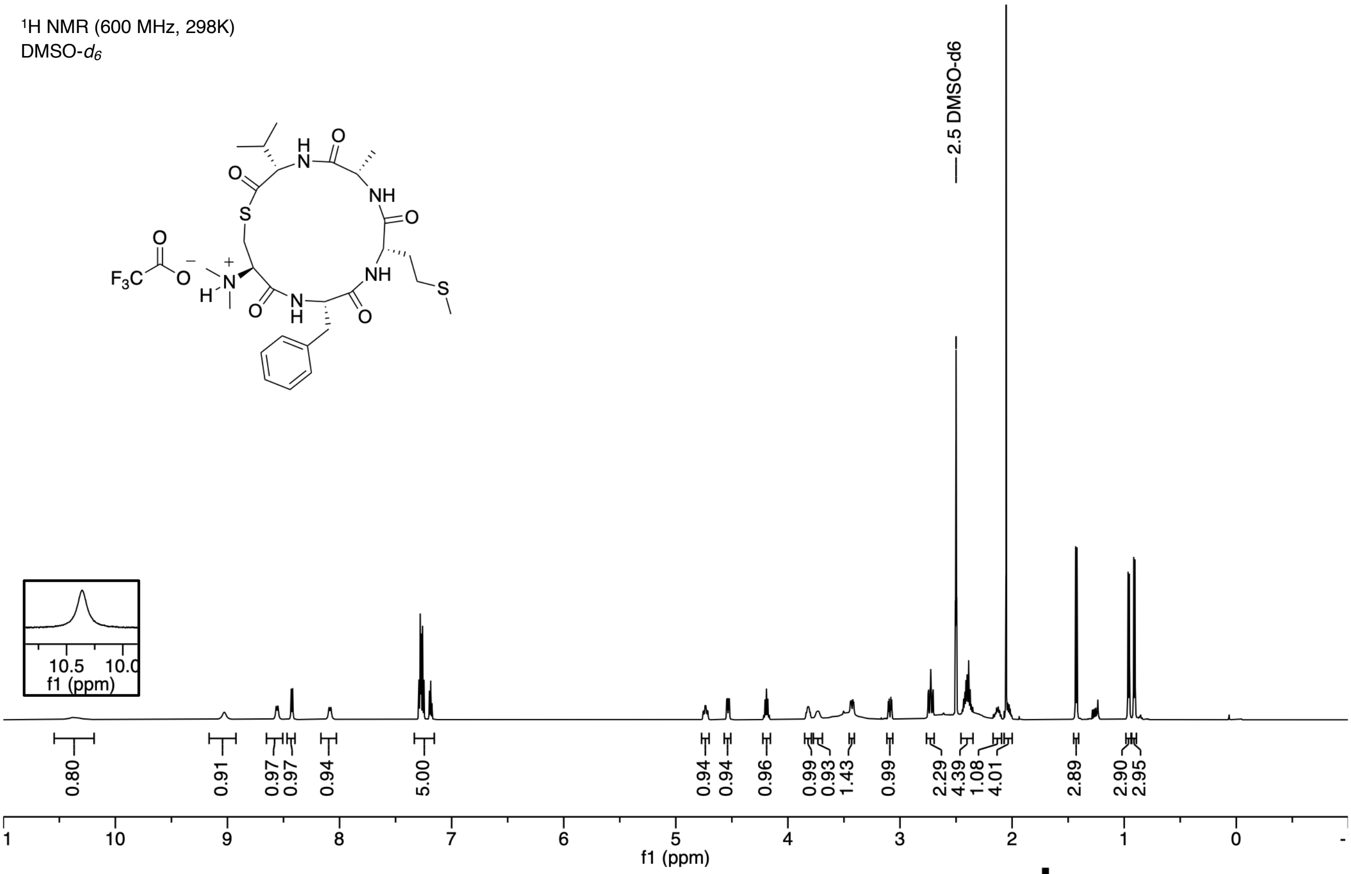

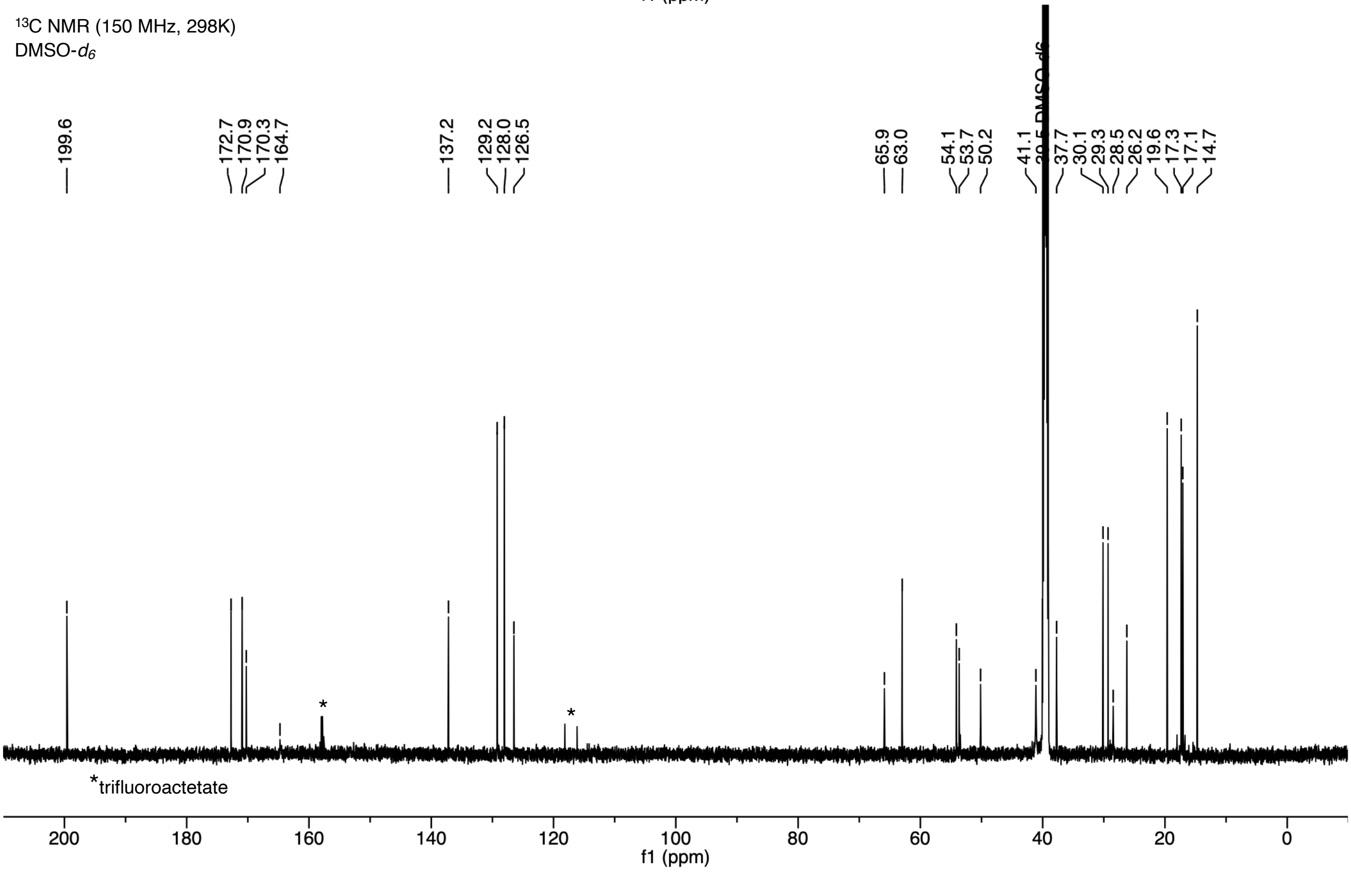

##### *N,N*-Dimethyl-P2 V5A (5)

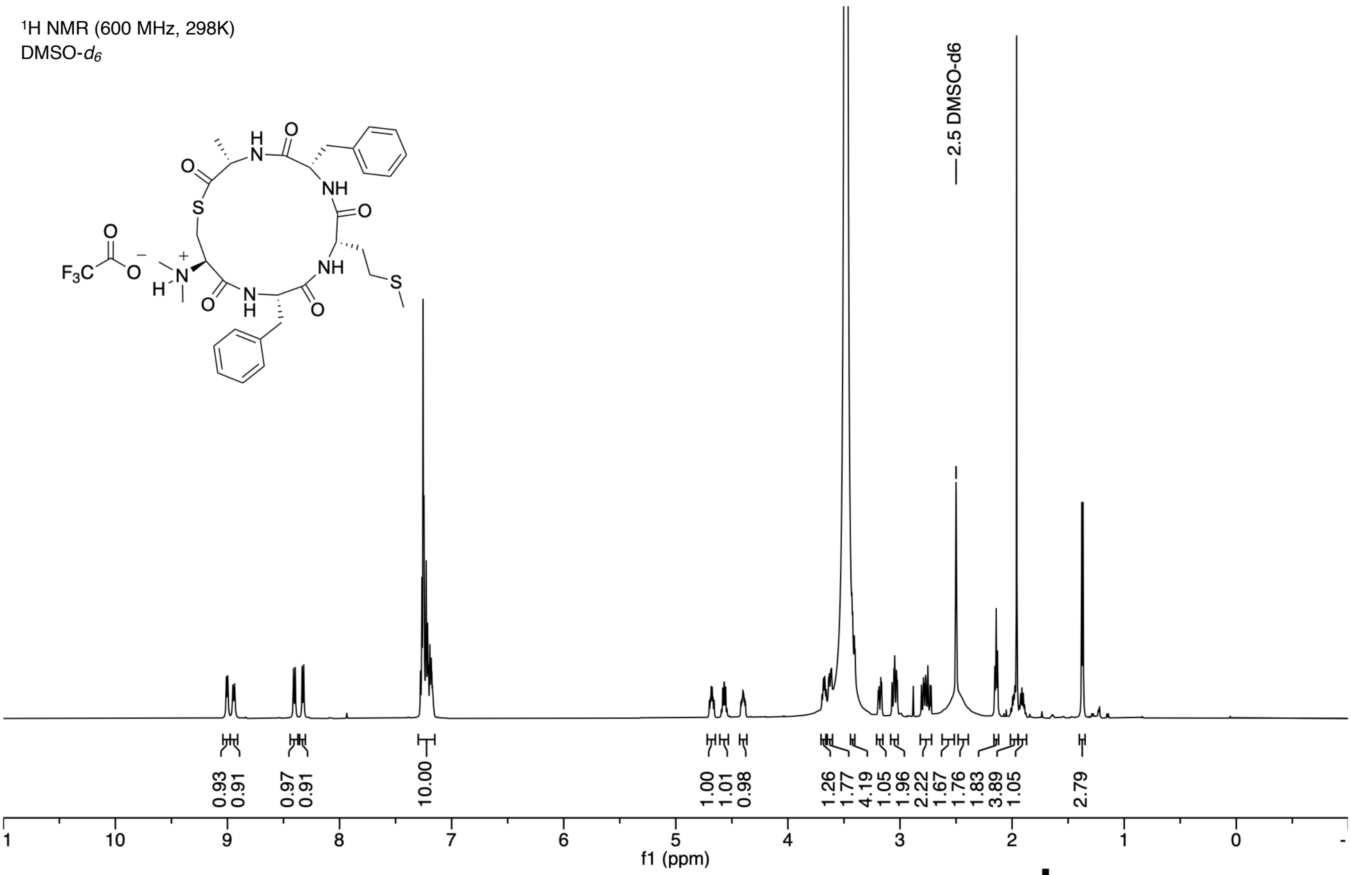

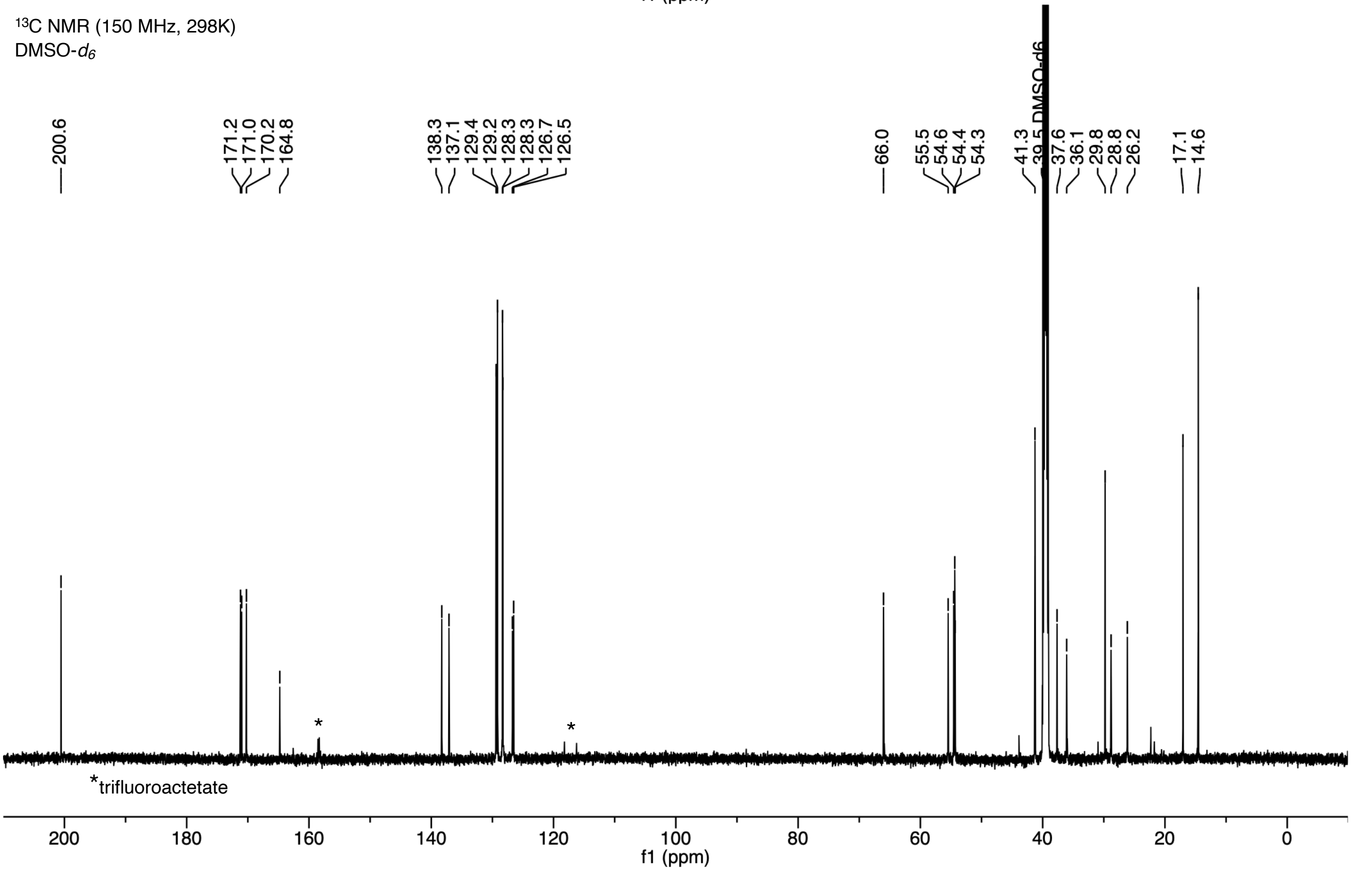

#### *N*,*N*-Dimethyl-P2 C1Dap (6)

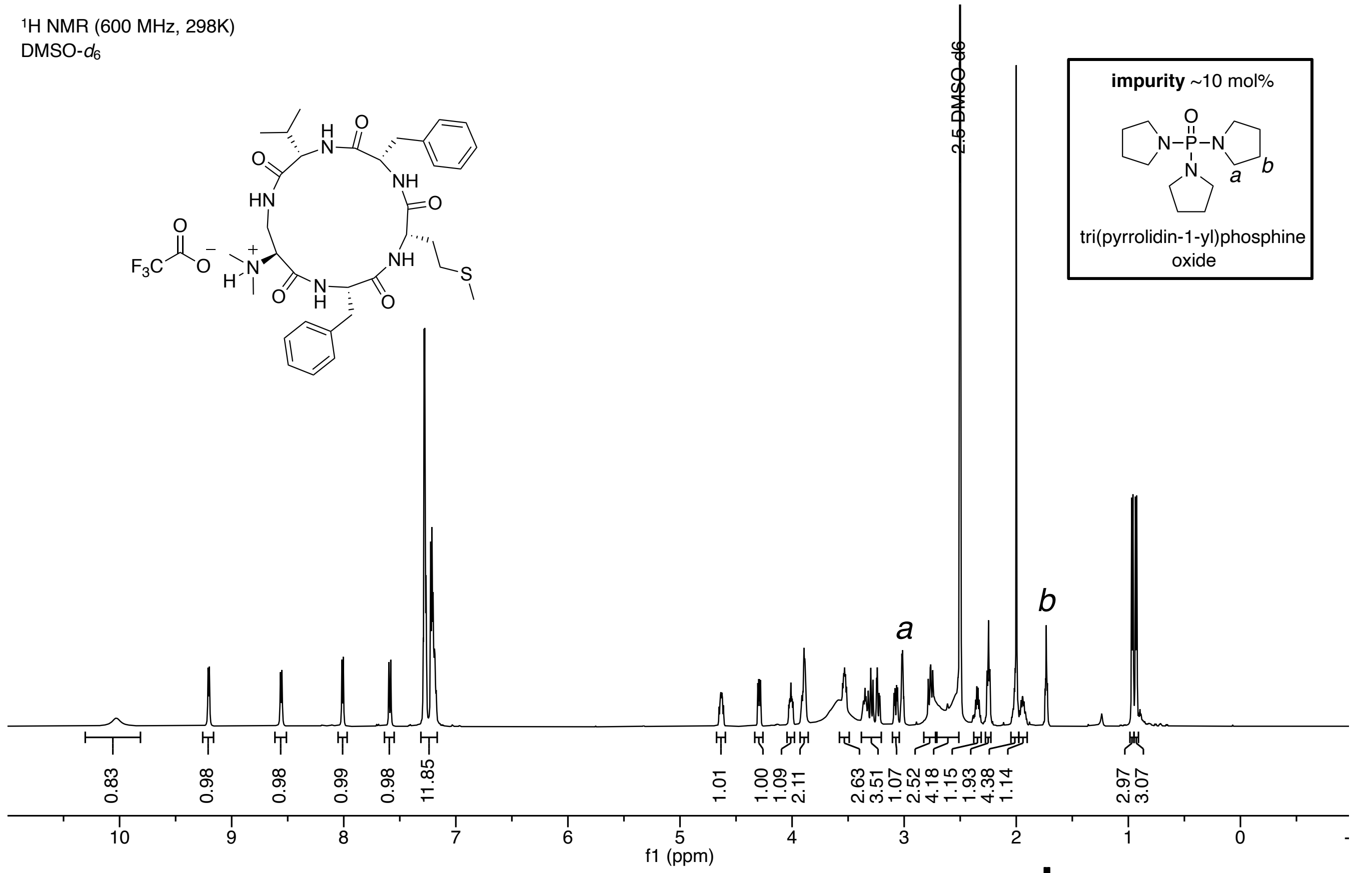

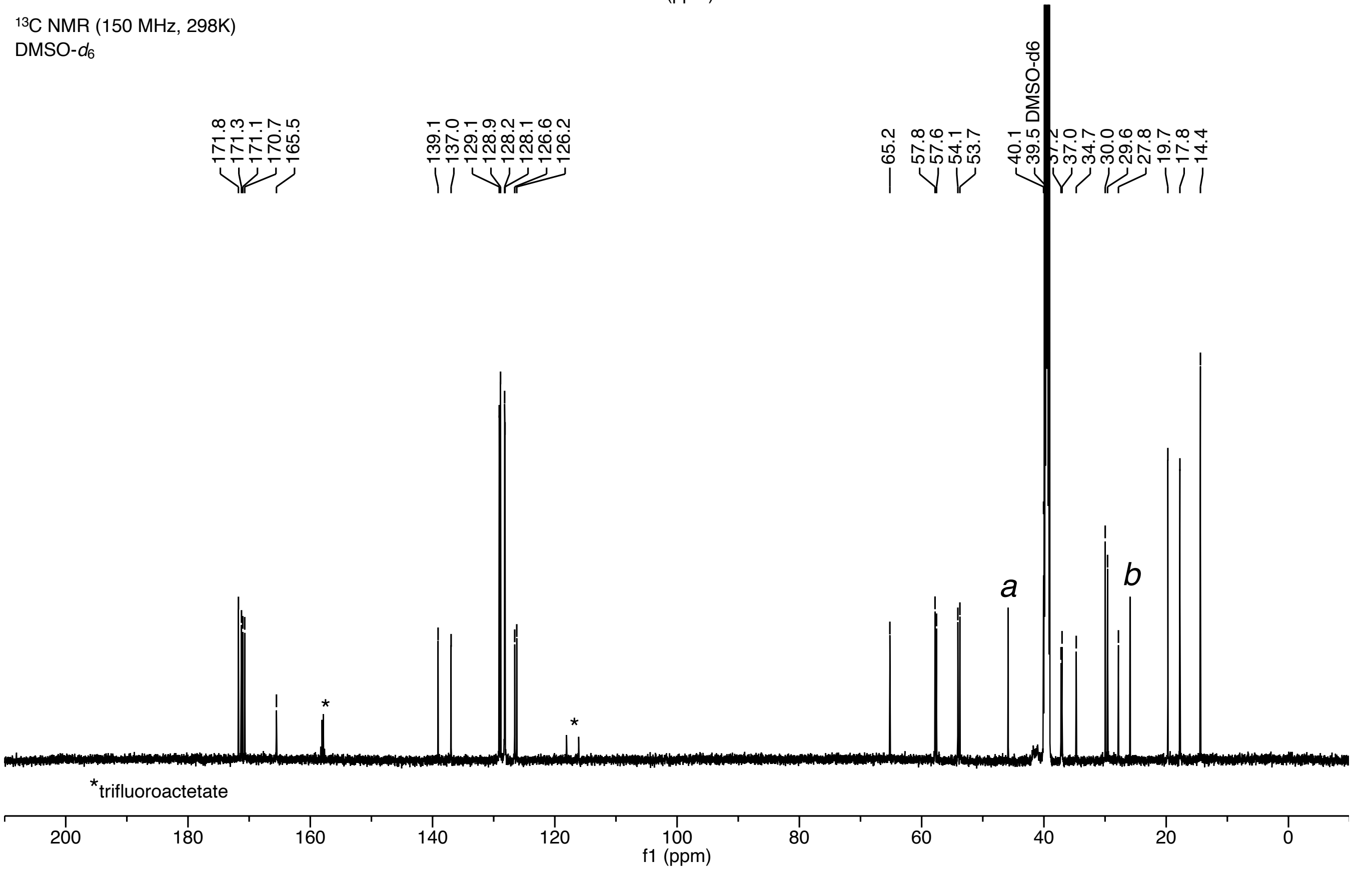

**P2 C1Dap (7)**

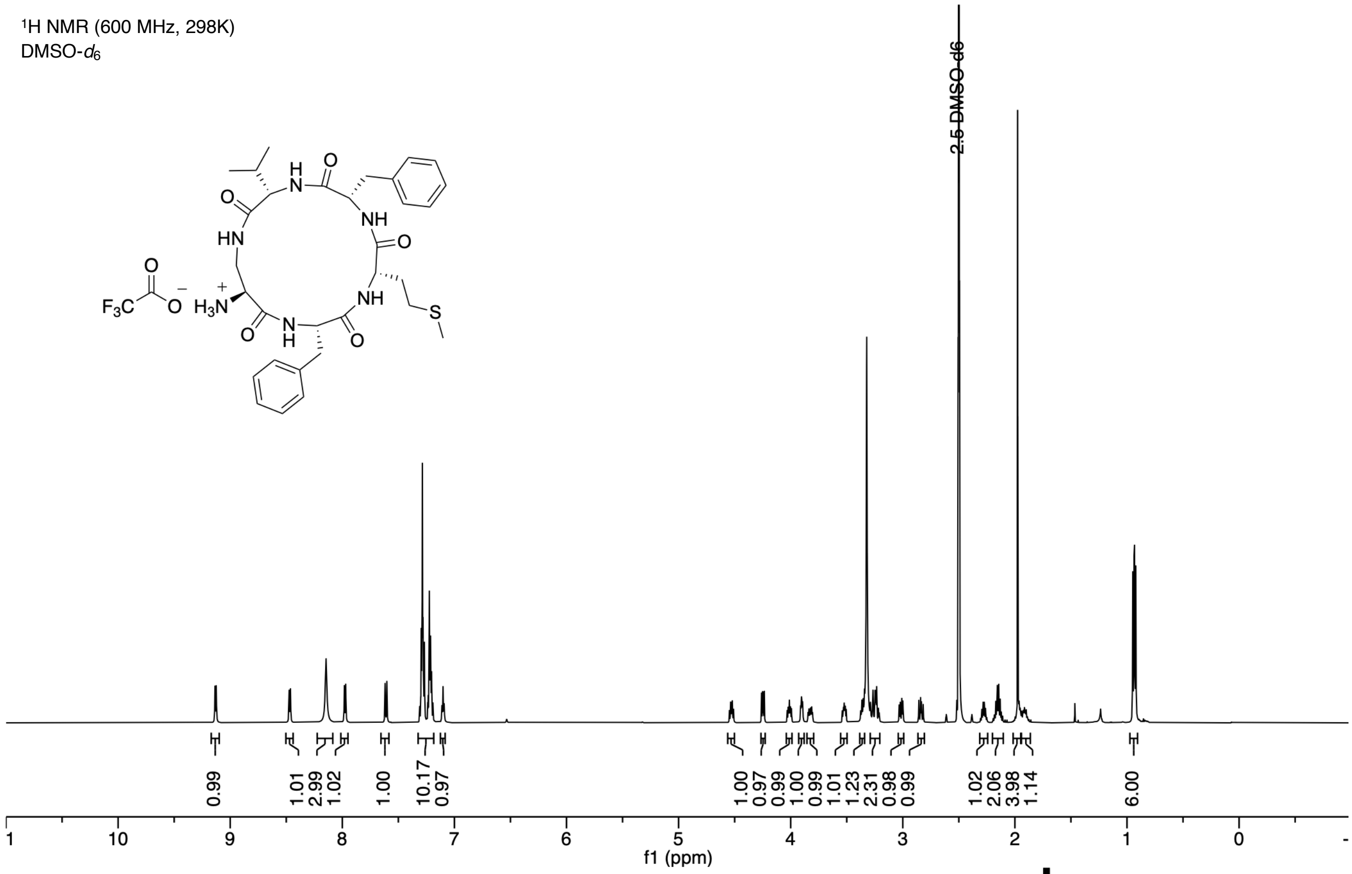
$^{1}$H NMR (600 MHz, 298K)
DMSO-$d_6$
2.5 DMSO-d6
0.99
1.01
2.99
1.02
1.00
10.17
0.97
1.00
0.97
0.99
1.00
0.99
1.01
1.23
2.31
0.98
0.99
1.02
2.06
3.98
1.14
6.00
f1 (ppm)

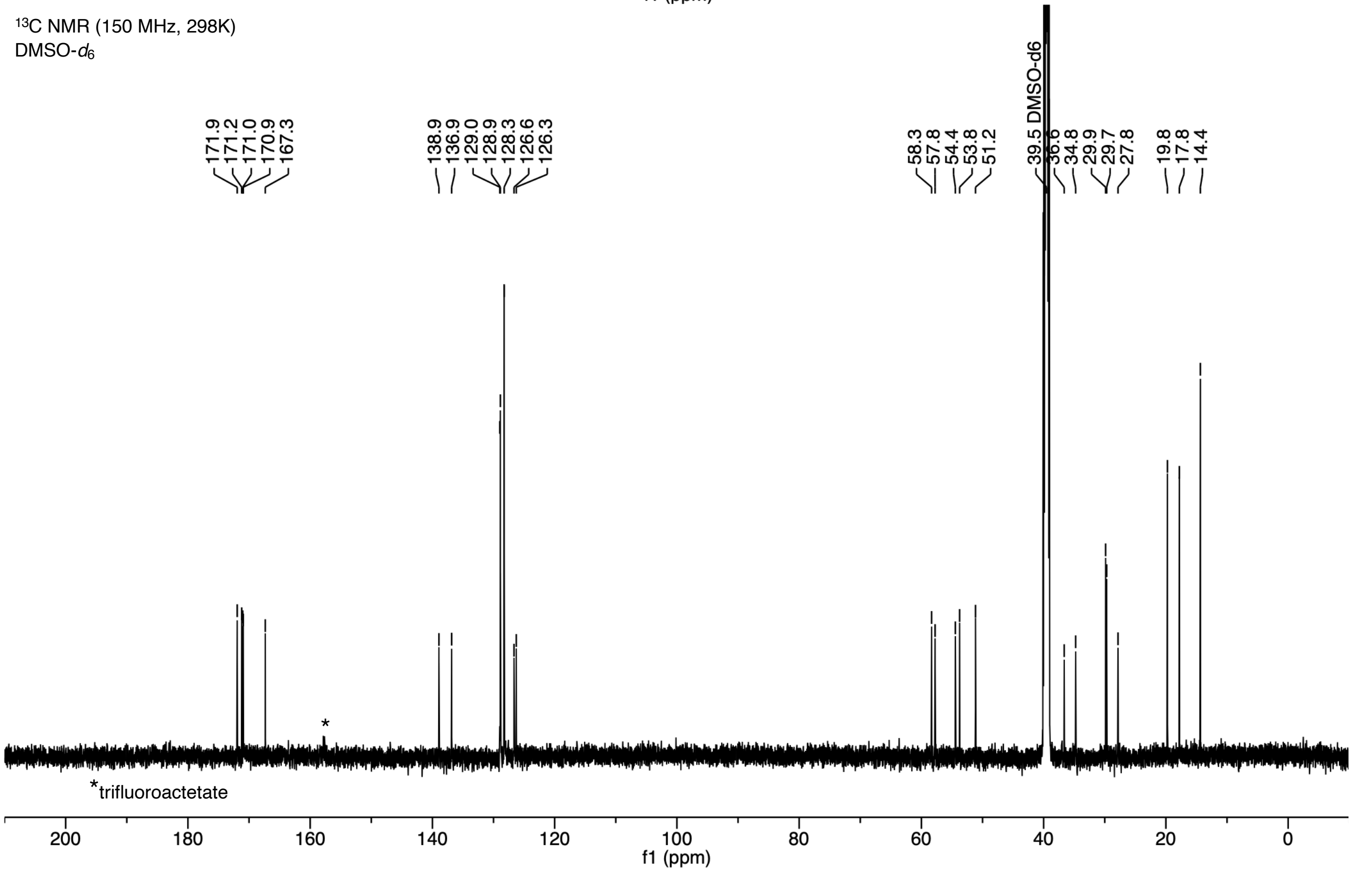
$^{13}$C NMR (150 MHz, 298K)
DMSO-$d_6$
171.9
171.2
171.0
170.9
167.3
138.9
136.9
129.0
128.9
128.3
126.6
126.3
58.3
57.8
54.4
53.8
51.2
39.5 DMSO-d6
36.6
34.8
29.9
29.7
27.8
19.8
17.8
14.4
*trifluoroactetate
f1 (ppm)

#### P2 C1Ser (8)

[1]H NMR (600 MHz, 298K)
DMSO-$d_6$

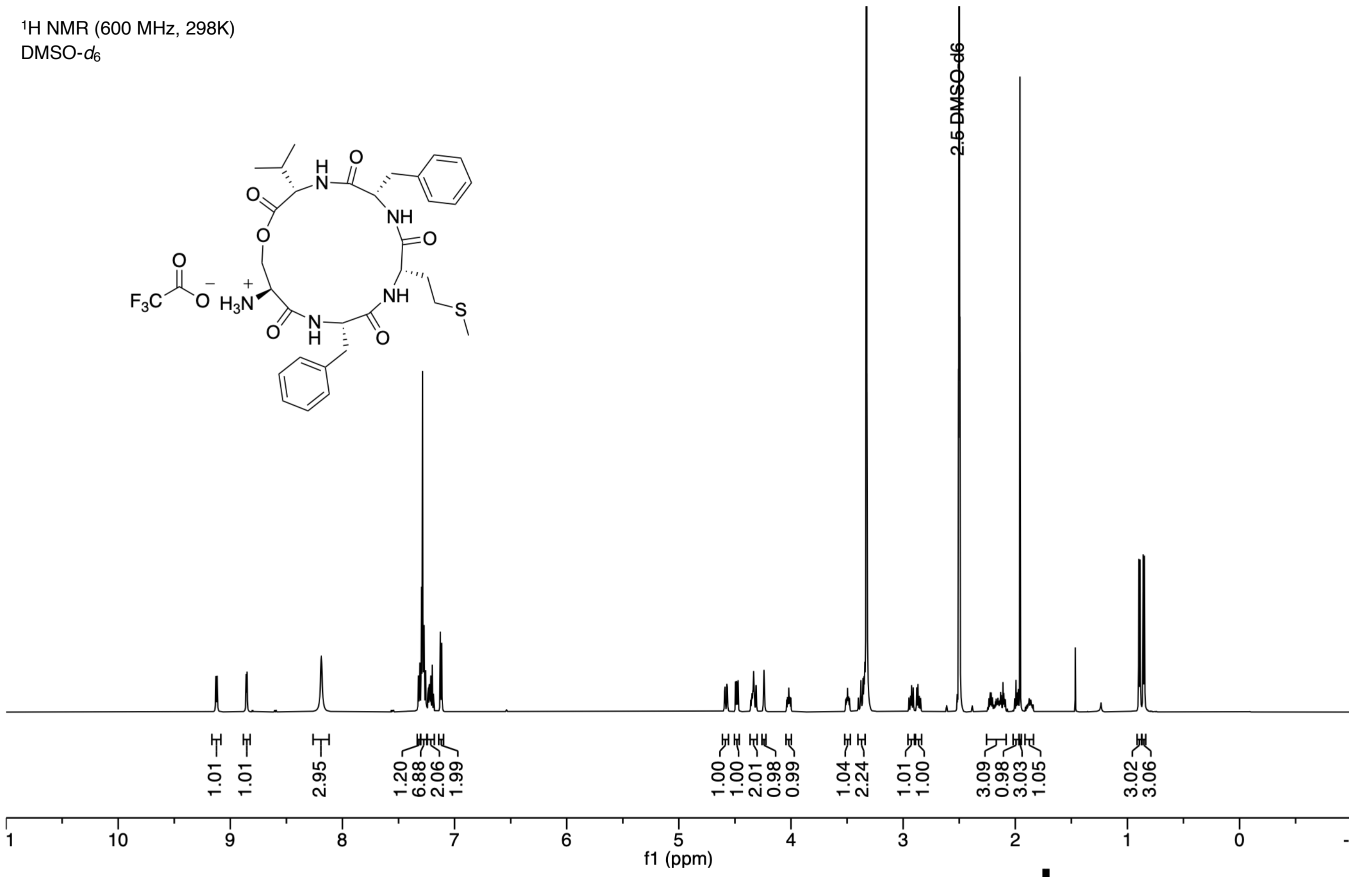

[13]C NMR (150 MHz, 298K)
DMSO-$d_6$

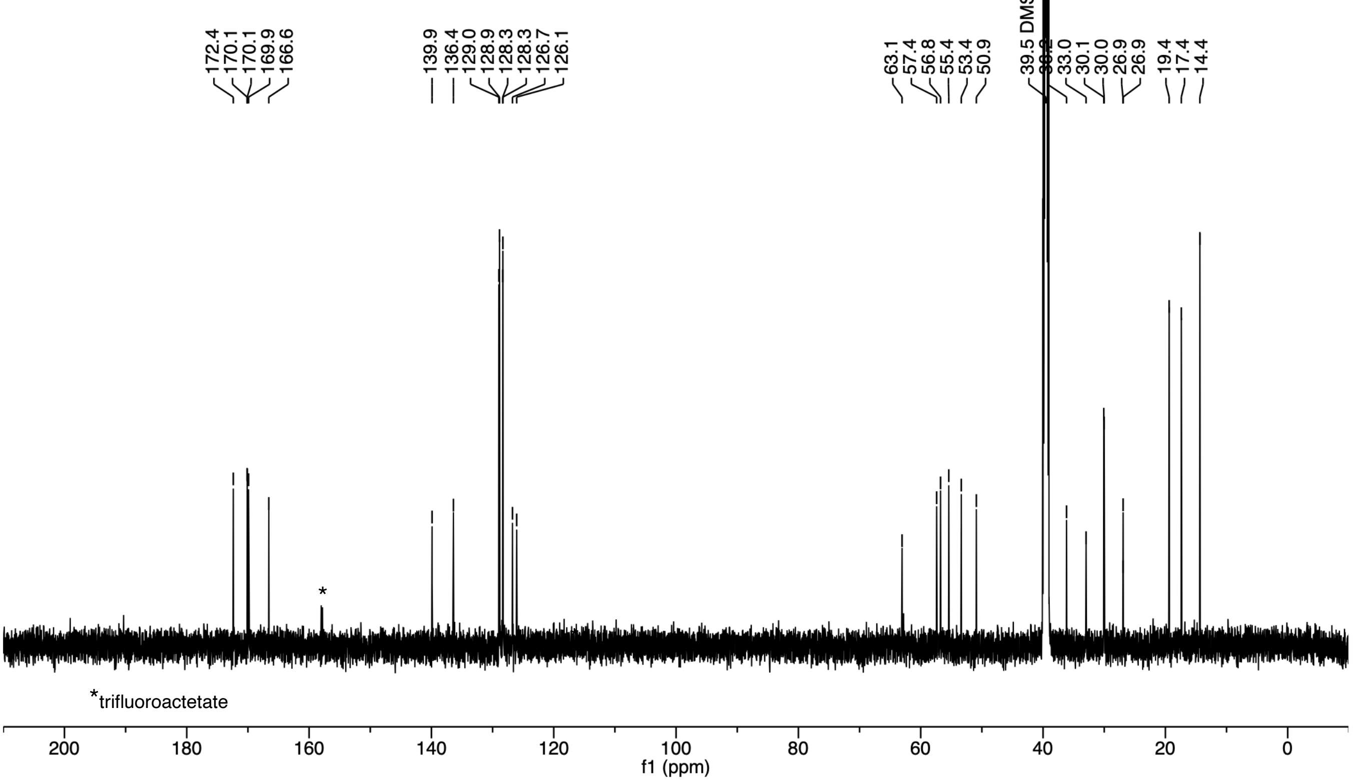

#### *N*,*N*-Dimethyl-P2 thioether (9)

$^{1}$H NMR (600 MHz, 298K)
DMSO-$d_6$

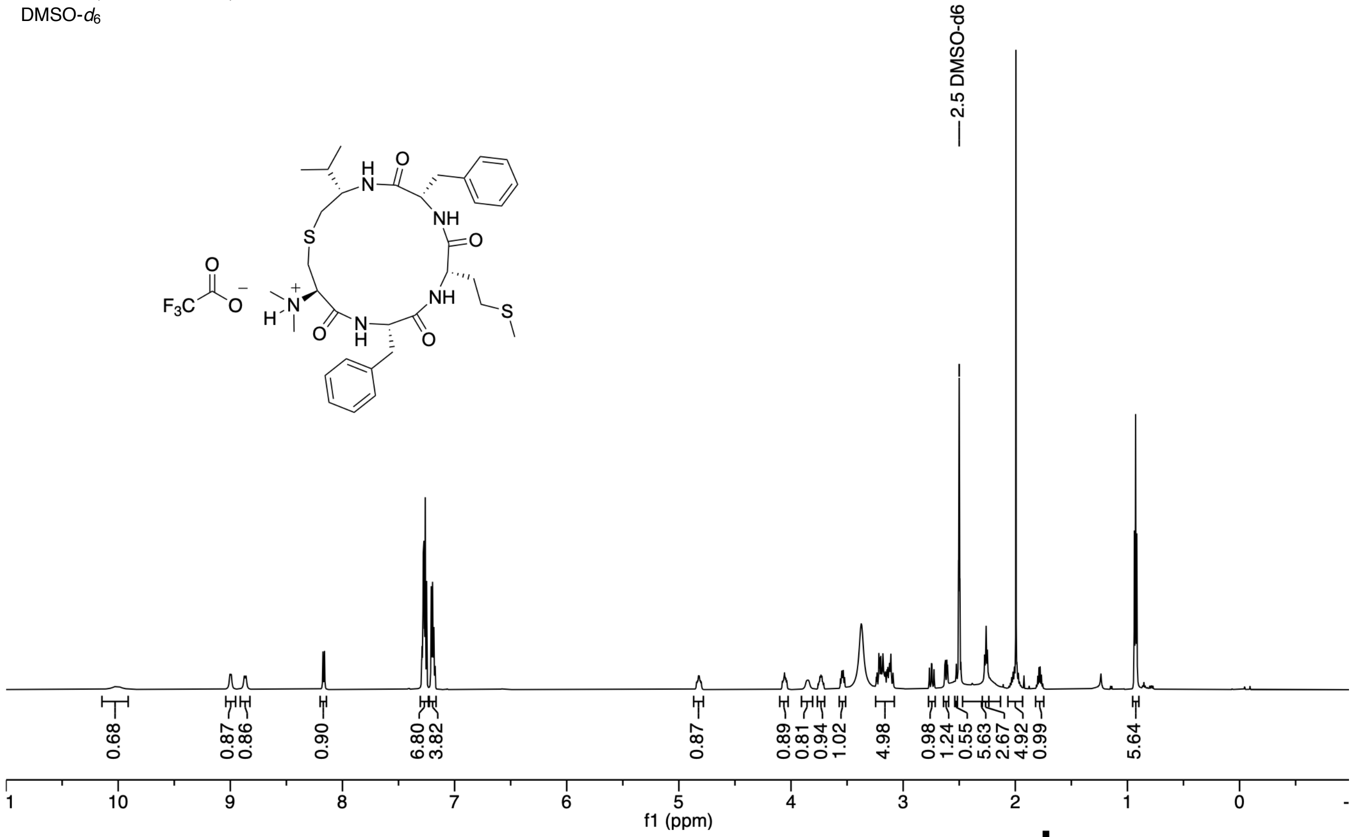

$^{13}$C NMR (150 MHz, 298K)
DMSO-$d_6$

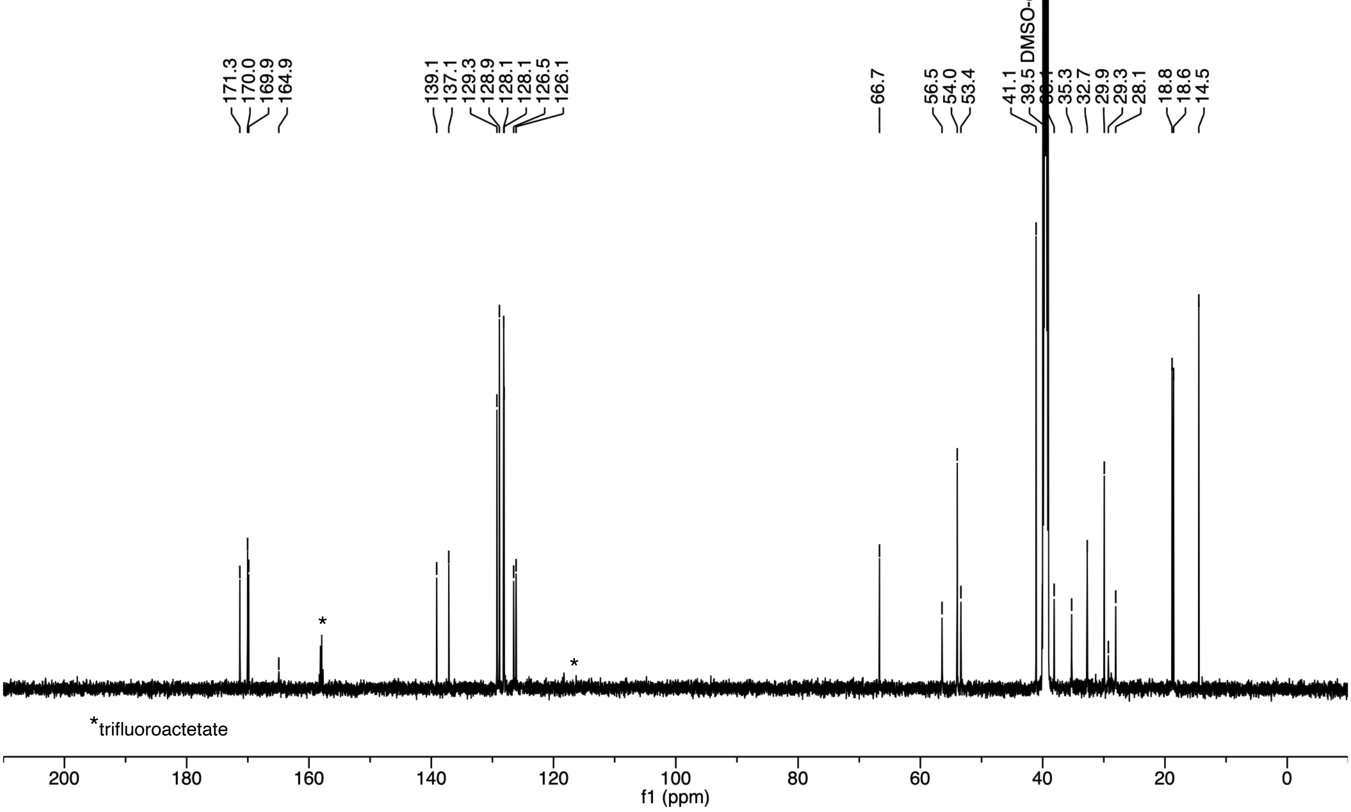

**P2 thioether (10)**

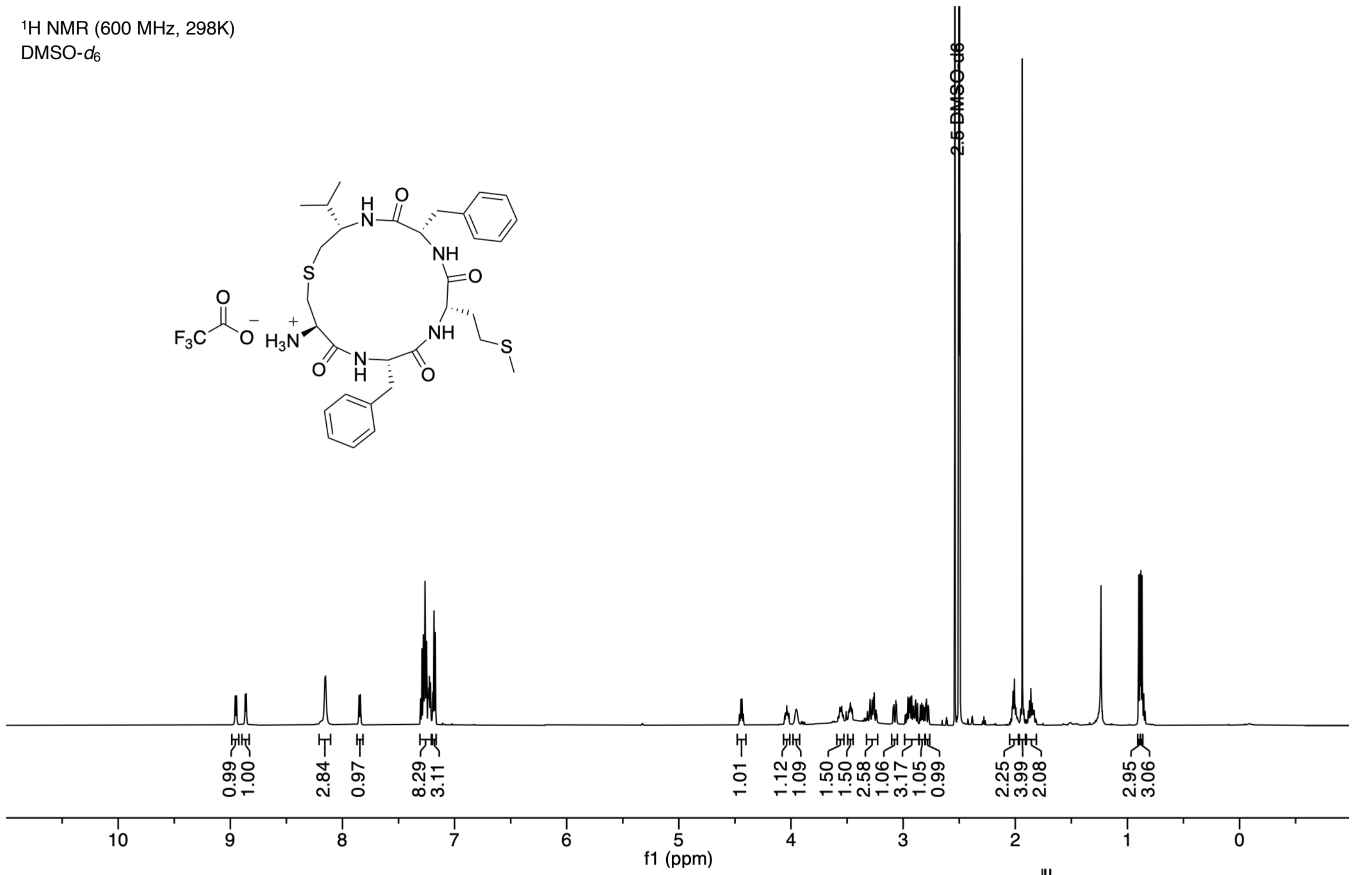

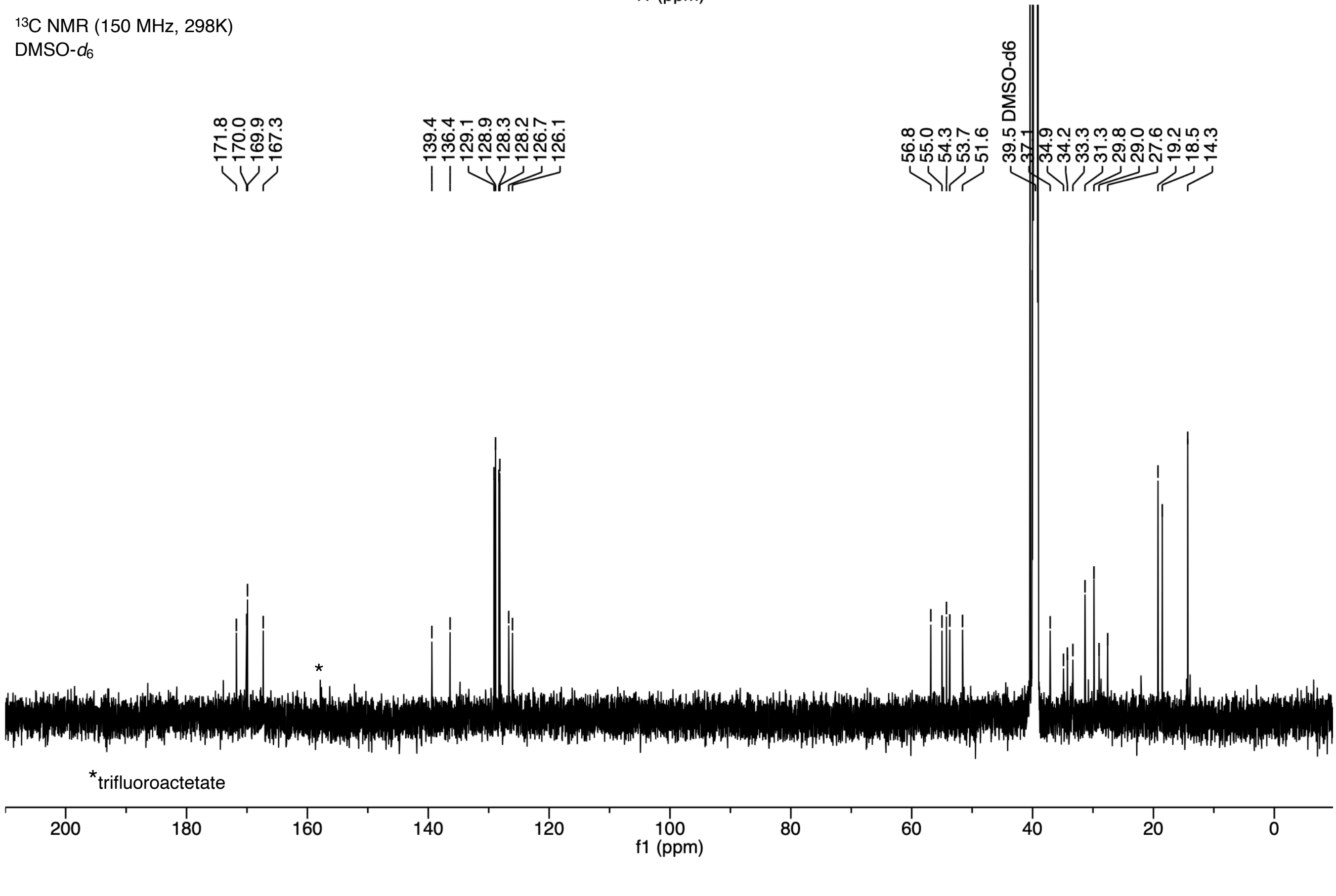
